# Population-scale variability at short tandem repeat loci reveals pathogenicity signature

**DOI:** 10.1101/2025.01.06.631535

**Authors:** Matt C. Danzi, Isaac R. L. Xu, Sarah Fazal, Egor Dolzhenko, David Pellerin, Ben Weisburd, Liedewei Van de Vondel, Chloe Reuter, Jacinda B. Sampson, Chiara Folland, Carolin K. Scriba, Gavin Monahan, Phillipa J. Lamont, Julie Wertz, Adriana Rebelo, Sophia B. Gibson, Daniel G. Calame, Haloom Rafehi, Penny Snell, Kate Kotschet, Kayli C. Davies, Igor Stevanovski, Ira W. Deveson, Danny E. Miller, Chia-Lin Wei, Jane Grimwood, Donna M. Muzny, Niall Lennon, Melanie Bahlo, Paul J. Lockhart, Matthew Wheeler, Anne O’Donnell-Luria, Stefan Wuchty, Gianina Ravenscroft, Michael A. Eberle, Kiran V. Garimella, Fritz J. Sedlazeck, Michael E. Talkowski, Michael C. Schatz, Evan E. Eichler, All of Us Research Program Long Read Working Group, Stephan Zuchner

**Author notes:** **Corresponding authors** Matt C. Danzi, PhD and Stephan Zuchner, MD, PhD. Contributed equally (Matt C. Danzi, Isaac R. L. Xu). Address: Dr. John T. Macdonald Foundation Department of Human Genetics and John P. Hussman Institute for Human Genomics, University of Miami Miller School of Medicine, 1501 NW 10^th^ Avenue, Miami, FL, 33136, USA.

## Abstract

Short tandem repeat (STR) instability is a major cause of disease association, yet their full length-distributions and sequence content have been difficult to resolve with short-read sequencing. By analyzing long-read sequencing genomes from 2,645 participants of the *All of Us* Research Program, we establish a comprehensive STR resource of 23 billion alleles covering a catalog of 4.4 million loci. We find that known disease-associated STR loci exhibit markedly elevated allelic variability relative to genome-wide distributions, which is most pronounced when based on the length of uninterrupted, pure repeat motifs (Odds Ratio: 244, p=1.62e-29). We establish the Pure Length Variability Index (PLVI) as a measure of this instability, which is robust across ancestries and replicated in 500 samples from the 1000 Genomes Project (Odds Ratio: 84, p=2.72e-28). To illustrate, we identify 55 unstable coding CAG and 5’-UTR CGG STR loci with this metric, 60% of which have not been associated with disease. From this set, we document potential pathogenic repeat expansions for *FAM193B* and *EP400* by integrating PLVI with single-genome length outlier analysis. These data establish a population-scale framework for STR discovery in rare and common diseases using long-read sequencing cohorts.

## Introduction

The genetic basis of many human diseases has been elucidated through large-scale studies of sequence variation. However, the contribution of short tandem repeats (STRs) to disease remains incompletely defined. Over 60 diseases have been attributed to expansions of STRs, with most discovered in the past ten years aided by recent technological advances^1,2^. Genetic studies into STRs have been hampered by technical challenges in accurately genotyping STRs at scale, the lack of population-scale data where the loci have been completely sequence-resolved, and a limited understanding of indicators for STR locus pathogenicity. For non-STR variation classes that alter protein sequences, several population-genetic markers of potential pathogenicity have been defined, such as minor allele frequency, constraint, evolutionary conservation, and computational modeling. This has enabled the identification of thousands of disease-causing genes containing strong-effect alleles, sometimes in only few unrelated genomes, in rare diseases, and subsets of common phenotypes. The strongest biological effects are usually linked to rare genetic changes, the by far largest group within the spectrum of population allele frequencies. Correspondingly, in nearly all repeat expansion diseases (REDs), most pathogenic alleles are exceptionally rare occurrences of outsized repeat unit counts or overall length of the repeat. Decades of in-depth, locus-specific RED studies have also shown the important effects of repeat interruptions on phenotypic expression as well as intergenerational and somatic STR stability^3^. Complex motif compositions, frequent association with Alu elements, and read-length limits of short-read sequencing have thus far proven formidable barriers to large-scale STR exploration in disease. Thus, a high-fidelity allelic catalogue of STR loci derived from long-read sequencing (LRS), analogous to gnomAD, will enable superior interpretation of individual genomes^4^. Such a dataset will also enable the systematic exploration of more general indicators of locus pathogenicity, repeat stability, and the role of genic positioning of STRs. This study of 2,645 HiFi and 500 Oxford Nanopore Technologies (ONT) LRS genomes from multiple ancestries, applying a 4.4 million loci STR catalogue^5^, has generated an unparalleled resource of STR alleles. The distributions in length and composition of individual STR loci allow for efficient genome-wide outlier analyses in individual genomes and correlations of phenotypes to STR genotypes. We show at a genome-wide level that repeat length variability, particularly when defined by the longest pure segment (LPS), emerges as the strongest predictor of STR pathogenicity. All known pathogenic STR loci, except poly-alanine loci, fell above the 95th percentile of most variable STRs. In other words, a population derived cohort, without explicit patient enrichment and lacking the observation of pathogenic STR outliers, allows for the prediction of disease-associated STR loci. This resource provides a powerful framework linking STRs to disease, which combines the Pure Length Variability Index (PLVI), genome-wide STR outlier analysis in individual genomes, and additional orthogonal data. This LRS resource will enable reverse genetic screens, where a set of high PLVI loci require genetic association to phenotypes; ideal for studies of large biobank collections with LRS genomes and broad phenotypic information. The complete data resource can be accessed through the *All of Us* Researcher Workbench, with PLVI scores and other summary statistics also available through Zenodo^6^ and TR-Explorer^5^.

## Results

### A long-read, sequence-resolved STR resource

The *All of Us* Program Long Read Working Group performed HiFi genome sequencing on 2,645 multi-ancestral individuals (v8 data release). Approximately 85% of these individuals have electronic health records (EHR) available in the AOU Researcher Workbench^7^. This dataset allowed us to perform the most detailed, long read-based, multi-ancestral STR allele analysis to date – a powerful baseline for future genotype-phenotype comparison studies. We applied the TRGT^8^ software, configured with the TR-Explorer catalog of over 4.4 million STR loci^5^, to these 2,645 HiFi genomes to produce a sequence-resolved call set of 23,310,379,710 STR alleles. STR length remains the defining metric for REDs. We observed that STR length variation over all loci spans nearly four orders of magnitude with nearly every locus displaying length variability (4,436,423 / 4,439,672; 99.93%) (Supplementary Figure 1A). In order to measure robustness to sequence read coverage and ancestry, we split the 2,645 genomes into two cohorts: a set of 543 high-coverage (all >=25x sequence read coverage; median coverage 32x) samples to serve as the gold standard and “discovery cohort” and a larger set of 2,102 low-coverage samples (all <25x; median coverage 12x) to serve as the “validation cohort” (Figure 1A). All coverage and ancestral subgroups showed Pearson correlation coefficients with the discovery cohort above 0.9 for the 0^th^ through 99^th^ allele length percentiles (Figure 1B). Thus, even relatively low-coverage genomes will generate reliable results for our subsequent analyses. Throughout the manuscript, the validation cohort will be presented both as a unified set and split into individual ancestry groups for specific analyses. We use the following six ancestry groups based on the labels used in gnomAD^4^ and the ancestry predictions made on the *All of Us* short-read cohort^9^: African (AFR), Admixed American (AMR), East Asian (EAS), European (EUR), Middle Eastern (MID), and South Asian (SAS). However, the MID ancestry group was omitted from population-specific analyses due to the low number of individuals from that group in the validation cohort. We also collected a multi-ancestral “replication cohort” of 500 ONT samples from the 1000 Genomes Project Long-read Sequencing Consortium sequenced to median coverage of 35x to demonstrate cross-technology replication of major findings^10^. The ONT replication cohort shows less agreement at lengths below the 50^th^ percentile but maintains strong agreement with the discovery cohort at the upper quantiles (Figure 1B). Combined, both the HiFi and ONT datasets provide population-scale access to 3,145 genomes or 6,290 alleles per locus.

**Figure 1:**
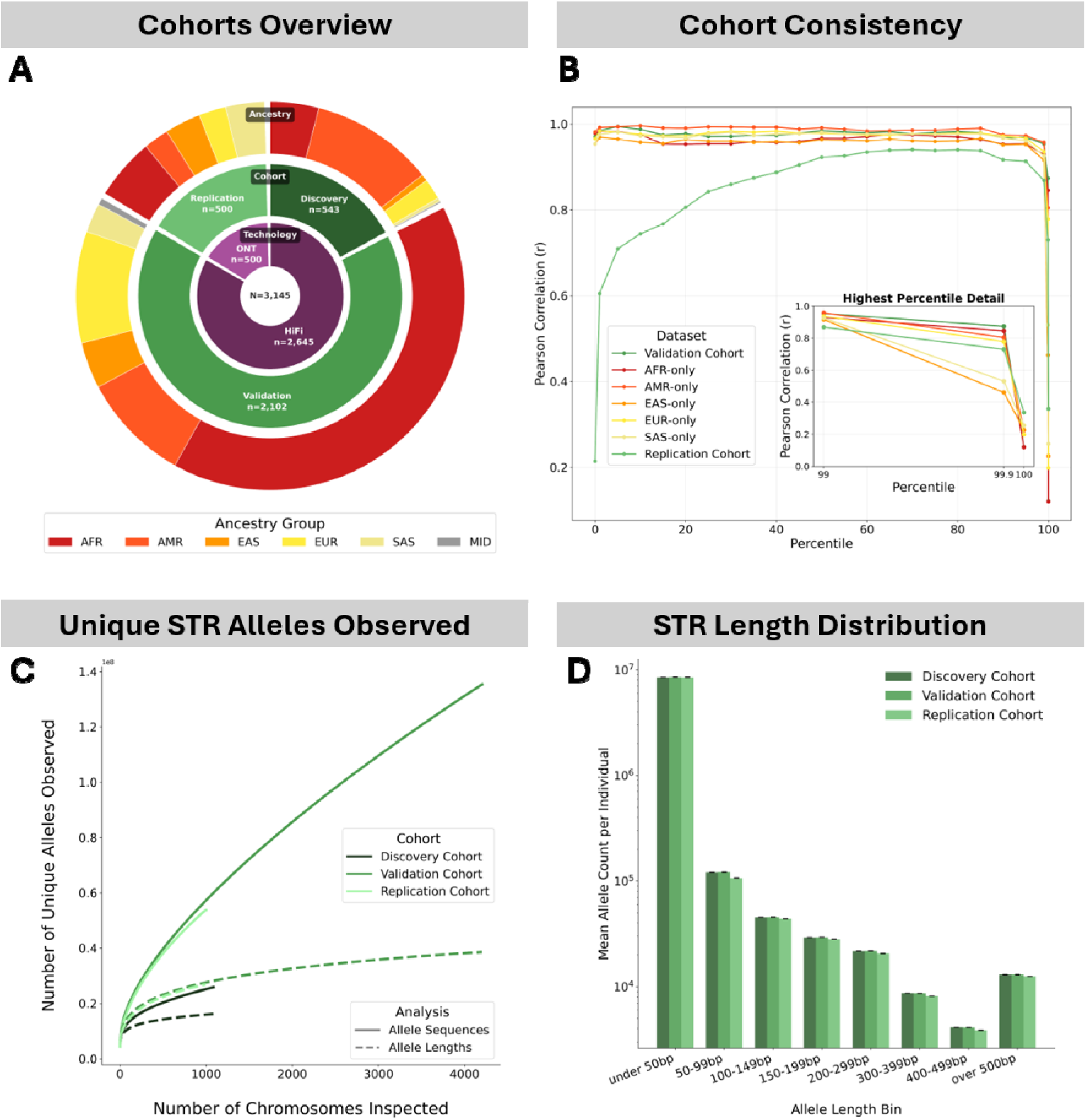
Overview of the cohorts. A) Sunburst plot depicting the samples used in this study broken down by sequencing technology, cohort, and ancestry group. B) Line plot of Pearson correlation coefficient between the length percentiles observed in the validation and replication cohorts (and each ancestry group within the validation cohort individually) against the discovery cohort. Inset: zoomed-in view of the 99^th^, 99.9^th^ and 100^th^ percentiles. C) As individuals are added to either the discovery, validation, or replication cohorts, unique new alleles are observed for both, basepair resolved variations of ‘allele sequences’ (solid lines) as well as different ‘allele lengths’ (dashed lines). D) Bar plot of the mean number of alleles of various lengths, present in each individual in each cohort. Error bars represent standard error of the mean.

Long-read datasets of STRs provide access to two historically difficult domains of STR analysis: sequence content of alleles, and better examination of alleles longer than 150bp. First, while STRs are known for their exceptionally high copy number variation, the nucleotide-level resolution of this dataset revealed that sequence variation greatly exceeded length variation in these cohorts (Figure 1C). We notice that novel allele sequence observations continue to grow linearly as additional genomes are added, albeit at a lower rate for the discovery cohort than the validation or replication cohorts (Figure 1C). Both the ONT and lower coverage HiFi datasets likely harbor more sequence error than the high coverage HiFi dataset, which inflates the number of unique allele lengths and sequences observed in those datasets. Second, while most STRs in an individual are shorter than 100bp, we identify 45,000 STRs longer than 200bp per individual, which would be difficult to evaluate accurately solely with srWGS approaches (Figure 1D).

### STR sequence composition variation

To analyze this sequence variation identified by our long-read approach, we parsed alleles into segments of repeating motifs. This revealed compositional differences beyond simple short interruptions including complex non-reference motif patterns. We further observed that these non-reference motif patterns often occur in multiple individuals and demonstrate length polymorphism of their own. This is illustrated for the *FGF14 GAA* repeat locus causing the common ataxia SCA27B^11^ (Figure 2A). Thus, for more consistent comparison of ‘intra-locus’ motifs, we calculated the lengths of the ‘longest pure (uninterrupted) segment’ (LPS) with its corresponding motif for each of the 23 billion alleles. This reduces the allelic complexity for each STR locus (Figure 2A, B) and emphasizes the length distributions of the different LPS motifs observed in the discovery cohort (Figure 2C). This representation uncovers how the different LPS motifs show unique length distributions, with the SCA27B-causing AAG motif exhibiting the smallest mean LPS size in the general population. Scaling this analysis to all 4.4 million loci in the discovery cohort revealed abundant motif heterogeneity, with ∼55 novel LPS motifs being observed in each additional genome studied (Figure 2D). This is reminiscent of early cohorts of whole exome studies that yielded an unexpected excess of rare variation^12^.

**Figure 2:**
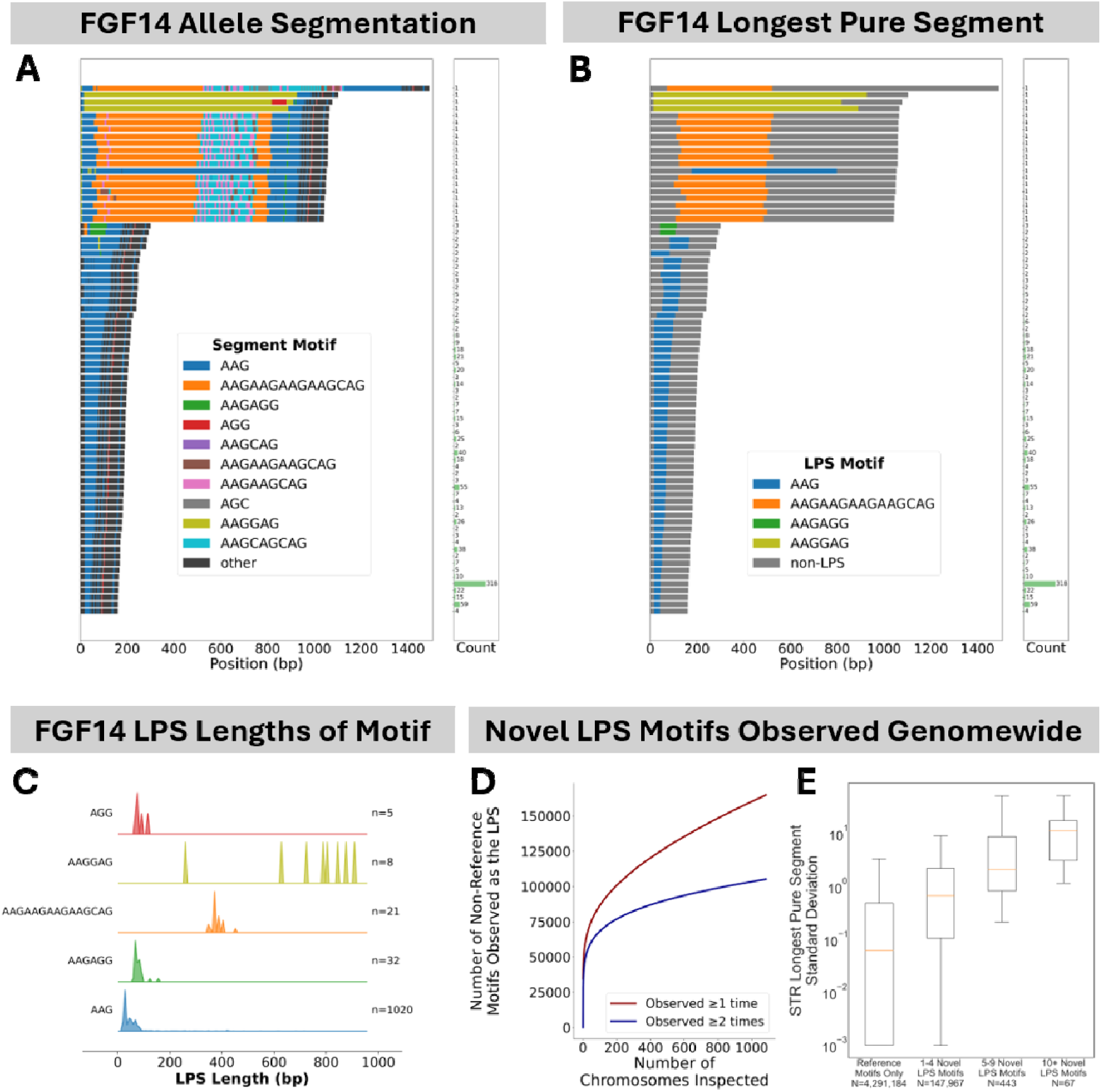
Sequence-resolved STR variation in 3,145 genomes. A) Waterfall plot showing allele segmentation at the SCA27B locus in the discovery cohort. B) The same waterfall plot with colors now only showing the longest pure segment (LPS) and all other segments set to grey. C) Ridge plot of the lengths of the longest pure segment for the SCA27B locus in the discovery cohort stratified by LPS motif. D) Saturation curve of novel LPS motif discovery showing number of previously unseen novel LPS motifs observed per individual as individuals are added to the cohort. The red curve shows the first observation of each novel LPS motif, while the blue curve tracks the second observation. E) Boxplot of standard deviation in the length of the LPS among groups of loci which only exhibit the reference motifs, 1-4 novel LPS motifs, 5-9 novel LPS motifs, and 10 or more novel LPS motifs.

Despite this compositional complexity, we found that 95.9% of STR loci exhibit only a single LPS motif in the discovery cohort. Interestingly, in 3.4% of the loci that show only a single LPS motif, it is not one currently represented in the TR-Explorer catalog. Thus, in total, 3.3% of STR loci exhibit non-reference LPS motifs. We found that these loci show greater length variation than the loci that do not exhibit LPS motif variation and this pattern increases with the number of LPS motifs (Figure 2E). We also observed a larger degree of LPS length variation in longer tandem repeat motifs (12+bp) compared to shorter motifs (Mann-Whitney U-Test; generalized odds ratio: 26.35, p-value<1e-10) consistent with their higher mutation rate^13^. Surprisingly, trinucleotide repeats show markedly less variation than any other motif length (Supplementary Figure 3A) (Mann-Whitney U-Test; minimum odds ratio of pairwise comparisons: 1.75, p-value<1e-10) even when restricting to intergenic loci (Supplementary Figure 3B) (Mann-Whitney U-Test; minimum odds ratio of pairwise comparisons: 1.78, p-value<1e-10).

### High allelic length variability identifies disease-associated STR loci

Having established the extraordinary range of allelic variability across STR loci, we next asked whether disease-associated loci occupy a distinct position within this distribution. As shown in Figure 3A, disease-associated STR loci fall overwhelmingly at the extreme tail of the allele length variability distribution, a pattern that prompted a more systematic investigation. A few prior studies using LRS approaches on small numbers of samples have noted that many disease-associated STRs are highly polymorphic in the general population^14,15^. However, the associations shown were modest and were not replicated across ancestry groups, cohorts, sequencing technologies, or tandem repeat catalogs. Crucially, neither study incorporated sequence content variation in their analyses.

**Figure 3:**
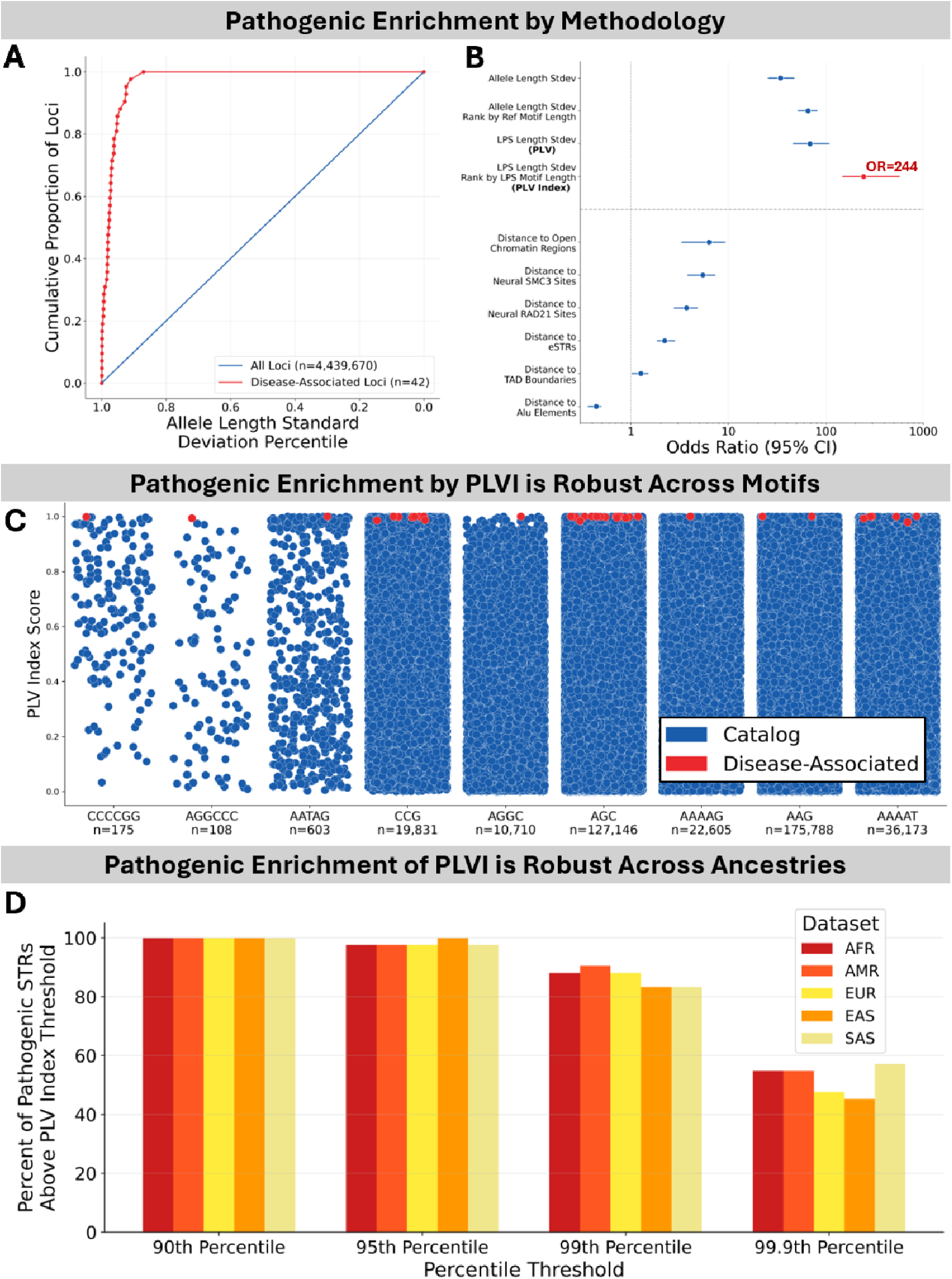
High allelic length heterogeneity identifies disease-associated STR loci. A) Cumulative proportion of catalog and disease-associated loci observed with increasing values of the percentile of the standard deviation of allele length. B) Forest plot of the generalized odds ratio (with 95% confidence interval) comparing the enrichment of disease-associated loci applying PLVI, other population-derived variation measures, and established, annotation-based measures. C) Percentile of PLVI scores at disease associated STR loci with one strip for each relevant simplified motif. For each locus, the most common motif seen at the locus is used. The height is the percentile from 0-1. Catalog points are in blue and known disease-associated loci are in red. D) Grouped bar plot of the percentage of disease-associated STRs falling above each percentile threshold (x-axis) for each individual ancestry group within the validation cohort. All groups are analyzed by PLVI.

To measure the extent of this effect in our cohorts, we focused on a set of 42 STRs whose expansion is unequivocally associated with disease^16^ (see Supplementary Data for details on selection). In the discovery cohort, the standard deviation of allele length at RED loci was significantly greater than observed across the complete STR catalog (Mann-Whitney U-Test; p<1e-10; generalized odds ratio = 34.1) (Figure 3B). Specifically, 36 of the 42 disease-associated STRs (85.7%) were in the top five percent of most variable loci genome-wide as measured by standard deviation of allele length; 12 of the 42 RED loci (28.6%) were in the top one percent (Figure 3A). Next, we applied the LPS approach to this analysis considering only the single most commonly observed motif at each locus rather than measuring overall STR allele length. This increased the enrichment (Mann-Whitney U-Test; p<1e-10; generalized odds ratio = 68.6) (Figure 3B), including placing 39 of 42 RED loci among the five percent most variable (Supplementary Figure 5A). We term this measure of the standard deviation of LPS length of an STR as the Pure Length Variability (PLV). Since STR variability is affected by motif length, we next grouped all loci by STR motif length. Within each motif length stratification, we ranked STR loci by their PLV value, yielding a metric we term the Pure Length Variability Index (PLVI). Using PLVI, we observed a remarkable 88.1% (37/42) of the RED loci among the top one percent most variable loci. More than half of the RED loci fell within the top 0.1 percent most variable loci (59.5%; 25/42) (Supplementary Figure 5A), leading to a large increase in overall levels of enrichment (Mann-Whitney U-Test; p<1e-10; generalized odds ratio = 243.9) (Figure 3B). This demonstrates that disease-associated STRs are among the most variable STR loci in their motif group in the human genome, based on a general population sample. This effect can be observed using non-sequence-resolved measures of allele length variation, but it is substantially improved by incorporating LPS length variation and LPS-based motif grouping.

Prior studies have identified numerous other factors that enrich for disease-associated STRs^17,18^, including proximity to TAD boundaries, open chromatin regions, and neural cohesin binding sites. We tested each of these in our dataset alongside PLVI and the other population-derived variation measures introduced above (Figure 3B). All previously identified annotation features replicated as statistically significant. However, none came close to the effect size of PLVI, and the population-derived variation measures as a group substantially outperformed all genomic annotation features. This demonstrates that locus variability in the general population is a far more powerful predictor of disease association than genomic location. The strength of this signal is visually apparent in Figure 3C, where disease-associated loci cluster overwhelmingly at the ceiling of the PLVI distribution across motif groups.

We performed several replication studies of this effect. First, the validation cohort (2,102 genomes) nearly exactly matched the findings of the discovery cohort (543 genomes) (Supplementary Figure 6A). A stratification of the validation cohort into individual ancestry groups also found nearly identical behavior across each of those subsets (Figure 3D). This is especially notable for RED loci that are known be expanded into pathogenic range in founder populations yet would be highly prioritized in every studied ancestry group. For example, SCA31 is the third most common spinocerebellar ataxia in Japan but is almost entirely absent in European and North American cohorts^19^. However, this locus has a PLVI above the 99.97^th^ percentile in the AFR, AMR, EUR, and SAS populations, in addition to the EAS population in our validation cohort. This demonstrates that it is well-prioritized by this approach even in populations that do not manifest the disease. Second, we repeated this analysis in the replication cohort of ONT genomes and confirmed high consistency with the two HiFi cohorts, differing by only a few percentage points (Supplementary Figure 6A). Finally, the influence of the underlying STR catalogue on this measure was studied in the discovery and validation cohorts by applying the Adotto catalogue of 1.7 million loci created with significantly different definitions of TR regions. The observed enrichment of known RED loci into the top one percent of PLVI scores was strikingly similar (Supplementary Figure 6B). This replication across technologies, STR catalogs, and ancestry groups demonstrates the robustness of this observation as a pathogenicity signature of all but a few REDs.

### Retrospective analysis of disease-causing TR linkage regions, GWAS signals, and individuals with pathogenic expansions

For an orthogonal test of PLVI as a locus prioritization tool, we retrospectively applied it to published linkage regions of 15 diseases now known to be caused by STR expansions^11,20–38^ (Figure 4A, Supplementary Figure 8, Supplementary Table 1). In 13 of 15 cases, the disease-associated STR ranked above the 99th percentile of all loci within its linkage region by PLVI. More strikingly, in 10 of 15 cases it ranked within the top 3 STRs in the region; in some instances among tens of thousands of candidates. For example, *HTT* ranked 1st among 4,232 STR loci in the HD linkage region; *RFC1* ranked 1st among 3,778 in CANVAS; and *SAMD12* ranked 3rd among 11,130 in *FAME1*. In the largest region tested, a 63 Mb interval for NIID*, NOTCH2NLC* still ranked 15th among 61,228 loci, placing it in the 99.98th percentile. These results demonstrate that a simple population-derived polymorphism metric could have immediately directed investigators toward the correct locus in the majority of these historically protracted gene discovery efforts.

**Figure 4:**
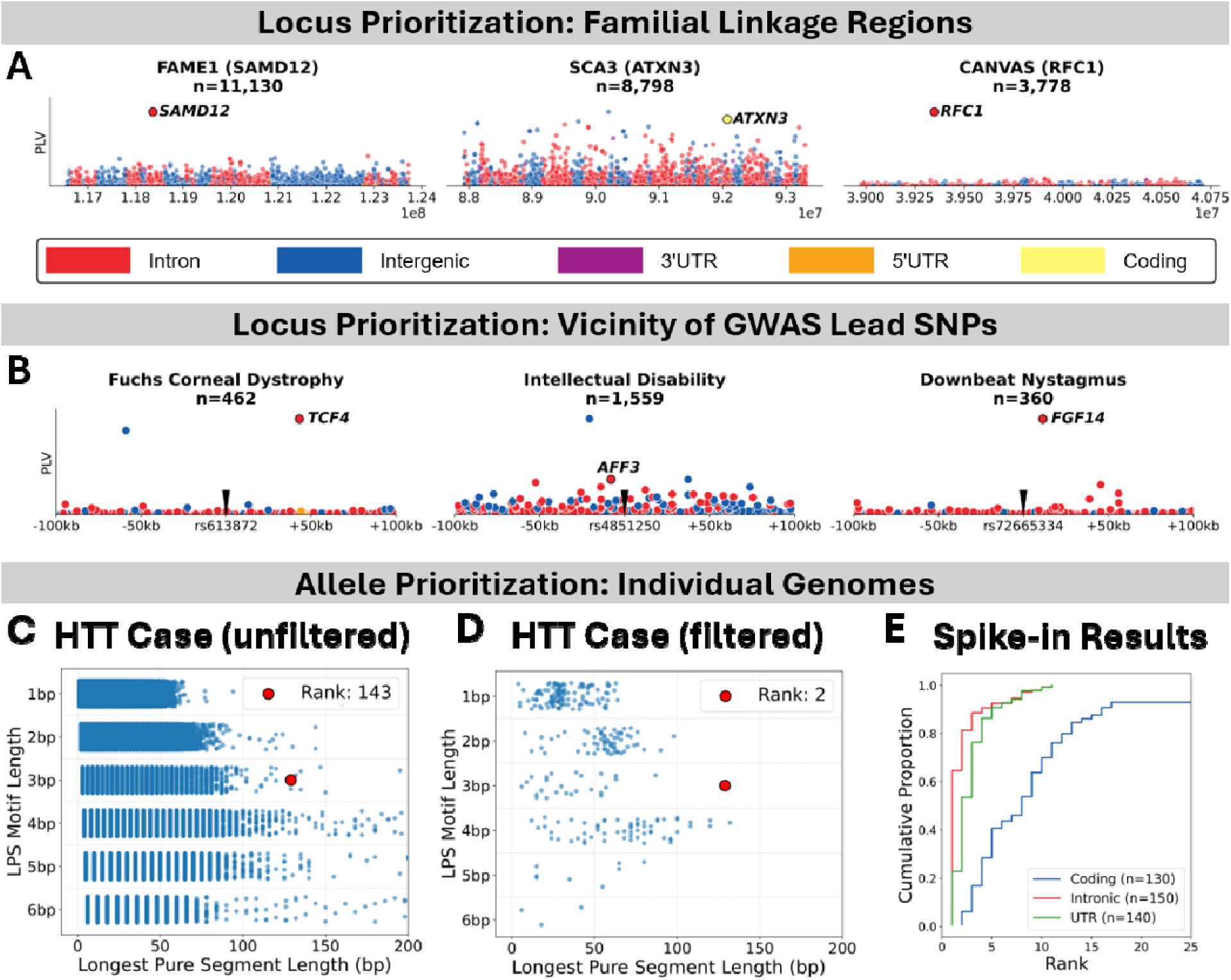
Retrospective analysis of disease-causing STR linkage regions, GWAS signals, and individuals with pathogenic expansions. A) Series of Manhattan plots of STRs across three published linkage regions. In each panel, the x-axis is the genomic coordinates across the linkage region span and the y-axis is the PLV score of each STR. Disease-associated STRs are specifically labelled in each plot. B) Manhattan plots of STRs within 100kb of each of three published GWAS SNPs (black pin), with the associated phenotype identified above each panel. In each panel, the x-axis is the genomic distance from the identified SNP, and the y-axis is the PLV score of each STR. Disease-associated STRs are specifically labelled in each plot. C-D) Swimlane plots showing the prioritization of a pathogenic-length *HTT* allele in an individual. C) No filtering for variable loci or outlier alleles. D) Loci filtered to only those above the 95^th^ percentile of PLVI scores and alleles filtered to only those longer than the 99.9^th^ percentile in the discovery cohort. E) Empirical cumulative distribution of 42 pathogenic-length STRs spiked into each of 10 HPRC samples. The 13 coding, 15 intronic, and 14 UTR STRs are plotted as separate lines.

We next examined three replicated GWAS signals whose causal variant has since been identified as a nearby STR expansion: *TCF4* for Fuchs corneal dystrophy^39^, *AFF3* for intellectual disability^40^, and *FGF14* for downbeat nystagmus^41^. Mapping all STRs within 100kb of each index SNP, the causal STR ranked first or second by PLVI in all three cases (Figure 4B). This retrospective analysis highlights an approach to further dissecting the thousands of replicated GWAS associations without current molecular understanding^42^. Notably, this prioritization was performed without ancestral matching to the original GWAS cohort.

In order to assess the usefulness of this database of allelic variation for the analysis of individuals with rare diseases, we next tested our ability to detect now-disease-associated STR expansions in individuals. First, analogous to the filtration of common SNVs from a sample, we used the discovery cohort as a collection of common variation to ask how many ‘outlier’ expansions we detect per individual in the validation cohort. Outliers were defined as repeat alleles whose length was longer than the 99.9^th^ percentile in the discovery cohort. The 99.9^th^ percentile was chosen to avoid issues related to late-onset and incomplete penetrance of diseases. Using allele length as the measure, we observed a median of 3,654 outlier tandem repeat alleles per individual in the validation cohort (Supplementary Figure 9A). Using the LPS length as the measure reduced this to 2,251 outliers per individual. Further selection to just the loci with the five percent highest PLVI scores reduces this down to a median of 155 outliers per individual (Supplementary Figure 9B). Illustrating this, Figure 4C shows a swimlane plot for an individual with a pathogenic-length *HTT* allele without allelic length outlier or locus PLVI score filtering (essentially the status quo prior to this work). This analysis would have resulted in the *HTT* expansion being ranked 143^rd^. Figure 4D shows the same individual again with the allele and locus filtering, which lifts the pathogenic allele to the 2^nd^ highest rank. Extending this approach with long-read sequencing data from five additional individuals with REDs – collected independently for this analysis and not included in the three cohort (two with SCA27B, one with SCA2, one with SCA3, and one with DRPLA) – we prioritized the disease-causing expansion with a median rank of 2 among STRs (Supplementary Figure 10A-E). All six cases had the disease-causing expansion ranked in the top 10 STRs. We further performed a larger study of this approach with 420 simulated samples, which revealed that over 90% of intronic and UTR disease-causing expansions were identified within the top 5 ranked STRs in each genome, and coding expansions were ranked within the top 16 STRs (Figure 4E, see Supplementary Results for more information).

### Prospective analysis of coding CAG loci identifies candidate ataxia STR in *EP400*

Pursuant to the prediction above that disease-causing STRs are likely to be among the most polymorphic for their motif length, we decided to interrogate the tail of the spectrum for the two most frequent causes of REDs: coding CAGs and 5’-UTR CGGs (presented in the next section). CAG repeats within coding regions are responsible for at least 12 REDs. For all 8,199 CAG repeats in coding regions, we plotted the standard deviation of the LPS length (PLV, which underpins the PLVI score) on the y-axis against the median LPS length of the locus on the x-axis (Figure 5A). This illustrates that the currently disease-associated coding CAGs are indeed some of the most extreme outliers of the entire set. There is also a group of 13 additional loci without currently known phenotype association, which show similar properties (LPS standard deviations of at least 8 and median LPS lengths of at least 35bp). The 95^th^ percentile of PLV is plotted as a green dashed line to show that all marked loci lie above the recommended PLVI threshold. Of those 13 loci, 12 were reproduced when analyzing the validation cohort with the same thresholds (highlighted in green). We predict that these 12 loci represent very strong candidates for novel REDs.

**Figure 5:**
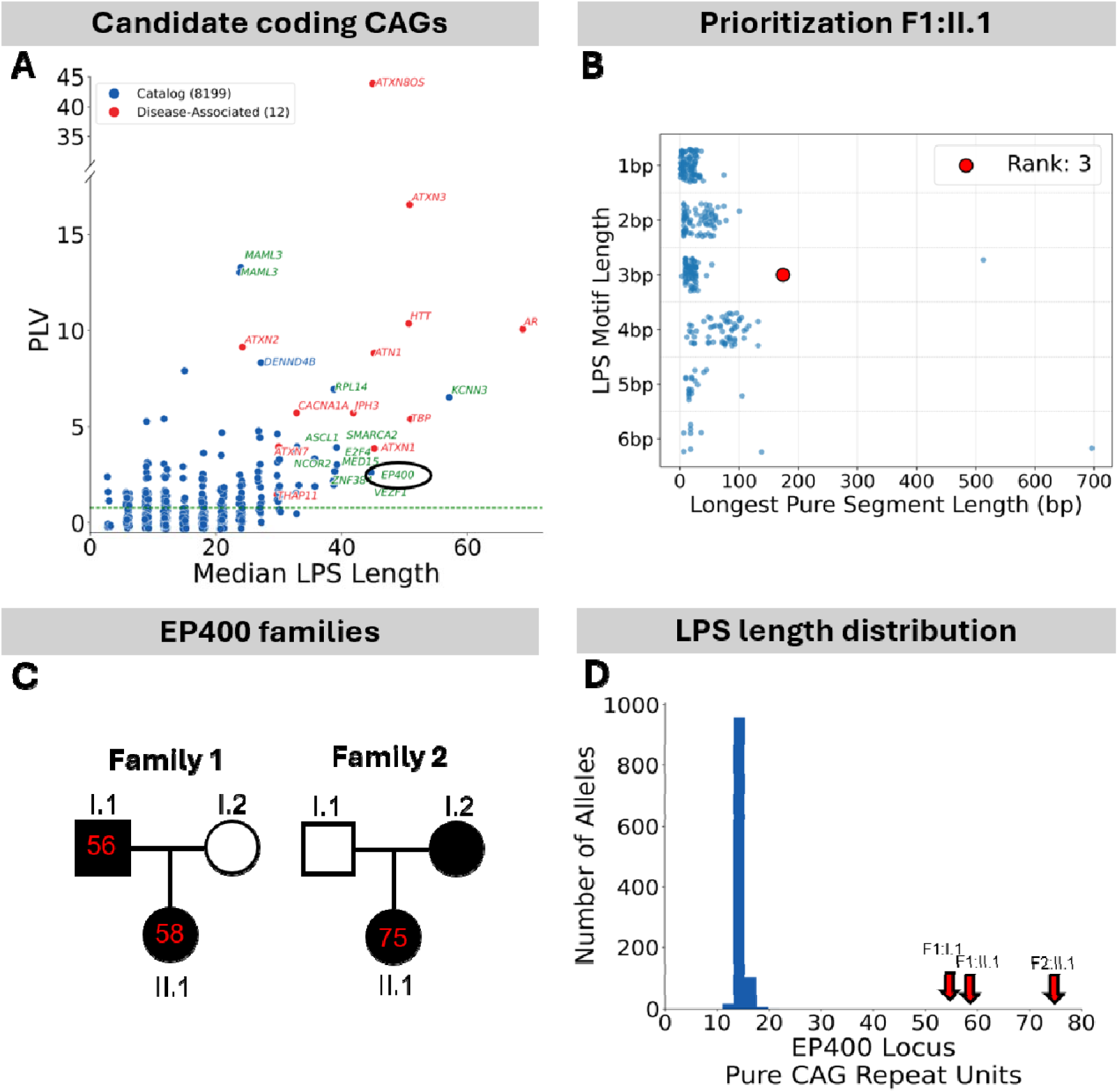
Investigation of coding CAG repeats in *EP400* as a candidate for spinocerebellar ataxia. A) Scatterplot of the coding CAG repeats in the discovery cohort plotting the PLV score (y-axis) against the median LPS length (x-axis). Disease-associated STRs have their gene names written in red. Other seeming outlier loci have their gene names written in blue (if they did not replicate in the validation cohort) or green (if they replicated in the validation cohort). B) Swimlane plot showing the prioritization of the *EP400* STR in the proband of Family 1 by combining PLVI and 99.9 percentile STR outliers. C) Pedigree of the two identified families with spinocerebellar ataxia. Number of CAG repeats measured at the *EP400* locus is given for each measured family member. D) Histogram of LPS length for the CAG motif at the *EP400* locus in the discovery cohort. Red arrows mark the allele sizes observed in the three affected individuals who were not part of this cohort.

Analysis of approximately 100 existing LRS samples of ataxia patients identified expansions of a coding CAG repeat in *EP400* (circled in Figure 5A) as the third largest candidate allele in one affected individual (Figure 5B). In that family (Family 1), the father and daughter both presented with spinocerebellar ataxia at ages 42 and 43, respectively (see Supplementary Results for complete details), and showed pure tracts of 56 CAG repeats in the father and 58 in the daughter (Figure 5C). We additionally identified Family 2, in which a mother and son both presented with spinocerebellar ataxia at ages ∼35 and 15, respectively, and the son carried a pure tract of 75 CAG repeats at the *EP400* locus (the mother is deceased and material is not available for this study) (Figure 5C). Each of these expansions represented large outliers from the general population distribution, where the longest pure CAG segment at this locus was 24 repeats (Figure 5D). The complete allele sequences in this cohort are visualized in Supplementary Figure 15A. We further genotyped the *EP400* repeat in the 243,043 individuals from the *All of Us* Research Project’s v7 short-read genome cohort (Supplementary Figure 15B which revealed a population distribution of repeat lengths consistent with that observed in the smaller long-read cohort. *EP400* is highly constrained for SNV variation (pLI=1, LOEUF=0.35 in gnomAD v4.1.1^4^), *de novo* deleterious variants have been associated with neurodevelopmental disorders^43^, and the protein is highly expressed in the cerebellum. The pattern of expression is reminiscent of several other genes that cause spinocerebellar ataxia through poly-glutamine expansions, including *ATXN2*, *ATXN3*, and *ATXN7* (Supplementary Figure 16). This poly-glutamine repeat was also predicted to be pathogenic by the RExPRT tandem repeat pathogenicity prediction AI tool with a perfect score of 1.0^44^. While preliminary, we believe this expansion in *EP400* is a strong candidate for a novel spinocerebellar ataxia.

### Prospective analysis of 5’-UTR CGG loci identifies candidate OPDM STR in *FAM193B*

Unstable 5’-UTR CGG repeats are responsible for at least 10 REDs. For all 6,416 CGG repeats in 5’-UTRs, we plotted the standard deviation of the LPS length (PLV) on the y-axis against the median LPS length of the locus on the x-axis (Figure 6A). The 95^th^ percentile of PLV (green dashed line Figure 6A) confirms that the currently disease-associated 5’-UTR CGGs represent extreme outliers. We identify a set of 23 unstable CGG repeat loci (i.e. LPS standard deviations of at least 8 and median LPS lengths of at least 45bp) currently unrelated to disease. 21/23 of these loci were confirmed when analyzing the validation cohort with the same thresholds and, thus, represent very strong candidates for novel REDs.

**Figure 6:**
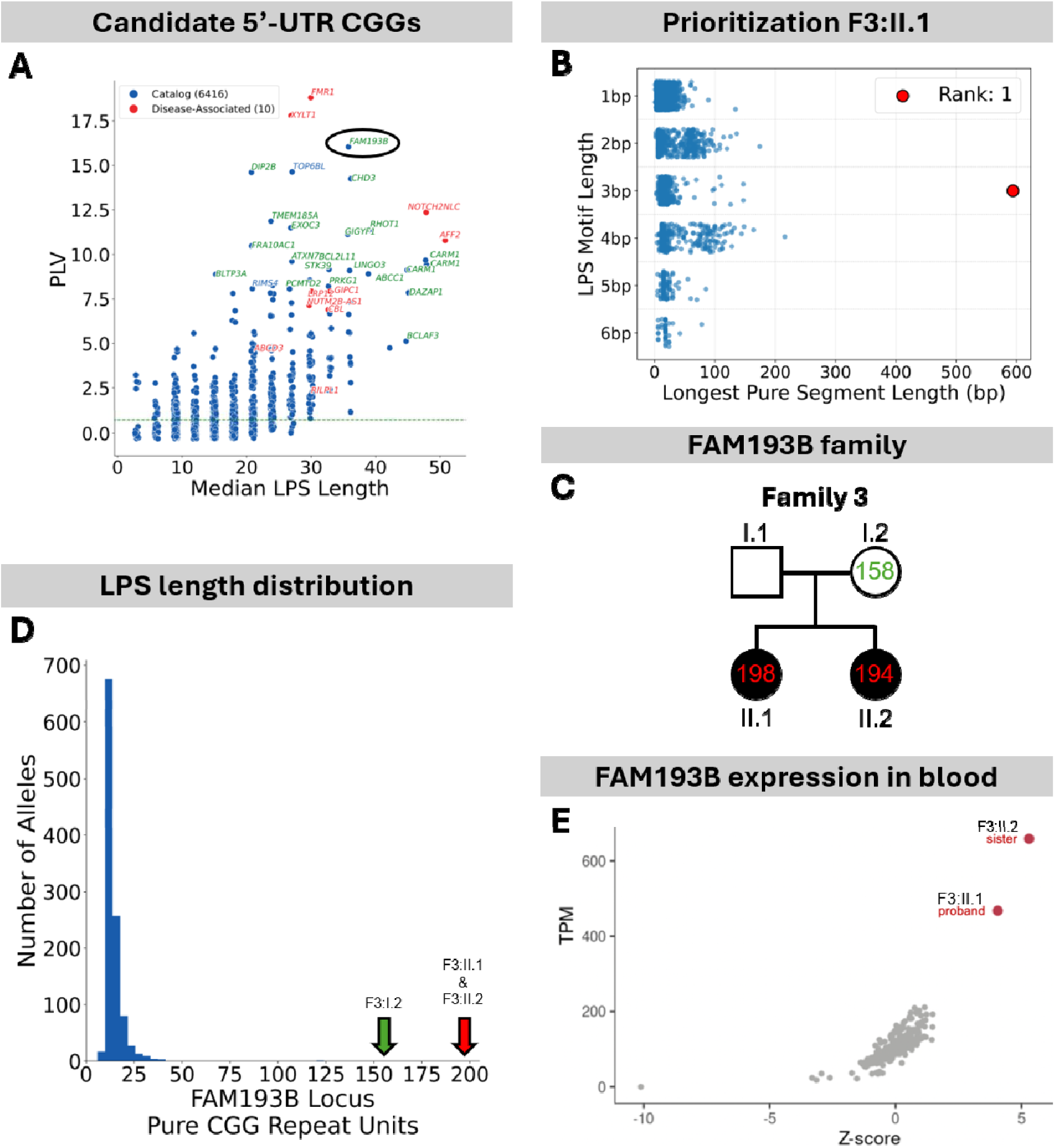
Investigation of a 5’-UTR CGG repeat in *FAM193B* as a candidate for OPDM. A) Scatterplot of the 5’-UTR CGG repeats in the discovery cohort plotting the PLV score (y-axis) against the median LPS length (x-axis). Disease-associated STRs have their gene names written in red. Other seeming outlier loci have their gene names written in blue (if they did not replicate in the validation cohort) or green (if they replicated in the validation cohort). B) Swimlane plot showing the prioritization of the *FAM193B* STR in the proband of Family 3 by combining PLVI and 99.9 percentile STR outliers. C) Pedigree of the family with oculopharyngodistal myopathy (OPDM). Number of CGG repeats measured at the *FAM193B* locus is given for each measured family member. D) Histogram of LPS length for the CGG motif at the *FAM193B* locus in the discovery cohort. Red arrows mark the allele sizes observed in the two affected individuals. E) Scatterplot of transcript expression of *FAM193B* in blood samples from the two affected sisters of Family III against the distribution of *FAM193B* expression in the peripheral blood of 282 control samples.

The highest-ranking candidate locus uncovered with this approach is a CGG repeat in the 5’ UTR of the *FAM193B* gene (circled in Figure 6A). The expansion of several CGG repeats located in 5’ UTRs have been implicated in oculopharyngodistal myopathy (OPDM): *NOTCH2NLC*, *GIPC1*, *LRP12*, *RILPL1*, and *ABCD3*. Analysis of approximately 50 existing LRS samples of OPDM patients identified an expansion of the *FAM193B* CGG repeat prioritized as the top-ranked allele in one affected individual (Figure 6B) and predicted to be pathogenic by RExPRT (score of 0.997)^44^. In this family (Family 3), two siblings presented with OPDM at ages 49 and 51, while their parents were unaffected (see Supplementary Results for complete details) (Figure 6C). We identified pure CGG repeat alleles of 198 and 194 copies in the two affected siblings inherited from a shorter tract of 158 pure CGG repeats in the unaffected mother. The longest allele in the father was 16 pure CGG repeats (biallelic genotypes for all four individuals presented in Supplementary Results). Each of these expansions represent rare outliers, longer than any observed in the general population (Figure 6D; Supplementary Figure 17A) based on analysis of the *All of Us* long-read and short-read cohorts of 243,043 population controls (Supplementary Figures 17A, B). Follow-up analysis of blood derived DNA and RNA from this family demonstrated that *FAM193B* was not methylated by this expansion and expression was strongly upregulated, compared to 282 control blood samples (Figure 6E). This supports a disease mechanism analogous to other OPDM REDs including *NOTCH2NLC*, *GIPC1*, *LRP12*, *RILPL1*, and *ABCD3* which exhibit a toxic gain-of-function through transcript over expression and non-AUG RAN translation^45,46^. While additional OPDM patients carrying a *FAM193B* expansion will be required to confirm these findings, we implicate this locus as a strong candidate for disease.

## Discussion

The genetically diverse variant class of STRs has resisted routine genome-wide investigations in genotype-phenotype studies. Given the still significant diagnostic gap in rare diseases and the lack of molecular understanding of most GWAS signals, a pressing need exists to further explore the contributions of STRs to human diseases and traits. Unlike protein-coding single nucleotide variants, where allele frequencies, ancestral effects, and genetic constraint are well-characterized^4^, high fidelity, base-pair resolved STR genotypes are just beginning to emerge at scale. Compared to SNVs, STRs have additional layers of complexity, including motif turnover, repeat length variation, and repeat motif purity — all of which have been shown to be relevant for disease association^47^. The resource we developed based on LRS will assist in removing many of the prior obstacles and suggests powerful strategies for the discovery of disease-linked STRs.

We have created the most comprehensive, multi-ancestral dataset of STR alleles to date, which details, at base-pair resolution, the motifs, interruptions, and length distributions at 4.4 million loci. The complete dataset can be accessed through a featured workspace in the *All of Us* Researcher Workbench, while PLVI values, length distributions, and other summary statistics are available in TR-Explorer^5^ and downloadable through Zenodo^6^. We use this dataset to investigate several domains of interest: genetic ancestry differences and similarities (Figures 1B-C, 3C, Supplementary Figure 6B), HiFi vs ONT technology biases (Figures 1, 2A-B, 3C, Supplementary Figures 1, 2, 3, and 9), long-read vs short-read differences (Supplementary Figures 13, 23-27), and coverage depth-dependent effects for HiFi genomes (Supplementary Figures 1, 2, 6, and 9). We show that 99.93% of the loci in the TR-Explorer catalog have some degree of length variation, and even more (99.95%) exhibit sequence variation. Yet only 3.3% exhibit sequence changes large enough to alter the LPS motif in the 6,290 haplotypes studied. This subset shows extraordinary levels of motif variability, such that we have not yet observed a saturation in novel allele sequences or novel LPS motifs in our cohorts. This resource of over 23 billion STR alleles will provide a critical baseline for future population studies.

Further we introduce a robust metric to facilitate the discovery of disease-causing STRs, which we dub the Pure Length Variability Index (PLVI), based on the observation that disease-causing STR loci are consistently among the most highly variable loci for their motif length in the population. We replicate this finding across three long-read cohorts, four catalogs of repeat loci, the major genetic ancestry groups, and HiFi, ONT, and even Illumina sequencing technologies. While the PLVI signature does not apply to the poly-alanine STRs or the two diseases caused by single repeat unit changes, it constitutes a highly sensitive and specific association for disease-associated STRs. We hypothesize that this signature exists because currently known disease-associated STRs represent mutable elements near a functional optimum. They are driven by selection to acquire moderately long, pure segments, as this provides evolutionary advantage. This has been suggested for huntingtin (Huntington disease) and cognitive function^48^, and CAG repeat length in the androgen receptor (Spinal and Bulbar Muscular Atrophy) tuning androgen signaling^49^. However, long, pure segments are unstable, which drives the high population-level variation we observe despite selection against very large segments, as they cause disease (see Supplementary Results). Rare disease STR studies thus far have essentially identified a subset of loci with high PLVI by detecting uniquely expanded events linked to a phenotype. This interpretation would explain the observation that the PLVI is robust across ancestries, even in cases where the disease manifests primarily in a single ancestry group. In practical terms, it is well documented that several REDs primarily occur on ancestry-specific haplotypes, yet our results show that these loci can be prioritized by PLVI in any ancestral group studied (Figure 3D). In addition, this ‘goldilocks’ reasoning does imply that high PLVI loci may be involved in more common traits and diseases, which are typically identified in GWAS. In fact, we show for three of the very few known GWAS loci with an STR as the functional foundation, that these STR had a high PLVI (Figure 4B). We thus predict that the PLVI measure will be an effective prioritization tool in future studies that aim to identify the underlying molecular driver allele in GWAS signals.

As proof-of-principle, we applied the STR allelic catalogue in combination with the PLVI approach to unresolved rare disease genomes. As shown in Figures 5 and 6, in single LRS genomes, we were able to rank *EP400* in third place and *FAM193B* as the number one candidate gene for cerebellar ataxia and OPDM respectively. Familial segregation, RNA expression data, and the identification of an unrelated family for *EP400* provided additional support although more families will be needed to firmly establish a pathogenic association. Nevertheless, the key advance is that genomic mutational properties are used to identify the most plausible disease loci. Associations of STR with disease are now feasible in the rarest of phenotypes and depend less on statistical linkage to a genomic interval in large or multiple families.

Our study has several limitations which will need to be addressed in future work. First, we excluded variable number tandem repeats (VNTRs) from our analysis. VNTRs are tandem repeats with motif length over 6 base pairs. They represent additional challenges beyond STRs due to their greater degree of motif variation, nearly unlimited motif space, and still-developing VNTR catalogues. We thus do not expect that the PLVI will simply extend into the VNTR field. Second, our validation dataset had relatively low coverage depth. Nevertheless, even at modest LRS depth (∼12x), the repeat length distributions are highly correlated to the 30x discovery cohort (Figure 1B). By including an ONT dataset from the 1000 Genomes Project, we further provide dual replication of the allelic metrics and the effects of PLVI. As high-quality LRS datasets will grow in size and ancestral diversity, our findings will be refined and extended. The principal discoveries, however, are well founded and replicated. Finally, any novel gene – phenotype correlation requires significant evidence. Due to the character of this manuscript and the focus on providing much needed STR metrics for the entire field, we show still somewhat limited evidence for *EP400* and *FAM193B*. These findings are, however, in line with the increasing rarity of novel gene identifications, such as *TBC1D7*^50^, *NAXE*^51^, *RAI1*^52^, and *TYMS*^53^, which each identified 1-3 families carrying the expansion in their initial publications. In this sense, we aimed to demonstrate the opportunity of identifying STR loci in few LRS genomes. Follow up studies on these candidate genes will be required.

In summary, we present the largest base-pair resolved STR catalogue to date in conjunction with the definition of a highly impactful ranking system via Pure Length Variability Index (PLVI). These results remove major methodological hurdles of STR – phenotype association studies and unlock the potential for biological discoveries in the genomic space of STRs.

## Supporting information

Supplementary Results

## Methods

### Long-Read Sequencing

First, a cohort of 1,027 individuals who self-identified as black or African-American were selected for HiFi sequencing to a median coverage of 8x on Sequel II machines as part of the *All of Us* Research Project Long Reads Working Group’s phase 1 effort. These samples were all sequenced at the Hudson Alpha sequencing center. These samples are the subject of Garimella, et al^7^ and were released in CDR v7 in the *All of Us* Researcher Workbench.

Second, a cohort of 1,075 individuals representing a variety of ancestry groups were selected for HiFi sequencing to a minimum coverage of 12x (actual median 13x) as part of the *All of Us* Research Project Long Reads Working Groups phase 2 effort. None of these individuals had over 25x sequencing coverage. These samples were sequenced at the Hudson Alpha and Broad Institute sequencing centers and were released in CDR v8 in the *All of Us* Researcher Workbench. These samples were then combined with the 1,027 samples from CDR v7 to form the validation cohort in this manuscript.

Third, a cohort of 543 individuals were selected for HiFi sequencing with a minimum coverage of 25x (actual median 32x) as part of the *All of Us* Research Project Long Reads Working Group’s phase 2 effort. These samples were sequenced at the University of Washington and Baylor College of Medicine sequencing centers and were released in CDR v8 in the *All of Us* Researcher Workbench. These samples compose the discovery cohort in this manuscript.

### Genome Alignment and TRGT Processing

All HiFi samples were genotyped with TRGT v5.0.0^8^ against the TR-Explorer v1.0.1 catalog^5^ of 4.4 million loci on GRCh38. This was chosen as the primary catalog for its completeness, precise boundaries, and novel ‘variation cluster’ system for determining when neighboring loci should be analyzed together or separately.

### TRGT post-processing to calculate longest pure segment length

The longest pure segment (LPS) was then calculated for each allele by using regular expressions to identify segments of perfect repetition of various input k-mers. The reference motifs provided in the catalog were used as the initial seed set of k-mers and were evaluated first. If any segment composed of a repeating reference motif accounted for over half of the allele’s length, then that motif was used as the LPS motif and the length of that segment was used as the LPS length. In most alleles, this was able to allow quick resolution of the LPS motif and length. When no reference motif met this condition, all k-mers up to 20bp long which occurred at least 3 consecutive times were searched over with the same methodology used for the reference motifs.

### LPS Motif Simplification

In this schema, motifs are grouped together that are equivalent by any process of reduction, shifting, or reverse complementation and then are written in minimal alphabetical order. To create statistically supported distributions of repeat variations, we also only used the most common LPS motif observed in alleles at each locus (presumed wildtype).

### PLV and PLVI Definition

The Pure Length Variability (PLV) score is the standard deviation of the LPS length of a TR locus, considering only the most common LPS motif. The Pure Length Variability Index (PLVI) is then calculated by grouping loci by their motif length (using the most common LPS motif for each locus) and then ranking loci within each group by the PLV score.

### Statistical Analyses

The Mann-Whitney U Test, Fisher’s Exact Test, Spearman Correlation, and Pearson Correlation were conducted using scipy version 1.10.1 running in python 3.10. Cohen’s D effect size was computed in python 3.10.

## Resource Availability

The data created through the *All of Us* Program Long Read Data release CDRv7 (April 2023: https://support.researchallofus.org/hc/en-us/articles/14769699298324-v7-Curated-Data-Repository-CDR-Release-Notes-2022Q4R9-versions) and CDRv8 (February 2025: https://support.researchallofus.org/hc/en-us/articles/30294451486356-Curated-Data-Repository-CDR-version-8-Release-Notes) are available through the *All of Us* Research Program researcher workbench (https://researchallofus.org/). Code to reproduce the results presented in this manuscript is available in the corresponding Github repository (https://github.com/ZuchnerLab/AoU_TRs_PLVI_Manuscript). The code as well as the resulting data files are available through a featured workspace in the *All of Us* Research Program researcher workbench (https://workbench.verily.com/workspaces/longreads_trs_plvi_manuscript/). Locus-level summary tables of allele length and LPS length distributions are available through Zenodo (https://doi.org/10.5281/zenodo.19239224)^6^. Additional data that support the findings of this study are available on request from the corresponding authors (M.C.D. and S.Z.).

## Acknowledgements

This work was supported by the National Institutes of Health (NIH) National Human Genome Research Institute (grant R21HG013397 to M.C.D. and S.Z.), the *All of Us* Research Program (1OT2OD037907 to S.Z.), the Undiagnosed Disease Network (5U01NS134353 to S.Z.), and NINDS (5R01NS072248 to S.Z.). Research reported in this publication was supported, in part, by the National Human Genome Research Institute of the National Institutes of Health (NIH) under Award Number OT2OD038111 (to C-L.W., E.E.E.) and by the Medical Research Future Fund (MRF2024989) to P.J.L., M.B., and H.R. and National Health and Medical Research Council (GNT2028681) to P.J.L. D.E.M. is supported by NH grant DP5OD033357. The content is solely the responsibility of the authors and does not necessarily represent the official views of the NIH. This work was also supported by an NHMRC Ideas Grant to G.R. (APP2002640). G.R. is supported by an Australian NHMRC Investigator Grant (APP2007769), the Patricia Kailis Fellowship in Rare Diseases and Hearts and Minds Investment Limited (HM1). E.E.E. is an investigator of the Howard Hughes Medical Institute. We thank all the individuals who participated in this study. We gratefully acknowledge *All of Us* participants for their contributions, without whom this research would not have been possible. We also thank the National Institutes of Health’s *All of Us* Research Program for making available the participant data examined in this study. D.P. holds a Fellowship award from the Canadian Institutes of Health Research.

