## Supplementary Results for "Population-scale variability at short tandem repeat loci reveals pathogenicity signature"

**Supplementary Information**

### **SUPPLEMENTARY METHODS**

#### **Sequencing cohort generation**

##### *HiFi (AoU)*

First, a cohort of 1,027 individuals who self-identified as black or African-American were selected for HiFi sequencing to a median coverage of 8x on Sequel II machines as part of the All of Us Research Project Long Reads Working Group’s phase 1 effort. These samples were all sequenced at the Hudson Alpha sequencing center. These samples are the subject of Garimella, et al^1^ and were released in CDR v7 in the All of Us Researcher Workbench.

Second, a cohort of 1,075 individuals representing a variety of ancestry groups were selected for HiFi sequencing to a minimum coverage of 12x (actual median 13x) as part of the All of US Research Project Long Reads Working Groups phase 2 effort. None of these individuals had over 25x sequencing coverage. These samples were sequenced at the Hudson Alpha and Broad Institute sequencing centers and were released in CDR v8 in the All of US Researcher Workbench. These samples were then combined with the 1,027 samples from CDR v7 to form the validation cohort in this manuscript.

Third, a cohort of 543 individuals were selected for HiFi sequencing with a minimum coverage of 25x (actual median 32x) as part of the All of Us Research Project Long Reads Working Group’s phase 2 effort. These samples were sequenced at the University of Washington and Baylor College of Medicine sequencing centers and were released in CDR v8 in the All of Us Researcher Workbench. These samples compose the discovery cohort in this manuscript.

##### *ONT (1KG)*

Five hundred samples generated by the 1KGP Long Read Sequencing Consortium^2^ were collected for use as the replication cohort in this manuscript. These samples had a median coverage of 35x.

##### *srWGS (AoU)*

The All of Us Research Project CDR v8 contained a release of nearly 250,000 short-read whole genome sequencing (srWGS) samples. From that cohort, we genotyped the *FAM193B* and *EP400* candidate loci in 243,043 using ExpansionHunter. We also genotyped a larger set of ~170,000 loci in 689 srWGS samples which were also individuals with HiFi sequencing data in CDR v7.

#### **Tandem Repeat Genotyping**

All HiFi samples were genotyped with TRGT v5.0.0^3^ against the TR-Explorer v1.0.1 catalog^4^ of 4.4 million loci on GRCh38. This was chosen as the primary catalog for its completeness, precise boundaries, and novel ‘variation cluster’ system for determining when neighboring loci should be analyzed together or separately.

Additionally, the HiFi samples were genotyped with TRGT two other catalogs. First, the Adotto v1.0 catalog of 1.7 million loci^5^ was used as a complement to the TR-Explorer catalog in order to verify that the observed association between allelic variance and disease association was not catalog-specific. This catalog was chosen for its broad locus definitions and permissive inclusion criteria, which provided a distinct counter to the approach taken by the TR-Explorer catalog. Finally, a smaller catalog of ~170,000 loci measured to be polymorphic in the 1000Genomes dataset was used for a direct comparison of TRGT calls to ExpansionHunter^6^ calls because this catalog was available for both pieces of software when this project began.

#### **Allele post-processing**

##### *LPS calculation*

TRGT and Medaka each produce consensus allele calls for each locus where they emit a result. Both programs were provided the individual’s sex so they emitted the correct number of alleles for the X and Y chromosomes. The Longest Pure Segment (LPS) was then calculated for each allele by using regular expressions to identify segments of perfect repetition of various input k-mers. The reference motifs provided in the catalog were used as the initial seed set of k-mers and were evaluated first. If any segment composed of a repeating reference motif accounted for over half of the allele’s length, then that motif was used as the LPS motif and the length of that segment was used as the LPS length. In most alleles, this was able to allow quick resolution of the LPS motif and length. When no reference motif met this condition, all k-mers up to 20 bp long which occurred at least 3 consecutive times were searched over with the same methodology used for the reference motifs.

##### *Generation of the statistics tables*

After the LPS length and motif had been calculated for each allele, we compiled that information, along with the raw allele sequences and overall allele lengths into large “allele info” tables containing all the individuals in a cohort, with separate tables for each cohort. These “allele info” tables were split by chromosome for convenience. Statistics were generated by first loading a cohort’s “allele info” table into memory using the polars library in python and then computing the necessary percentiles, standard deviations, means, and median absolute deviations. For each cohort, this was done first operating on the overall allele length values to create allele length stats tables. Those tables included a measure of the unique sequences observed in addition to the standard columns shared among all the tables.

Second, the operations were repeated on the LPS length column without consideration of the LPS motif, which created the per-locus LPS stats tables. Those tables contained a measure of the composition polymorphism score of each locus. The Composition Polymorphism Score (CPS) serves as a measure of motif heterogeneity observed at a TR locus and was defined in the TRGT manuscript^7^. Specifically, for each TR locus, we calculated the Jaccard similarity between each pair of observed LPS motifs, weighted by the number of haplotypes in which that motif was observed as composing the LPS. The CPS was then the sum of all weighted Jaccard similarity scores divided by the total number of possible pairs at that locus. The CPS score ranges from 0 to 1 with low values indicating consistency in the LPS motif across samples and high values indicating high levels of heterogeneity in the LPS motif across samples.

Third, the statistics operations were repeated on the LPS length column separately for each LPS motif, which created the per-motif LPS stats tables. These tables contain more rows than the others since each locus can exhibit multiple LPS motifs. This table has an additional column specifying the LPS motif for each row.

These three tables were created for each cohort and catalog combination. Additionally, in the validation cohort, the tables were computed again using the subsets of individuals belonging to each predicted ancestry group.

#### **Medaka Genotyping of ONT data**

The replication cohort of ONT samples were analyzed with the TR-Explorer v1.0.1 catalog using Medaka Tandem^8^.

#### **Comparison of srWGS to HiFi**

We genotyped TRs in 689 matched pairs of samples with both short-read and long-read sequencing. These 689 samples were a subset of the total set of 1,027 samples. TRs in short-read genomes were genotyped with ExpansionHunter (EH) and ExpansionHunter Denovo (EHDn) and TRs in long-read genomes were genotyped with TRGT. We compared repeat sizes of TRs genotyped by the two different methods, and using TRGT as the ‘truth set’, we calculated precision and recall values for EHDn calls.

For each sample, we filtered EHDn results to select for TRs with at least five anchored in-repeat-reads (IRRs). Next, we used bedtools intersect to find the intersection of any TRs identified by EHDn and TRGT, as well as those that were unique to each tool. For intersecting TRs, we compared motifs as well as reads/sizes produced by each tool. We categorized the EHDn TRs into four groups:

A. Expanded: EHDn TRs that are above the base pair size threshold according to the TRGT calls.
B. Not expanded: EHDn TRs that are below the base pair size threshold according to the TRGT calls.
C. Motif mismatch: EHDn TRs that have a different motif from the corresponding TRGT calls.
D. De novo: EHDn TRs that are not found in the adotto catalog used for the TRGT calls.

The base pair thresholds we used for comparison were 125, 150, 175, 200, and 225 bp. To calculate the sizes using the TRGT data, we counted the lengths of segments of major motifs, allowing for ‘fuzzy matching’. To check if a TR crossed the threshold, we added together the sizes of all segments in that region matching the motif of interest. Using the specified categorizations above, we calculated the precision and recall rates of EHDn at these different thresholds. These metrics were defined as:

1. $\boldsymbol{Precision=}\frac{\boldsymbol{Expanded TRs}}{\boldsymbol{Expanded TRs + Not expanded TRs + Motif mismatched TRs + De novo TRs}}$
2. $\boldsymbol{Recall=}\frac{\boldsymbol{Expanded TRs}}{\boldsymbol{TRGT TRs \geq bp threshold}}$

Note that when calculating recall, we only used the larger allele for TRGT data since this dataset is allele specific while EHDn only provides a single read count per TR locus and does not contain information on zygosity. We also repeated this analysis using a threshold of 175 bp, and filtering for EHDn TRs with at least 10 and 15 anchored IRRs.

*Comparison between de novo and cataloged TRs*

We combined all the TRs above five anchored IRRs for all samples and separated them into de novo TRs, and all other TRs (cataloged). Next, we used bedtools merge to combine any overlapping TRs within groups. To investigate which types of repeats are represented in the different groups, we downloaded the Simple Repeats track from UCSC’s Table Browser. This dataset categorizes repeats by coordinates into different groups. After calculating frequencies of TRs in the different categories, we computed fisher’s exact test odds ratios for each category, comparing the two broad groups (de novo and cataloged).

We also calculated frequencies of different motifs within the two groups and their corresponding odds ratios. The graphs represent only significantly enriched motifs within any of the groups. Finally we investigated the nucleotide compositions of the motifs and calculated the frequencies and odds ratios in both groups.

*Comparison between recalled and unrecalled TRs*

We used bedtools intersect to compare EHDn and TRGT TRs for each sample. The resulting files were concatenated then split into recalled and unrecalled groups. Recalled TRs were defined as any EHDn TRs that intersected with TRGT TRs. Bedtools merge was used to combine any overlapping TRs within groups. We investigated nucleotide compositions of the motifs and calculated their frequencies and odds ratios.

To calculate the GC percentage, we took the complete TRGT sequence for each allele, removed the 25 bp flanking sequence on either end and counted the number of G and C nucleotides in the sequence. We then divided this by the sequence length, and this gave us the GC content of the allele. We compared the GC content of alleles in the two groups and performed a Wilcoxon test to assess significance and calculated a Cohen’s d effect size.

To calculate purity, we again removed the flanking 25 bp sequence on either size of the repeat from each allele. We then identified the most common motif within the allele by taking the motif which occurred most frequently. We then counted the number of times the motif occurred and divided it by the length of the sequence and multiplied by 100 to get the percentage purity.

The longest pure segment size was calculated by taking the major motif segments from previous processing and identifying the longest segment of these and calculating its size. These segments allow for some fuzzy matching, or minor interruptions. Finally, the number of unique motifs was calculated by counting the number of motifs per allele. This was counted from the major motifs identified, which was part of previous data preprocessing.

#### **Segmentation algorithm**

The segmentations of the allele sequences depicted in the waterfall plots throughout the manuscript were determined as follows. First, we collect all k-mers within a specified size range (typically 1-20 bp, unless otherwise noted), reducing each k-mer to its lexicographically simplest form and then keeping only the unique remaining k-mers. We also remove redundant k-mers that would be equivalently stated as a shorter sequence. For example, AAGAAG would be reduced to simply AAG. Second, we filter out k-mers that only appear as subsequences of other k-mers. For example, if there are no ‘AG’ occurrences outside of ‘AAG’, then ‘AG’ would be excluded. This prevents mono- and di-nucleotide k-mers from being unnecessarily prevalent. Third, we take this list of candidate k-mers and sort them by their length (longest first) and then by frequency of occurrence at this locus across the cohort (most common first). Finally, we iteratively mask each allele sequence according to its matches with each k-mer in the list, going in the order described. If a segment is attributed to a k-mer earlier in the list, it cannot be claimed by a later k-mer. This prevents shorter k-mers that are part of longer ones from inappropriately masking out regions that would be better described by the longer motif. For example, a region that would be best described as (AAG)_n_(AAGAAGAGG)_n_(AGG)_n_ needs to be segmented by the AAGAAGAGG motif before it is segmented with the AAG or AGG motifs in order to reach this optimal answer. For plotting clarity, we typically only utilize the 10 most common k-mers for segmentation and leave the remaining sequences as ‘other’.

#### **Discovery curve algorithm**

The discovery curves of either novel allele lengths, novel allele sequences, or novel LPS motifs were all generated through Monte Carlo sampling of the analyzed cohort of alleles. To do this, we first collected the number of observations for each unique instance of the factor being “discovered” (either the allele length, the allele sequence, or the LPS motif sequence). We then sampled the discovery factor at a given locus from this distribution without replacement and recorded how after how many individuals we observed each new occurrence (each novel allele length, allele sequence, or LPS motif sequence). We then accumulated these results across all the loci in the catalog. Finally, we repeated this process 1,000 times and plotted the mean of the results. a

#### **Selection of disease-associated STRs**

To determine the set of disease-associated STRs used in this study, we began with the complete set of disease-associated tandem repeat loci on STRchive^9^ on January 1^st^, 2026. We then removed the VNTR loci and the loci coded as yellow or red, indicating lower levels of support for the disease relationship. For the work presented in the main manuscript, we further removed the poly-alanine loci (*ARX*, *NIPA1*, *HOXA13*, *PHOX2B*, *SOX3*, *HOXD13*, *ZIC2*, *RUNX2*, *PABPN1*, and *FOXL2*) and the two loci which are pathogenic from single copy number changes (*MIR7-2*, and *COMP*), although we do discuss those loci later in the supplementary results.

#### **Methodology of identifying enrichment of disease-associated STRs among the most polymorphic loci**

Enrichment of disease-associated STRs among the sets of the most polymorphic loci was quantified primarily through Mann-Whitney U Tests (for determining which factors increased enrichment) and Fisher’s Exact Tests (for examining specific percentile cutoffs). We tested a variety of factors for their effect on this enrichment, including standard deviation of total allele length, median allele length, difference between 99.9^th^ percentile and median allele length, median absolute deviation of allele length. We also inspected versions of each of these measures looking at LPS length instead of total allele length and then repeated the analysis using only the most common LPS motif for the LPS length tests. All of these tests were also performed with normalization to repeat motif and repeat motif length: for the allele length-based tests, the reference motif was used for each locus, while for the LPS-based tests, the LPS motif was used.

#### **Linkage regions analysis**

Published linkage regions were collected from their respective manuscripts and lifted over to GRCh38 coordinates when necessary. The standard deviation of the most common LPS motif was collected for each locus within the linkage region and then plotted. The rank of the pathogenic STR overall and within its genomic biotype (5’UTR, coding, intronic, 3’ UTR, or intergenic) was recorded.

#### **GWAS regions analysis**

The three GWAS hits now known to be driven by STRs were collected from the NCBI GWAS catalog. For each locus, we collected all STRs in our catalog within 50kb. The standard deviation of the most common LPS motif was collected for each locus within the linkage region and then plotted. The rank of the causal STR overall and within its genomic biotype (5’ UTR, coding, intronic, 3’ UTR, or intergenic) was recorded.

#### **Outlier identification analysis**

First, the 99.9^th^ percentile of allele length was recorded for each catalogued locus using the discovery cohort. For each sample, we then counted how many alleles in that individual surpassed this threshold.

Second, we repeated this process but using the LPS length instead of the total allele length. This required matching of the LPS motif in an individual with at least one individual in the discovery cohort.

Third, we further filtered the LPS-length outliers to only the set of loci which are also in the top 5 percent of PLVI scores.

#### **Outlier prioritization analysis**

Individuals with rare diseases had either HiFi or ONT sequencing performed and their tandem repeats genotyped for the TR-Explorer v1.0.1 catalog using either TRGT or Medaka, respectively. Their longest pure segment lengths and motifs were called as was done for the discovery, validation, and replication cohorts. Then, they were subjected to outlier filtering as described above. The final set of outlier STR alleles were then prioritized according to the allele’s LPS length. The rank of the pathogenic STR expansion was noted both overall and within motif length group. The lengths of these outlier STRs were then plotted using swimlane plots to separate them by motif length.

#### **TR Constraint**

##### *Model training*

The pre-trained Nucleotide Transformer V3 model with 650 million parameters was downloaded from Hugging Face (<https://huggingface.co/InstaDeepAI/NTv3_650M_pre>)^10^. Using Pytorch Lightning, the model was fine-tuned to simultaneously predict six regression tasks for each input TR locus. These tasks were the values of 1) the standard deviation of the length of the longest pure segment (LPS) for the most common LPS motif; 2) the LPS motif variability; 3) the number of unique motifs observed as the LPS motif; 4) the standard deviation of the length of the total span of the locus; 5) the standard deviation of the LPS across all motifs observed at that locus; and 6) the Composition Polymorphism Score (CPS) as a measure of motif variability. TR loci within 10kb of a gene as defined by Gencode genes basic annotation version 49 were held out. Loci more than 10kb away from a gene were then split into a training dataset composed of 90% of the loci and a validation dataset composed of 10% of the loci. For each locus, the Nucleotide Transformer V3 model was given the tandem repeat sequence as present in the GRCh38 reference genome padded by 75 bp on each side. The training and validation sets were further filtered to remove data outliers in the following manner: the values for measures 1, 4 and 5 had to be below the 99th percentile in the training set and the values for measures 2, 3, and 6 had to be below the 99.9th percentile in the training set. Values were then standardized to facilitate prediction (training mean subtracted and then divided by the standard deviation in the training set).

The model was trained for 10 epochs using the AdamW optimizer set with a learning rate of 1e-6. The loss function was the sum of the mean-squared error terms of each of the six regression heads. Performance on the validation set was recorded after each epoch. The trained model was then used on the set of loci within 10kb of genes to predict the “expected” values of variation for each of the six measures, though only measure 5 is utilized for the manuscript.

##### *Model evaluation*

Performance was evaluated first on the technical basis of the spearman correlation between the predicted and observed values for each of the six regression heads on the held-out validation set of loci more than 10kb away from genes, which should be largely under neutral selection. This is meant to assess the degree to which the model is able to accurately predict the behavior of STR loci under neutral selection.

After making predictions on the set of loci within 10kb of genes, a second assessment of the model’s ability to accurately predict the behavior of STR loci under neutral selection was performed by analyzing the 20 autosomal CODIS loci. These loci are well-characterized to be under neutral selection^11^ and were used as a control for this same purpose in earlier analyses with the same goal^11^.

After these tests of the model’s ability to predict the behavior of STR loci under neutral selection, we evaluated the observed-to-expected ratio created by comparing the model’s output with the observed values at several different sets of biologically-relevant loci. These loci included protein-coding regions, expression-associated STRs (eSTRs) from Fotsing et al^12^, disease-associated STRs, STRs unique to humans (ab initio from Sulovari, et al^13^), and we also included the CODIS loci in these tests as well. This evaluation was meant to show a differential behavior of the model across these biologically different regions, with protein-coding regions hypothesized to be more constrained, CODIS loci to be neutral, and recently-evolved regions to be more anti-constrained. To put this performance in context, we compared the model with an analogous approach developed several years earlier by Gymrek et al^11^ as well as genic constraint from gnomAD^14^ and the reference span of the repeat to serve as a naïve baseline.

### **SUPPLEMENTARY RESULTS**

#### **Characterization of length variation**

Most prior work on genome-wide polymorphism of TRs has focused on length variation due to the limitations of short-read sequencing: even algorithms that accurately estimate allele lengths beyond read length are not able to accurately parse complex patterns of motif variation or return consensus sequences for these longer alleles. Long-read sequencing, on the other hand, is not subject to these limitations, allowing us to investigate these complexities in our dataset.

We observe a wide range of length variation among the loci in our three cohorts, though the cohorts are broadly similar to each other (Supplementary Figure 1A). Several classes of loci exhibit differing levels of variability, which is broadly consistent across the cohorts (Supplementary Figure 1B).

Since short-read data tends to struggle to accurately size TRs over ~175 bp, we investigated the frequency of TRs that are shorter than 175 bp in >99% of alleles but expand to be greater than 175 bp in <1% of alleles (computed separately for each cohort). These types of events can be difficult to catch with short-read genome data, but are important for selecting rarely expanded loci that may potentially demonstrate pathogenicity. We observe a median of 181 such rarely-expanded loci per individual in the discovery cohort (Supplementary Figure 1C), a much higher estimate than our previous findings of three rare TR expansions per individual based on short-read data^15^. However, this median value fluctuated considerably across the three cohorts.

#### **Characterization of LPS length variation**

The histogram of the pattern of LPS length variation across all the TR loci shows for each cohort a clear power law distribution with a small fraction of TRs being much more variable than the vast majority (Supplementary Figure 2a). While nearly all the TR loci in the discovery cohort demonstrate some level of length variation (4,436,423 / 4,439,672; 99.93%), we observed that TR loci detected by Sulovari and colleagues^13^ as non-repetitive in Great Apes (human ab-initio loci) exhibit significantly greater LPS length variance than the catalog as a whole (Mann-Whitney U-Test; p<1e-10; generalized odds ratio = 4.5) (Supplementary Figure 2b), revealing high levels of intra-species tandem repeat variation at tandem repeat loci unique to humans. Furthermore, we observed that disease-associated loci exhibit significantly greater levels of LPS length variance in this control population compared to the catalog (Mann-Whitney U-Test; p<1e-10; generalized odds ratio = 7.8) (Supplementary Figure 2b). This pattern also clearly replicated in the validation and replication cohorts. However, this result is not specific enough to proactively identify novel pathogenic loci, delineating them from other hyper-polymorphic loci that are unlikely to be pathogenic, such as the twenty CODIS loci used in forensic analyses. For known disease-associated loci, the distribution of allele lengths observed is provided in Supplementary File 1 and the distribution of LPS lengths observed is provided in Supplementary File 2.

We next used the motif associated with each LPS measurement for the unbiased identification of novel motifs within TRs. A saturation analysis with this approach revealed that even after examination of 543 genomes in the discovery cohort, approximately 56 novel LPS motifs were observed with each additional sample added to the set (Supplementary Figure 2c). This indicates that many more samples need to be analyzed to observe all possible LPS motifs. This effect was even larger in the validation and replication cohorts, despite the validation cohort being approximately four times larger. This suggests that the LPS motif is more likely to be spuriously called as a novel motif in ONT and lower coverage HiFi data than in ~30x HiFi data.

Analysis of the standard deviation in LPS length for loci grouped by the length of their most common LPS motif revealed that trinucleotide loci exhibited less variation than the other motif lengths, while larger loci with 12 bp or longer motifs exhibited the most variation (Supplementary Figure 3a). To determine whether this result was driven by the greater prevalence of trinucleotide loci within coding regions than other motif lengths, we repeated the analysis on the subset of loci in intergenic regions and found that the same pattern remained (Supplementary Figure 3b).

#### **Characterization of LPS motif variation**

We next compared the observed motifs that produced each LPS and plotted their frequency of divergence from the reference motif (Supplementary Figure 4A). We found that 96.6% of loci had no non-reference motifs that accounted for the LPS in any of their alleles, suggesting motif invariance. In contrast, 0.5% of loci had non-reference LPS motifs in over 90% of their alleles, suggesting either high degrees of motif polymorphism at those loci or misspecification of the reference motifs. Most interesting were the 1.7% of loci where in less than 1% of alleles, a non-reference LPS motif was observed. These loci exemplify the kind of motif polymorphism that would be hard to detect with short-read sequencing technologies and may also demonstrate pathogenicity as observed in diseases such as CANVAS, FAME, SCA31, and SCA37.

We next examined the periodicity of the LPS motifs for each locus. For example, if a locus has a 5 bp reference motif, a non-reference LPS motif that is also 5 bp long represents a substitution change. This is a distinct motif, but not a novel period. In contrast, a non-reference LPS motif of 6 bp represents an insertion, which changes the period (Supplementary Figure 4B). One further example is if a substitution occurs every other motif copy, then we get a 10 bp non-reference LPS motif. This we also consider to be of the same period as the reference motif and refer to these scenarios as “interleaved” motif changes. Inspecting the allele database in this way, we found that interleaved changes accounted for nearly two-thirds of the novel motifs, while substitutions accounted for the remainder. This pattern was broadly consistent for both the rarely altered loci and the ubiquitously altered loci (Supplementary Figure 4C-D). We did not observe any novel LPS motifs that represented a change in the period of the repeat, except in loci with multiple adjacent repeats of different motif lengths. At those loci, the novel LPS motifs represented expansions of a smaller repeat of different motif length to become the longest in the variation cluster, rather than the conversion of a repeat into one of different periodicity.

#### **Limitations of the enrichment of disease-associated tandem repeats for the most polymorphic loci**

In the main text, we described the enrichment of disease-associated STRs among the most polymorphic loci (Figure 3, Supplementary Figure 5-6), but we noted a series of exclusions: poly-alanine loci, loci pathogenic due to alterations of only a single repeat unit, and loci whose repeat motif is 7 bp or longer (variable number tandem repeats [VNTRs]). In this section, we describe the patterns observed at those loci, show the extent of their deviation from the pattern of enrichment, and speculate on the reason why they behave differently than the set used in the primary analysis.

##### *Poly-alanine loci*

There are at least 10 currently known disease-associated loci to which we refer as the poly-alanine loci on the genes *ARX*, *NIPA1*, *HOXA13*, *PHOX2B*, *SOX3*, *HOXD13*, *ZIC2*, *RUNX2*, *PABPN1*, and *FOXL2*. All of these repeats are protein-coding and produce poly-alanine chains. Since there are numerous codons that produce the alanine amino acid, these repeats include several different motifs but are broadly known under the trinucleotide pattern GCN. The loci typically have a single primary motif (usually GCA) with one or more ‘interruptions’ (as viewed at the DNA level) of other trinucleotide motifs that still start with ‘GC’. Waterfall plots showing this pattern for *PHOX2B* and *RUNX2* are presented in Supplementary Figure 7A-B.

This phenomenon of motif variability that still contributes to the poly-alanine amino acid chain obviously poses a problem for the DNA-level LPS-based analysis that we employ, but that is not sufficient to explain their divergence from the other loci. For example, the *HTT* repeat has CAA interruptions that still encode glutamine, but it is very well prioritized by our method. These poly-alanine loci exhibit moderately elevated total allele length and LPS length variation for trinucleotides (mean of 81^st^ and 75^th^ percentiles, respectively) (Figure 3c and Supplementary Figure 7C).

Two other factors may contribute to the divergent behavior of this set of loci from the rest. First, the pathogenic threshold is lower for these repeats than for the included set of disease-associates STRs. Indeed, the poly-alanine loci comprise 16 of the 20 lowest pathogenic STR thresholds (by nucleotide length) listed on STRchive. Second, these are dominant diseases with an early age of onset. Together, these factors allow strong selection against longer repeat tracts at these loci to an extent not observed for the primary set of disease-associated STRs.

##### *Single-repeat alteration loci*

There are two diseases currently considered tandem repeat diseases which are caused by a single repeat unit expansion (*COMP*) or contraction (*MIR7-2*) in the repeat loci. Because of this pathomechanism, these loci are essentially invariant in the general population. As a result, they are obviously not prioritized by our approach.

##### *VNTRs*

There are four disease-associated VNTRs that were included in the version of the TR-Explorer catalog that we used, *EIF4A3*, *MUC1*, *PLIN4*, and *VWA*. These repeats range in motif length from 10 bp (*VWA*) to 99 bp (*PLIN4*). We observed statistically significant enrichment of these loci among the most polymorphic in the catalog when prioritizing by allele length (Mann-Whitney U-Test; p<1e-2; generalized odds ratio = 11.1) or LPS length (Mann-Whitney U-Test; p<1e-3; generalized odds ratio = 22.3), but these effects were no longer significant when the measures were normalized to motif length. This suggests that while VNTRs are a much more polymorphic group than STRs when measured at the nucleotide level, most of these disease-associated VNTRs are not remarkably polymorphic relative to other VNTRs with the same motif length. This indicates that a different metric will be needed to prioritize pathogenic VNTRs from among their set.

#### **Utilization of LPS variation metrics**

##### *Linkage region analyses*

We retrospectively examined published linkage regions of 15 known pathogenic loci (Figure 4a and Supplementary Figure 8). The most recent example of this is SCA4, attributed to a GGC exonic repeat in the *ZFHX3* gene ^16–20^. The linkage region for this disorder was identified nearly 30 years prior to finding the causal TR. In this cohort, while not as variable as intergenic and intronic repeats, the GGC repeat in *ZFHX3* was the third most polymorphic coding repeat in the 7.5 Mb linkage region. The CAG repeat in *THAP11* was the fifth most polymorphic coding repeat in the region and was identified as disease-associated in 2023^21^. We repeated this analysis using the published linkage regions for 14 other disease TR loci^22–35^. We found that in 13 of the 15 cases, the disease-associated STR locus had a PLVI greater than at least 99 percent of the loci in that region. In 10 of the cases, the disease-causing STR was ranked in the top 3 repeat loci within the region when utilizing the PLVI framework. The complete set of results is given in Supplementary Table 1. This approach, in combination with the other metrics mentioned above, could have assisted in gene discovery efforts by highlighting unstable regions of the human genome. Additionally, several disorders are still only referred to by their linkage region, as the causal gene has not yet been found. Investigation of tandem repeats in these linkage regions may find pathologically expanded loci according to our methodology.

##### *Synthetic outlier prioritization analyses*

As an extension of the within-individual outlier prioritization analyses presented in Figure 4 C and Supplementary Figures 9 and 10, we decided to perform a synthetic benchmark of this approach using the alleles detected in 10 individuals from the Human Pangenome Reference Consortium project (each sequenced to at least 30x coverage with HiFi data and processed identically to the samples in the discovery cohort). Into each of these 10 individuals, we spiked in the LPS lengths of minimum pathogenic-length expansions for each of the 42 disease-associated STR loci used in this work. In each case, the minimum pathogenic size was taken from STRchive. In Figure 4D, we plot the empirical cumulative distribution of the rank of these 420 spike-in analyses, separated by the biotype of the pathogenic locus (coding, UTR, or intronic). Since we used the minimum pathogenic-length expansion for each STR, we hope this represents a strong lower-bound of performance with this approach. We found that over half of intronic disease-causing expansions achieved a rank of 1, while 90% ranked within the top 4. For 3’- and 5’-UTR expansions, half were ranked in the top 2, while 90% ranked within the top 5. Coding expansions were the most challenging for this approach, where half were ranked in the top 8, while 90% ranked within the top 16. We believe this analysis demonstrates robust performance of our approach at prioritizing many types of disease-associated STR expansions.

#### **Variation at disease associated STR loci in healthy individuals**

Among the set of 52 disease-associated STRs examined in this study, there existed a wide spectrum of allele lengths within the 543 genomes of the discovery cohort with, predictably, noncoding loci displaying the most variation in length and the largest repeat sizes (Supplementary Figure 11).

Sequence level examinations of known pathogenic loci have garnered profound insights with clinical implications in examples such as *FMR1* and *HTT* ^36,37^. In the *FMR1* 5’UTR CGG repeat locus, AGG interruptions within the CGG repeat offer stabilizing effects during transmission. The risk of premutation alleles expanding to full mutation alleles exponentially increases in alleles lacking AGG interruptions^38^. We have analyzed *FMR1* repeat sequences to assess repeat variability at high resolution in a healthy control population. The most common repeat lengths in our cohort correspond well with literature reports (29-31 repeat units, 76.3% of all alleles). There was a range of AGG interruptions from 0 to a maximum of 5 interruptions. The proportion of alleles with 0, 1, 2, 3, 4, or 5 interruptions were 3.2%, 19.3%, 75.2%, 1.8%, 0.3%, and 0.1% respectively (Supplementary Figure 12A). However, the frequency of interruptions differed between repeat length groups (Supplementary Figure 12B). Smaller alleles (14-23 units, 8.8% of alleles) predominantly consisted of 0-1 AGG interruption alleles. Alleles with 24-34 units (83.7% of alleles) mainly consisted of 1-2 interruptions. There was more variability in the number of AGG interruptions in the high normal (35-44, 5.4% of alleles) range with 31.3% of alleles having three or more AGG interruptions and also in the gray zone (45-54, 1.6% of alleles) range with 21.4% of alleles having four or more AGG interruptions. While the consequences of sequence interruptions at the *FMR1* locus have been largely elucidated, these analyses suggest that similar exploration of sequence level variation at other loci genome-wide are now feasible. Such analyses, when coupled with phenotype data, may reveal the missing heritability of complex disorders and be a valuable addition to association/modifier studies.

#### **Enrichment of disease-associated STRs among most polymorphic loci evident in srWGS data**

Since we see the enrichment of disease-associates STRs among the most polymorphic loci in the long-read data, we wondered if the signal was present in the srWGS measurements of STRs as well. To investigate this, we utilized two independent cohorts of srWGS data. The first, termed the ‘AoU srWGS cohort’, is a set of 704 samples generated as part of the All of Us Research Project and are from a subset of the individuals in the validation cohort sequenced to approximately 30x with Illumina PCR-free srWGS and genotyped with ExpansionHunter on a catalog of approximately 170,000 loci. The second was the genotype data produced by EnsembleTR on the 1000Genomes and H3Africa cohorts, which was all sequenced to approximately 30x with Illumina PCR-free srWGS^39^. For each cohort, we calculated the standard deviation of the allele length for each STR locus and then computed the rank of each locus by that measure overall as well as grouped by their reference motif length. Since neither ExpansionHunter nor EnsembleTR emits actual consensus sequences (as is the case with all srWGS TR genotyping tools that can predict alleles longer than the read length, as far as we are aware), we could not directly replicate the LPS measure on this dataset.

We found that the disease-associated STRs were indeed enriched among the most polymorphic STRs in both cohorts after normalization of the ranks to within motif length groups (AoU srWGS cohort: Mann-Whitney U-Test; generalized odds ratio: 14.3; p-value< 1e-10; EnsembleTR cohort: Mann-Whitney U-Test; generalized odds ratio: 30.8; p-value< 1e-10). This resulted in 18 of the 25 (72.0%) disease-associated STRs falling within the top 5% by this measure in the AoU srWGS cohort and 28 of the 31 (90.3%) disease-associated STRs doing the same in the EnsembleTR cohort (Supplementary Figure 13). This effect is less pronounced than in the long-read sequencing data (which had 100% of the disease-associated STRs above this threshold) but is still clearly observable and replicable across two datasets created with distinct tools and catalogs.

#### **Network analysis of genes with the most and fewest polymorphic CAG loci**

We wondered if the genes containing the coding CAG STRs with the greatest variation in LPS length differed from those containing less variable coding CAG STRs. We found that the genes containing the known pathogenic coding CAG repeats had, on average, more protein-protein interactions than genes containing coding CAG repeats from the lowest percentile of LPS variation (Supplementary Figure 14A). The 12 genes containing the other high variance coding CAGs labelled in Figure 5A also showed enrichment for increased numbers of protein-protein interactions relative to the genes with coding CAG repeats from the lowest percentile of LPS variation (Supplementary Figure 14A), though to a lesser extent than the known pathogenic TRs. We further found that the genes containing the most variable coding CAGs had fewer steps between pairs of them in the protein-protein interaction network than the genes containing the least-variable coding CAGs (Supplementary Figure 14B).

#### **Investigation into the reason disease-associated STRs are among the most variable genome-wide**

Since we have established the enrichment for disease-associated STRs among the most variable STRs genome-wide robustly across five different cohorts, three catalogs, and three sequencing technologies and have discussed the exceptions to this rule, we next sought to investigate why this is the case. We believe the answer to that question varies across the pathogenic motifs, so we will discuss them separately. This investigation is not meant to be exhaustive, but rather to highlight important salient factors driving the observed association.

Our overarching perspective is that the disease-associated STRs are functional elements which have been driven by selection to have moderately long, pure segments, as this provides some kind of advantage (which differs by repeat type). However, the STRs are also under weak selection against expanding too much, as that is associated with disease. For most disease-associated STRs, that zone in between the selective pressures for expansion and contraction is long enough that the repeats are highly unstable, which ultimately causes the high population variance that we measure. We view this as analogous to genes which are both haploinsufficient and triploinsufficient: their expression must maintain a balance between being too high and too low. The loci which achieve that balance while maintaining a short enough STR to be relatively stable then behave more like the poly-alanine and single-shift loci, which are largely stable. They have only been observed to cause diseases because their pathogenic thresholds are relatively low. We will now describe this issue for several types of disease-associated STRs in greater detail.

##### *5’-UTR CGG repeats*

The TR-Explorer catalog documents 6,530 CGG repeats in the 5’-UTRs of at 4,097 distinct genes. These repeats are well known to influence the expression of the corresponding gene^12,40,41^, acting as rheostats to enable fine tuning of the gene’s expression on shorter evolutionary timescales. In line with this, the disease-associated 5’-UTR CGGs have a much longer median length (10 repeat units) than the overall set of 5’-UTR CGGs (3 repeat units), suggesting a selection process for longer alleles at these disease-associated loci, which remained far below the pathogenic thresholds (typically around 200 repeats for many of these diseases). This observation extends to the 1% most variable 5’-UTR CGGs, which have a median length of 9 repeat units, significantly larger than the loci exhibiting the 50% least variable 5’-UTR CGG repeats (Mann-Whitney U-Test; generalized odds ratio: 276.1; p-value<1e-10). However, we also observe that these genes with the 1% most variable 5’-UTR CGGs also exhibit greater genic constraint in gnomAD than the genes with the 50% least variable 5’-UTR CGGs (Mann-Whitney U-Test; generalized odds ratio: 1.6, p-value=4.7e-3). While far from conclusive, these analyses align with our hypothesis that the 5’-UTR CGG repeats with the greatest population variance are likely under selection for greater lengths than the population overall, likely reflecting the constraint of the gene itself, mirroring a pattern reminiscent of the haploinsufficent and triploinsufficient gene.

##### *Protein-coding CAG repeats*

Protein-coding CAG repeats exhibit a similar pattern to the 5’-UTR CGG repeats: the disease-associated coding CAGs have a much longer median length (15 repeat units) than the overall set (3 repeat units). This observation extends to the 1% most variable coding CAGs, which have a median length of 9 repeat units, significantly larger than the loci exhibiting the 50% least variable coding CAG repeats (Mann-Whitney U-Test; generalized odds ratio: 9.2; p-value<1e-10). We again observe that these genes with the 1% most variable coding CAGs also exhibit greater genic constraint in gnomAD than the genes with the 50% least variable coding CAGs (Mann-Whitney U-Test; generalized odds ratio: 1.6, p-value=6.6e-4). In line with our understanding of the normal function of poly-glutamine tracts in facilitating protein-protein interactions, we observe that the genes with disease-associated coding CAGs and the candidate coding CAGs in Figure 5a both show more protein-protein interactions than genes containing coding CAG repeats from the lowest percentile of LPS variation (Supplementary Figure 14A). Our notion that there is a selective advantage driving longer poly-glutamine tracts has been explicitly described for *HTT^42^*, but we believe it may hold true across a broader set of disease-associated repeats.

One other perspective to examine in this relationship is that the intergenic CAG loci should theoretically be under less constraint than the coding CAG loci and so we would expect greater amounts of variance at intergenic CAG loci under neutral selection. But the least-variable disease-associated coding CAG repeat has higher LPS standard deviation (1.62) than the 99^th^ percentile of intergenic CAGs (1.50). In fact, the 99th percentile of LPS standard deviation for coding CAGs is actually higher (1.55) than that of the intergenic CAGs (1.50). We further observe that the median length of the disease-associated coding CAGs (15 repeat units) is longer (though not statistically significantly) than the top 1% most variable intergenic CAGs (11 repeat units; Mann-Whitney U-Test; generalized odds ratio: 2.88, p-value=1.18e-1). We believe this further supports the notion that these disease-associated coding CAGs are likely under selection driving longer poly-glutamine tracts.

##### *Intronic AAAAT repeats*

Intronic AAAAT repeats again demonstrate this same pattern: the disease-associated repeats have a much longer median length (12 repeat units) than the overall set (3 repeat units). This observation extends to the 1% most variable intronic AAAAT repeats, which have a median length of 10 repeat units, significantly larger than the loci exhibiting the 50% least variable intronic AAAAT repeats (Mann-Whitney U-Test; generalized odds ratio: 48.0; p-value<1e-10). We again observe that these genes with the 1% most variable intronic AAAATs also exhibit greater genic constraint in gnomAD than the genes with the 50% least variable intronic AAAATs (Mann-Whitney U-Test; generalized odds ratio: 1.2, p-value=2.6e-2). One interesting additional feature we observe is that the genes with the 1% most variable intronic AAAATs are enriched for having their GTEx tissue with highest median expression be in the brain relative to the 50% least variable loci (Fisher’s Exact Test; odds ratio: 2.2, p-value=8.3e-5). While there is no well-characterized physiological role for AAAAT repeats of which we are aware, we believe their matching of the same pattern suggests that they may be under some selective pressure toward longer repeat lengths.

#### **EP400 candidate investigation**

##### *Family 1 clinical characteristics*

The proband, labeled I.1 in the pedigree (Figure 5C), came from a family with a history of degenerative ataxia in his father and paternal grandfather. He was one of 5 siblings, of which 3 had the disease. He was first clinically assessed at 61 years of age. He noticed in his late 40’s that his walking was unsteady, and he started having falls. His speech became dysarthric aged late 50’s. He had to retire aged 58 years. At that age he had an ataxic gait, saccadic smooth pursuit of the eyes, cerebellar dysarthria, and a mild intention tremor. A CT scan of the brain showed atrophy of the cerebellum and pons. Four years later, he had progressed to needing a wheeled walker for mobility. He also had urinary frequency. When last seen at the age of 83 years, he was confined to a wheelchair, and he had severe dysarthria and dysphagia. He had hyperreflexia in all four limbs. He died at the age of 84 years following an aspiration pneumonia.

The daughter of the proband, labelled II.1 in the pedigree (Figure 5C), presented with tremor at the age of 40 years. One year later she developed speech difficulty. Shortly after that, her gait became unsteady. Examination showed cerebellar dysarthria, mildly unsteady gait, and brisk deep tendon reflexes in all four limbs. MRI of the brain was unremarkable. One year later, she had progressed only slightly. When she was last seen at the age of 49 years, she had a wide-based unsteady gait, marked dysarthria, saccadic smooth pursuit, and intention tremor bilaterally. She had started using a wheelchair for longer distances.

##### *Family 1 repeat expansion characteristics*

Follow-up long-read sequencing using Oxford Nanopore Technologies (ONT) instruments confirmed the expansion to contain a pure tract of 56 CAG repeats in the father and 58 CAG repeats in the daughter. Since this repeat sits within a larger poly-glutamine stretch including CAA repeats, the length of the poly-glutamine region in these individuals are 71 and 73 amino acids, respectively. The location of the repeat on the GRCh38 assembly was chr12:132062548-132062611.

##### *Family 2 clinical characteristics*

The proband, labelled II.1 in the pedigree (Figure 5C), has a 7-year history of progressive neurological symptoms beginning at age 15, initially with untidy handwriting and declining overhead tennis serve, with subsequent discontinuation of university studies due to impaired motor skills and poor mobility. Brain MRI at age 16 showed mild cerebellar atrophy, without brainstem atrophy. The mother of the proband, labelled I.2 in the pedigree, received a clinical diagnosis of spinocerebellar ataxia (SCA) in her 30s, without a genetic cause identified. She died in her 40s. Three of her seven siblings, along with their mother, also had a similar progressive SCA phenotype, along with a daughter of an affected sibling.

By 21 years of age, the proband (II.1) was no longer mobile with a gait aid, had incoordination of all limbs, with dysarthria, dysphagia and intermittent diplopia. A urinary catheter was in situ for retention, with no prior urodynamic studies. Diet consisted of finely chopped foods and thin liquids, with no prior chest infections. Clinical examination revealed hyperlordosis and left truncal lean, mild head drop, moderate ptosis, exotropia with compensatory right eyelid closure (to attenuate diplopia), horizontal nystagmus, hypometric vertical saccades, cerebellar dysarthria with high pitched hypophonia, moderately weak cough, lower limb spasticity and bradyphrenia.

Patient II.1 expressed some reluctance about further investigations, which he considered had been futile for his mother, however was accepting of further genetic testing.

Patient II.1 subsequently evolved upper limb proximal atrophy and weakness without fasciculations, excessive daytime somnolence, and declining oral intake with weight loss, with plans for gastrostomy tube insertion for feeds. Sleep study showed mild sleep disordered breathing (AHI 15), with no nocturnal hypoventilation. Poor lip seal and incoordination of tidal breathing limited interpretation of respiratory function testing, with best results including FVC 2.06L (40%) and SNIP 18 (15%). Initial genetic testing showed negative results for SCAs (1, 2, 3, 6, 7), FRDA and FMR1. Results were normal range for Vit E, coeliac serology, copper and caeruloplasmin.

##### *Family 2 repeat expansion characteristics*

Capillary electrophoresis (sizing, 31 and 77 repeats) and demonstration that not paternally inherited. Sanger sequencing revealed the sequence composition as CAG(75)CAA(1)CAG(1).

##### *Other supporting evidence of EP400 CAG expansion pathogenicity*

The distribution of the lengths of the longest pure CAG segment in our discovery cohort of 543 individuals is presented in Figure 5D. The complete allele sequences in this cohort are visualized in Supplementary Figure 15A. We further genotyped the *EP400* repeat in the 243,043 individuals from the All of Us Research Project’s short-read genome cohort, which are presented in Supplementary Figure 15B, which revealed a population distribution of repeat lengths consistent with that observed in the smaller long read cohort. *EP400* is highly constrained for SNV variation and is highly expressed in the cerebellum. The pattern of expression is reminiscent of several other genes that cause spinocerebellar ataxia through poly-glutamine expansions, including *ATXN2*, *ATXN3*, and *ATXN7* (Supplementary Figure 16). Inspection of the distribution of expression of *EP400*, *ATXN2*, *ATXN3*, and *ATXN7* in GTEx revealed extremely high Pearson correlations of median TPM value across the tissues included in GTEx. We found a correlation of 0.89 for *ATXN2*, 0.83 for *ATXN3*, and 0.84 for *ATXN7* when each was compared to *EP400*. For comparison, the median correlation coefficient between *EP400* and other genes in GTEx is 0.23. We also observed that “Brain – Cerebellum” is the tissue with the second-highest expression in both *EP400* and *ATXN2*. In fact, those two genes share six of their top 7 tissues ranked by expression. This poly-glutamine repeat in *EP400* was also predicted to be pathogenic by the RExPRT tandem repeat pathogenicity prediction AI tool with a perfect score of 1.0.

#### **FAM193B candidate investigation**

##### *Family 3 clinical characteristics*

Two sisters presented for evaluation of muscular dystrophy. The proband first noticed symptoms at age 49, while her younger sister showed symptoms starting at age 51. Both sisters exhibited bilateral upper and lower muscle weakness, facial muscle weakness, and voice changes. EMG of the proband was significant for distal myopathy. Muscle biopsy identified rimmed vacuoles and chronic myopathy. A genetic panel yielded negative results for OPDM, FSHD, myotonic dystrophy 1, and myotonic dystrophy 2. Whole exome sequencing was performed on the proband and affected sister but was uninformative. Short-read whole genome sequencing was then performed on the two sisters and their parents, which was also uninformative. Next, Oxford Nanopore sequencing was performed on the two sisters and their parents. Finally, RNA sequencing was performed on blood from the sister and proband.

##### *Family 3 repeat expansion characteristics*

Sequencing of the FAM193B family was performed as part of the Undiagnosed Diseases Network. The two affected daughters and their parents were sequenced to 30x mean coverage with ONT R9 reads. These reads were aligned to GRCh38 with minimap2, phased with Whatshap, and the tandem repeat alleles were called with Medaka Tandem using the TR-Explorer v1.0.1 catalog. Since there was significant variation among the reads spanning this locus across the individuals, Sanger sequencing was performed to confirm the allele sequence. It is from this Sanger sequencing data that the LPS was determined. The genotypes are as follows:

Proband: 15 / 198

Affected sister: 15 / 194

Mother: 16 / 158

Father: 15 / 16

Analysis of the ONT data demonstrated that this CGG repeat expansion is not methylated in any members of this family.

The location of the repeat on the GRCh38 assembly was chr5:177554489-177554531.

##### *Family 3 gene expression data results*

RNA-sequencing was performed on fibroblasts from the proband and affected sister. Unfortunately, the parents were deceased by this time and so could not be included. Comparison of gene expression in these two individuals to a cohort of control fibroblasts revealed that *FAM193B* expression was over 10 standard deviations above the mean in both.

##### *Other supporting evidence of FAM193B CGG expansion pathogenicity*

The distribution of the lengths of the longest pure CGG segment at this locus in our discovery cohort of 543 individuals is shown in Figure 6D. The complete allele sequences in this cohort are visualized in Supplementary Figure 17A. We further genotyped the FAM193B repeat in the 243,043 individuals from the All of Us Research Project’s short-read genome cohort, which are presented in Supplementary Figure 17B, which revealed a population distribution of repeat lengths consistent with that observed in the smaller long read cohort. All control chromosomes carried fewer than 50 repeats, except one unaffected individual with ~124 repeat units. Further, this CGG repeat locus was predicted to be pathogenic by RExPRT with a score of 0.997. This gene was also prioritized as a strong pathogenic candidate for this particular family by Watershed-SV, a tool that incorporates expression data into structural variant prioritization models. That tool predicts that the expansion causes overexpression of the *FAM193B* gene.

#### **TR Constraint**

##### *TR Length Constraint results*

Finally, we sought to integrate all that we learned about TR variation in the above analyses into a formal metric to further refine interpretation of any TR locus and aid in the discovery of future novel pathogenic TRs. Given the success of gene-level measures of constraint in prioritizing disease genes in recent years, we aimed to develop an analogous measure of constraint to prioritize novel pathogenic TRs.

We fine-tuned a Nucleotide Transformer v3 “DNA language” model to predict the standard deviation of LPS length when given the reference sequence of a TR, padded by 75 bp on each side (Supplementary Figure 18A). This deep learning model was trained on TRs more than 10kb away from the nearest gene, which are presumably largely under neutral selection. Validation was performed on a held-out set comprising 10% of those loci. The model achieved Pearson correlation coefficient of 0.808 for LPS length variation on the validation dataset.

The model was then used to generate ‘expected’ length and motif variation rates for all TRs within 10kb of genes. These were compared to the observed variance rates to generate an observed-to-expected ratio, analogous to gnomAD’s measure of genic constraint (Supplementary Figure 19a). The complete results are available in Supplementary File 3.

We observed that, similar to non-synonymous SNVs in genes, most TRs exhibit moderate negative selection for length variation (observed-to-expected ratio below 1) (Supplementary Figure 19a). Encouragingly, the set of CODIS loci, which are highly polymorphic but under neutral selection for length changes, were well-predicted by the length constraint model, achieving a Pearson correlation coefficient of 0.603 on this set of loci (Supplementary Figure 19b). We also found that the predicted length variation was highly correlated with an earlier STR Constraint measure developed by Gymrek and colleagues^11^ (Pearson correlation coefficient of 0.836) (Supplementary Figure 19c). Evaluation of performance on the validation set of loci showed consistent ability to predict STR length variation regardless of the motif length of the locus (Supplementary Figure 19d).

While direct observation of TR LPS length variation yields interesting trends for pathogenic repeats, it ultimately fails to distinguish pathogenic loci from other highly polymorphic (yet seemingly not biologically relevant) loci like the CODIS set. Both these groups of repeats fall primarily in the top 10% of most polymorphic repeats for length variation (Supplementary Figure 20a). However, when using the observed-to-expected ratio based on our model, this limitation is alleviated. CODIS loci are identified as under neutral selection, protein-coding TRs show less variance than expected, and pathogenic and recently evolved TRs exhibit significantly more variance than expected (OR = 9.6, 95% confidence interval = 5.0 to 18.5, Fisher’s Exact Test p = 2.1e-11 for pathogenic loci; OR = 6.2, 95% confidence interval = 5.1 to 7.7, p = 3.1 e-56 for human ab-initio loci) (Supplementary Figure 20b).

These observations demonstrate the utility of this length constraint measure and highlight a crucial distinction between TR constraint and genic constraint. With TR constraint, both tails of the distribution are biologically relevant, whereas for genic constraint, solely the loci whose variation is selected against are medically significant. Consequently, we observe that TRs at protein-coding loci are largely on the lower end of the observed length variation distribution (Supplementary Figure 20a) and appear to be mostly under at least some degree of constraint (Supplementary Figure 20b).

Similar to the CODIS loci, the set of gene expression-associated STR (eSTR) loci identified by Fotsing and colleagues show generally high levels of LPS variation (Supplementary Figure 20a), consistent with their ability to affect gene expression. Interestingly, the constraint metric indicates that they have slightly more variance than expected with most entries falling in the top 30% of length constraint values (Supplementary Figure 20b).

We attempted to perform several orthogonal validations of the utility of this measure. First, we observed that the logarithm of TR length constraint is moderately correlated (Pearson correlation of 0.41) with the average age of onset among a set of 46 STR-associated diseases (Supplementary Figure 20c).

Second, we compared predictions by the TR Constraint model to five alternative approaches: the STR Constraint measure introduced by Gymrek and colleagues (referred to as “Gymrek Constraint”), the genic loss-of-function constraint score from gnomAD v4.0, the genic missense constraint score from gnomAD v4.0, the reference span of the STRs to serve as a naïve baseline, and a shuffled set of TR-Constraint predictions to serve as another naïve baseline. We also included the LPS length standard deviation generated in this work (referred to as “Observed Variance”) as another comparison point. We compared these seven values for their ability to prioritize disease-associated STRs, protein-coding STRs, and eSTRs (Supplementary Figure 21a-c). Our expectation was that disease-associated STRs should be prioritized at the extremes of the spectrum (most with high scores, but the poly-alanine and single-shift loci at the other end of the spectrum). We observed this pattern with TR Constraint, but the other approaches merely showed enrichment at the higher end of the spectrum (Supplementary Figure 21a). Next, we knew that protein-coding STRs should be constrained against variation, so we expected scores to be biased toward the lower end of the spectrum. We saw that Observed Variance fit that pattern very well, while TR Constraint and Gymrek Constraint showed a more muted effect of protein-coding STRs being slightly, but not overwhelmingly constrained (Supplementary Figure 21b). Finally, Observed Variance, TR Constraint, Gymrek Constraint, and Reference Span all showed a strong bias among the eSTR loci for higher scores (high variation, long reference span, or more variation than expected) with observed variance and reference span showing the strongest effects (Supplementary Figure 21c). Overall, these results indicate that TR Constraint performs broadly consistently with the established technique termed Gymrek Constraint, while being able to operate on a much wider range of STR loci. We also found that the reference span serves as a strong baseline. Surprisingly, the TR Constraint method did not clearly outperform direct usage of the LPS length variation measure in these tests.

A third orthogonal test was conducted using data from the recent article by Yoo, et al^43^ which introduced T2T reference genomes for six ape species identified 3,268 “Human Ancestor Quickly Evolved Regions” (HAQERs) which represent the regions that diverged the most between humans and other ape species. Since we suspected that our TR constraint measure may be related to positive and negative selective pressure, we thought this set of loci could help test that notion. We found that both the direct measurement of LPS length variation (Supplementary Figure 22a) and TR Length Constraint (Supplementary Figure 22b) showed enrichment for HAQER loci among both their highest and lowest deciles of loci. This finding reinforces the notion that this measure is successfully identifying loci under active selection and prioritizing them at the two tails of the TR Constraint distribution, regardless of whether they be under positive or negative selection.

This predictive model of TR variability can also be used to perform counterfactual analyses interrogating which aspects of a nucleotide sequence result in changes in the predicted TR variability. This enables in silico experiments such as investigation of which motifs and motif lengths are most conducive to greater TR variability, how flanking sequence characteristics influence TR variability, and determining the precise interplay in effect between LPS length and total TR length with interruptions as they influence TR variability.

#### **Long-read vs Short-read in 689 shared samples**

##### *ExpansionHunter*

Many studies have investigated TR lengths and variation using large cohorts of short-read data. Many approaches exist to profile TRs in short-read data and we will not attempt a comprehensive comparison of such methods in this work, but instead will just use one commonly used method called ExpansionHunter (EH). Previous work has demonstrated EH to be very accurate at predicting tandem repeats shorter than the read length and it typically gives broadening confidence intervals when attempting to estimate increasingly larger allele sizes beyond the read length^6^. Since we had access to 30x Illumina WGS data on 689 of the individuals in the validation cohort, we sought to compare EH performance with that of TRGT to determine at which size threshold it becomes less accurate and test whether motif variation exacerbates that effect.

For TR alleles that TRGT measures to be shorter than 150 bp, EH makes predictions that are highly consistent with TRGT: achieving an overall Pearson R^2^ value of 0.977. However, for alleles that TRGT judges to be longer than 150 bp, this relationship is greatly diminished (Supplementary Figure 23A). The Pearson R^2^ value drops to only 0.096. Many alleles are genotyped with reasonable accuracy by EH, but the allele lengths EH is willing to predict clearly plateau around 500 bp, while TRGT does not (Supplementary Figure 23A). Dissecting the sizing discrepancies between the tools further, we saw that the vast majority of alleles estimated to be shorter than 150 bp by TRGT agreed with their EH estimate within 1 bp (Supplementary Figure 23B), but the estimates diverged for longer alleles. For alleles over 250 bp, they most often differed by more than 16 bp. This gradual shift in the magnitude of disagreement between EH and TRGT as alleles lengthen illustrates the challenges of this work with short-read data and the advantages of using a high quality long-read dataset such as ours. There is a small set of alleles in Supplementary Figure 23A on the upper-left corner of the plot which TRGT calls as being over 500 bp, but EH calls as being under 100 bp. As best we can tell, these are sequencing errors in the HiFi data which produce long homopolymer stretches which we are not convinced are real. Further investigation of such events is necessary.

For the TRGT calls over 150 bp, we also plotted the absolute difference between the TRGT allele length and the corresponding EH allele length stratified by TR motif (Supplementary Figure 23C). This view shows that the AAAAT motifs represent some of the most dramatically mis-genotyped loci by EH. We also observe numerous genotyping discrepancies at known pathogenic loci that constitute shifts of several repeat units, which could prove problematic in medical genetics work using EH as a screening tool.

##### *ExpansionHunter Denovo*

In large cohorts sequenced by short-read approaches, the ExpansionHunter Denovo (EHDn) tool is often used to identify TRs that are expanded in one or several individuals in pursuit of novel pathogenic repeat expansions^44^. Hence, we investigated the accuracy of identifying large repeats using EHDn. We applied EHDn to 689 short-read genomes, and TRGT to their corresponding long-read counterparts. For this analysis, we calculated precision and recall values at different comparison thresholds.

EHDn does not estimate repeat size, but instead provides a measure of the number of anchored in-repeat-reads (IRRs) detected at a particular locus. High anchored IRR counts at TRs indicate the repeat is larger than the read length (>175 bp). Using TRGT genotypes as the ‘truth set’, we calculated precision and recall values for different thresholds of EHDn anchored IRRs and TRGT repeat sizes. We found that a measure of five anchored IRRs produced a median precision of 87.83% and recall of 1.02% for identifying repeats of 125 bp or larger among all samples. As we increased the size threshold in increments of 25 bp up to 225 bp, the median precision decreased to 66.41% and the recall increased to 2.88% (Supplementary Figure 24A, C). We also calculated precision and recall values for different anchored IRR thresholds, to measure their ability to identify repeats greater than 175 bp. We found that 5 anchored IRRs had a median precision of 80.95% and recall of 1.94%, and the precision increased with an increase in anchored IRRs, but the recall decreased (Supplementary Figure 24B, D).

While exploring the EHDn TRs, we found a subset that were not genotyped by TRGT that we have labelled as *de novo* TRs. We found that *de novo* TRs comprised 29.94 – 52.88% of all EHDn-identified TRs when filtering for those with anchored IRRs above five. As we increased the threshold to 15 anchored IRRs, the upper range of *de novo* TRs increases to 66.67%, with a median of 45.45% (Supplementary Figure 24E).

Next, we combined all the TRs with five or more anchored IRRs from each sample and removed duplicate loci within each group. Supplementary Figure 25A shows the distribution of TRs that were expanded, not expanded, had a motif mismatch, or were de novo. We then compared all the cataloged TRs to the *de novo* TRs. We found that *de novo* TRs were more likely to be defined as satellites (odds ratio [OR] = 17.09; 95% confidence interval [CI] = [5.92, 49.33]), and less likely to be defined as simple repeats (OR = 0.5; 95% CI = [0.43, 0.59]) or SINEs (OR = 0.70; 95% CI = [0.61, 0.79]) according to the UCSC repeat database. The majority of these *de novo* TRs were not categorized in this database (Supplementary Figures 25B, C). Upon investigating motif patterns, we found that catalogued TRs were enriched in AAAG (OR = 6.19; 95% CI = [4.39, 8.72]) and AAGGAG (OR = 5.02; 95% CI = [2.33, 10.80]) motifs while *de novo* TRs were enriched in AATGG (OR = 42.41; 95% CI = [18.67, 96.34]) and AACCCT (OR = 4.79; 95% CI = [2.15, 10.67]) motifs (Supplementary Figure 25D, E). We also discovered that *de novo* TRs were enriched in A+G+T combinations (OR = 5.13; 95% CI = [3.76, 7.00]) in their motifs, as well as A+C combinations (OR = 1.64; 95% CI = [1.39, 2.39]) (Supplementary Figure 26A, B).

We also drew comparisons between TRs that were recalled and those that were not. To do this, we combined all the TRGT genotyped TRs for each sample and categorized them into those that were recalled by EHDn, and those that were not. We found that unrecalled TRs were most enriched in A+T (OR = 11.76; 95% CI = [11.30, 12.25]) or A+C (OR = 15.12; 95% CI = [15.12, 16.54]) nucleotide combinations (Supplementary Figure 26C-D). Upon comparing the GC content of TRs that were recalled and unrecalled, we found surprisingly that recalled TRs have a higher GC percentage in their motifs (Cohen’s d = 0.62), are less pure in their motifs (d = -0.59) and are larger in LPS size (d = 3.25) (Supplementary Figure 27A-C). We did not see any significant difference in the number of unique motifs observed at specific loci between the two groups (Supplementary Figure 27D).

### **SUPPLEMENTARY FIGURES**

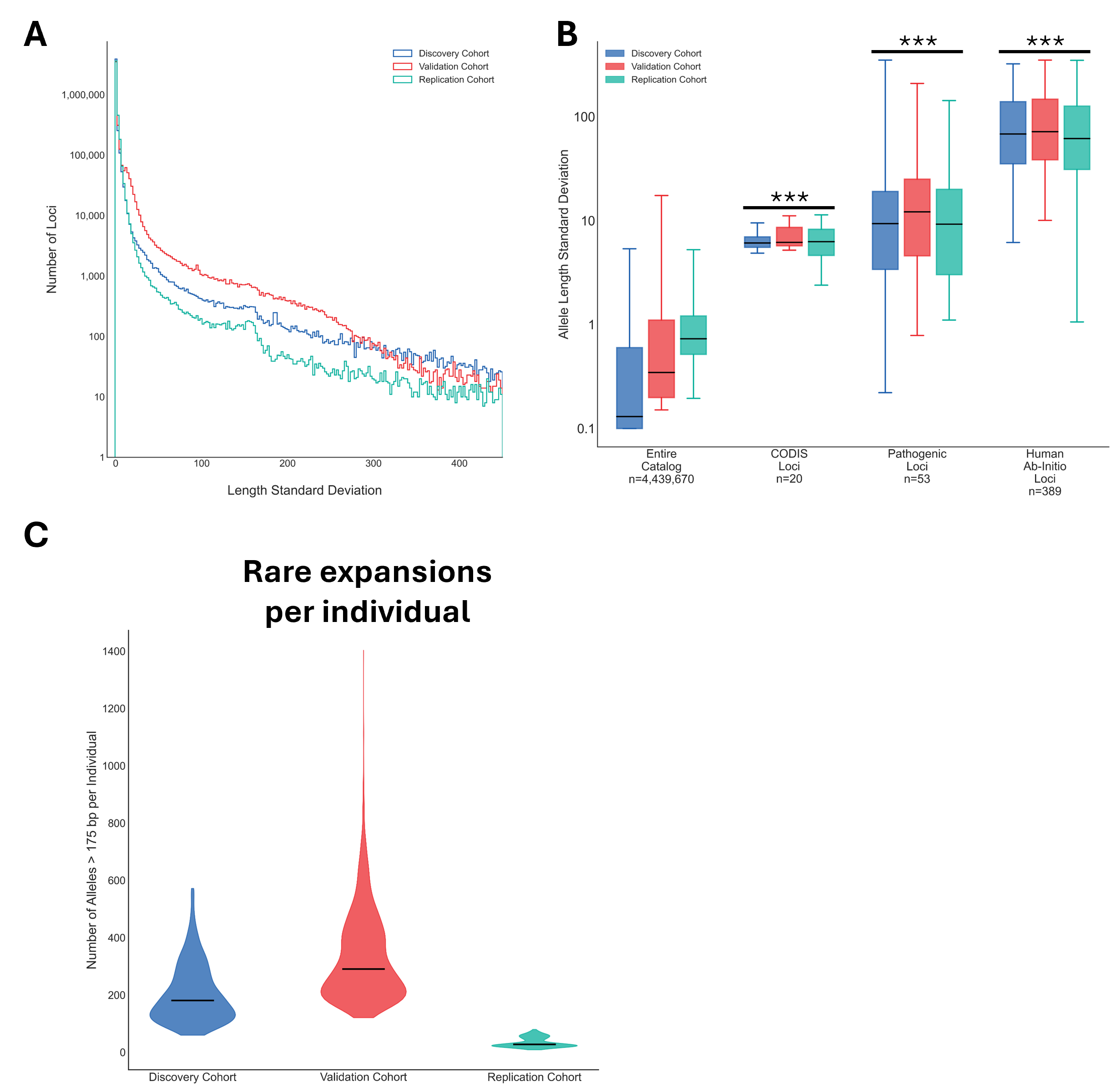

Supplementary Figure 1: Length variation in 23 billion tandem repeat alleles

A) Histogram of standard deviation of repeat length for each of the 4.4 million analyzed TR loci in each of the three cohorts. B) Boxplots of standard deviation of repeat length for the entire catalog, the known pathogenic loci, the set of loci identified by Sulovari and colleagues to be human-specific (human Ab-initio loci), and CODIS loci. *** indicates a p-value<10^-10^ by Mann-Whitney U-Test for each cohort subset relative to the complete catalog for that same cohort. C) Violin plots of the count of loci per individual that are longer than 175 bp in that person but are shorter than 175 bp in >99% of that cohort.

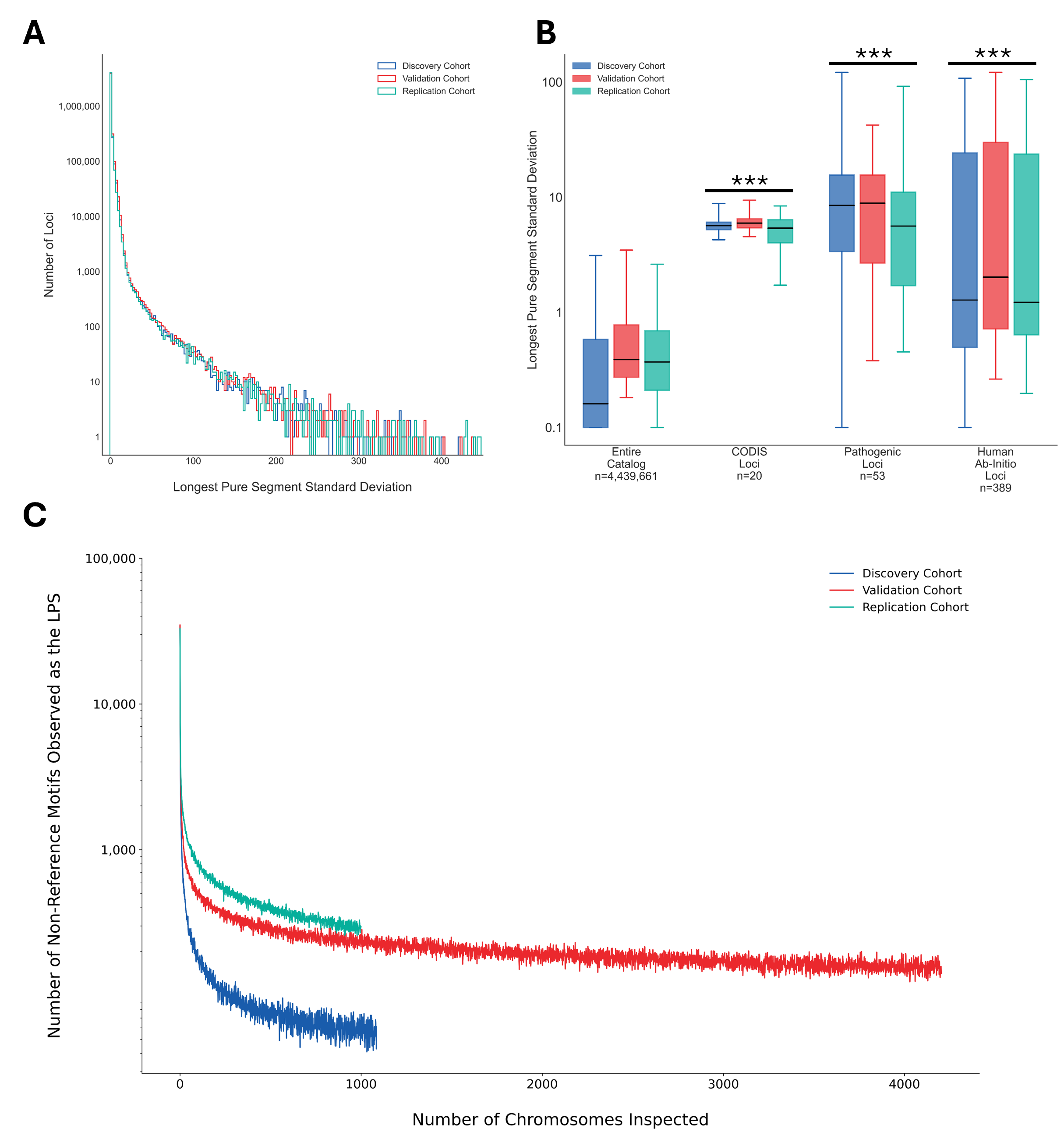

Supplementary Figure 2: LPS length variation overview

A) Histogram of standard deviation of longest pure segment (LPS) length for each of the 4.4 million analyzed TR loci in each of the three cohorts. B) Boxplots of standard deviation of LPS length for the entire catalog, the known pathogenic loci, the set of loci identified by Sulovari and colleagues to be human-specific (human Ab-initio loci), and CODIS loci. *** indicates a p-value<10^-10^ by Mann-Whitney U-Test for each cohort subset relative to the complete catalog for that same cohort. C) Discovery curve of the number of non-reference motifs observed as the LPS motif as individuals are added to each cohort.

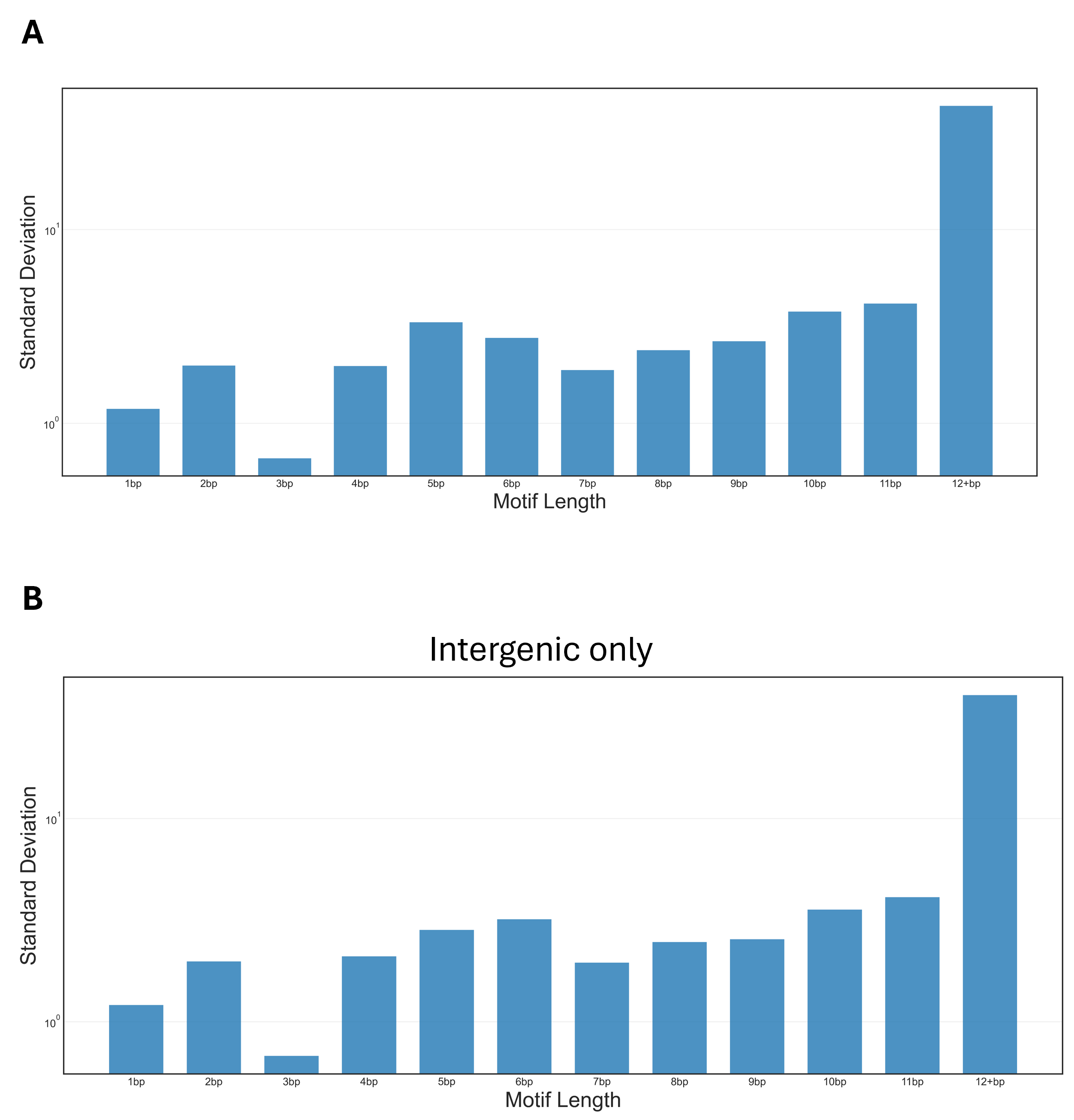

Supplementary Figure 3: LPS variation by motif length

Bar plots of the median standard deviation of LPS length for loci stratified by LPS motif length. A) genome-wide, and B) intergenic loci only.

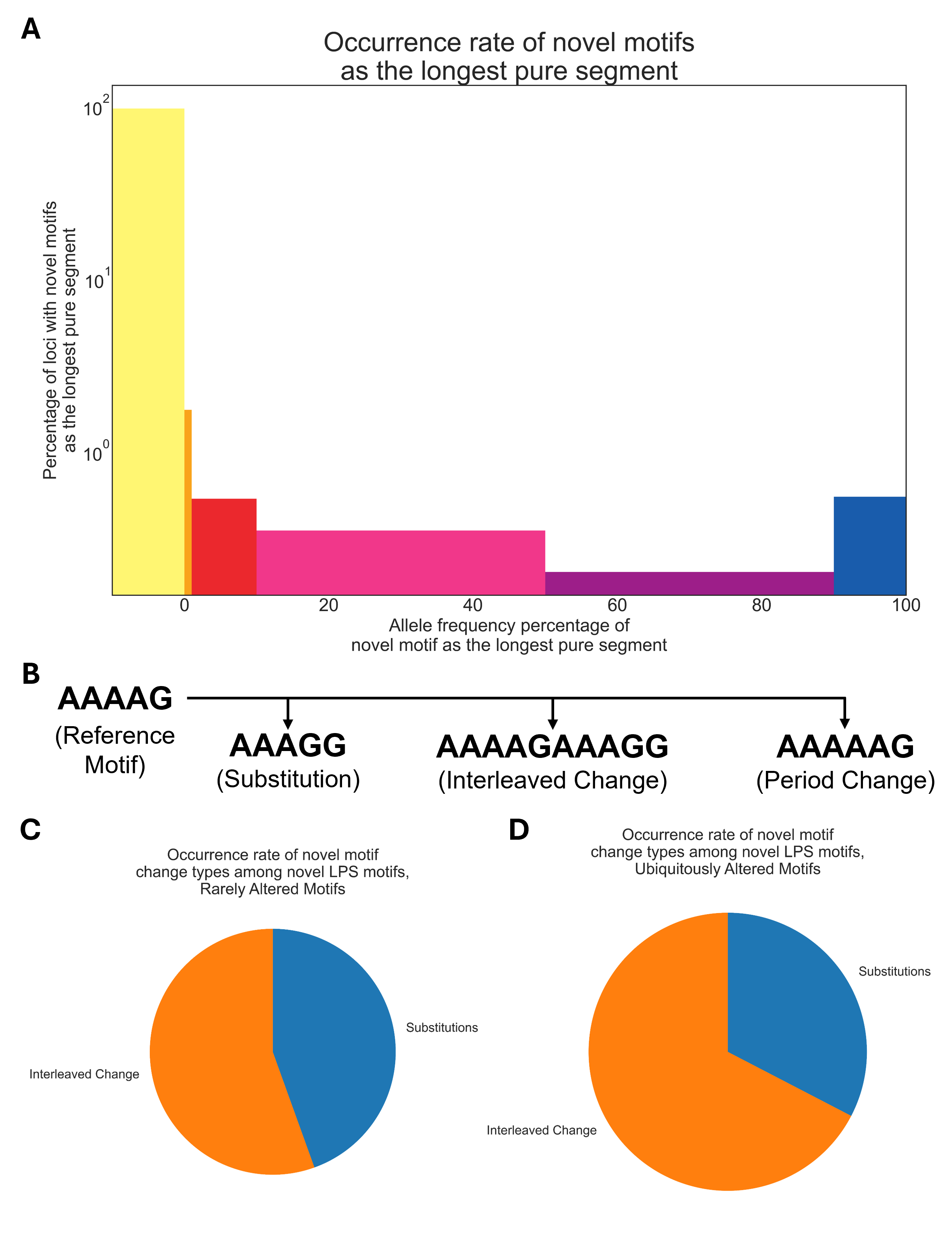
Supplementary Figure 4: Motif variation is a major source of polymorphism at tandem repeat loci

A) Variable-width bar plot of the proportion of loci found to harbor motifs not present in the reference catalog. To be counted, a motif must create the longest pure segment observed in an allele. B) Diagram of motif changes to a reference motif that depict it changing into various non-reference motifs which can be of the same period (first example), an interleaved pattern with a period that is a direct multiple of the reference motif (second example), and a non-reference motif period (third example). C and D) Pie charts of the occurrence rate of motif change type when the LPS is a novel motif among loci with C) rarely-altered (<1% of alleles non-reference motif) motifs and D) ubiquitously-altered (>99% of alleles non-reference motif) motifs.

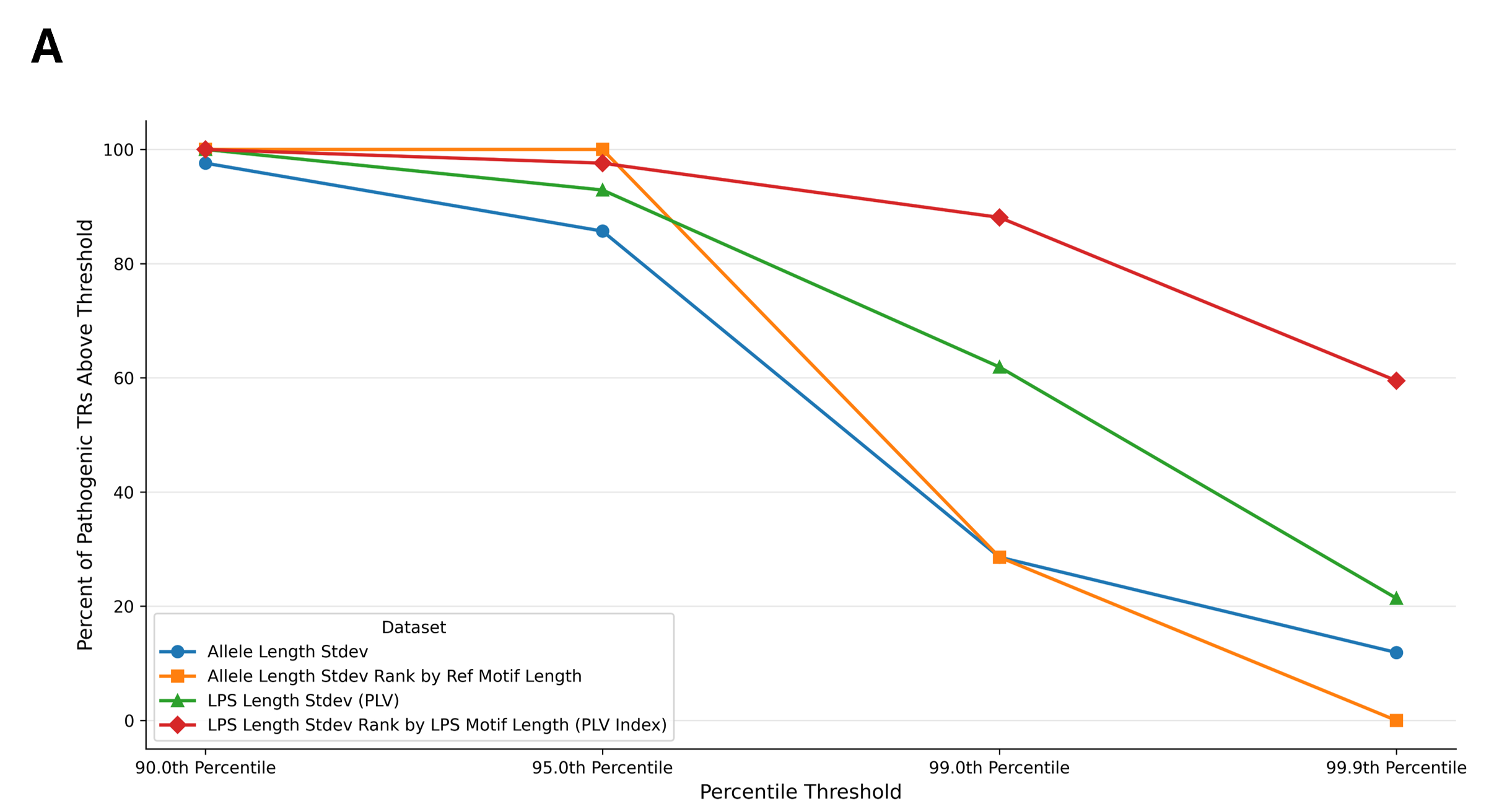

Supplementary Figure 5: Disease-associated STR enrichment by threshold and methodology

A) Line plot of the percentage of disease-associated STRs falling above each percentile threshold (x-axis) when the discovery cohort is analyzed by the standard deviation of either allele length (blue), allele length with reference motif length normalization (orange), LPS length (PLV) (green), or LPS length with LPS motif length normalization (PLV Index) (red).

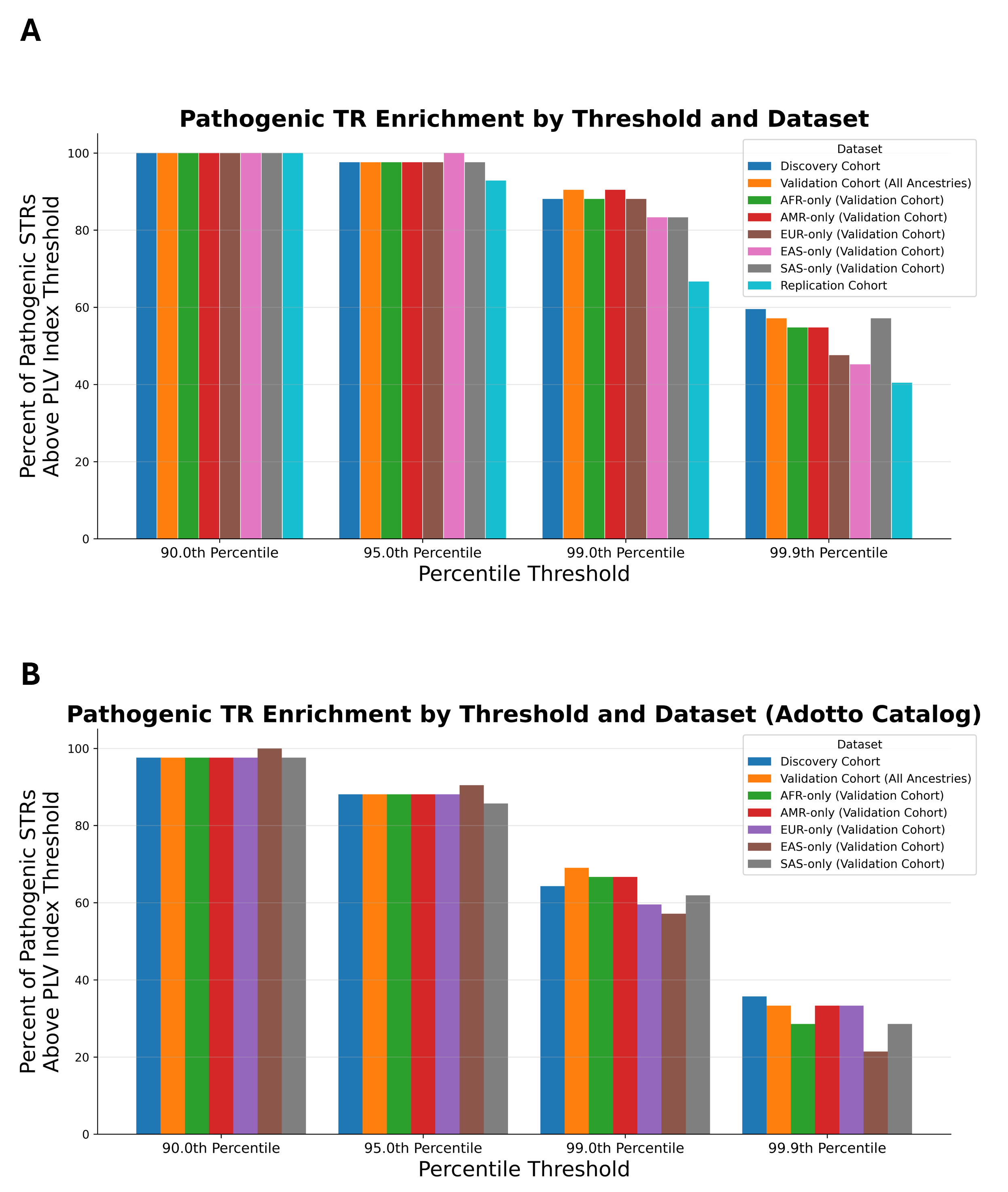

Supplementary Figure 6: Replication of disease-associated STR enrichment across cohorts, ancestries, and catalogs

A-B) Bar plots of the replication of the proportions of disease-associated STR loci among the top N percentile groups across the three cohorts and the genetic ancestry groups within the validation cohort. Each bar shows the percentage of disease-associated STR loci that ranked among the top N percentile in that dataset when using LPS length variation normalized by LPS motif length (PLV Index). The values of N used are the 90^th^, 95^th^, 99^th^, and 99.9^th^ percentiles. A) TR-Explorer catalog. B) Adotto catalog.

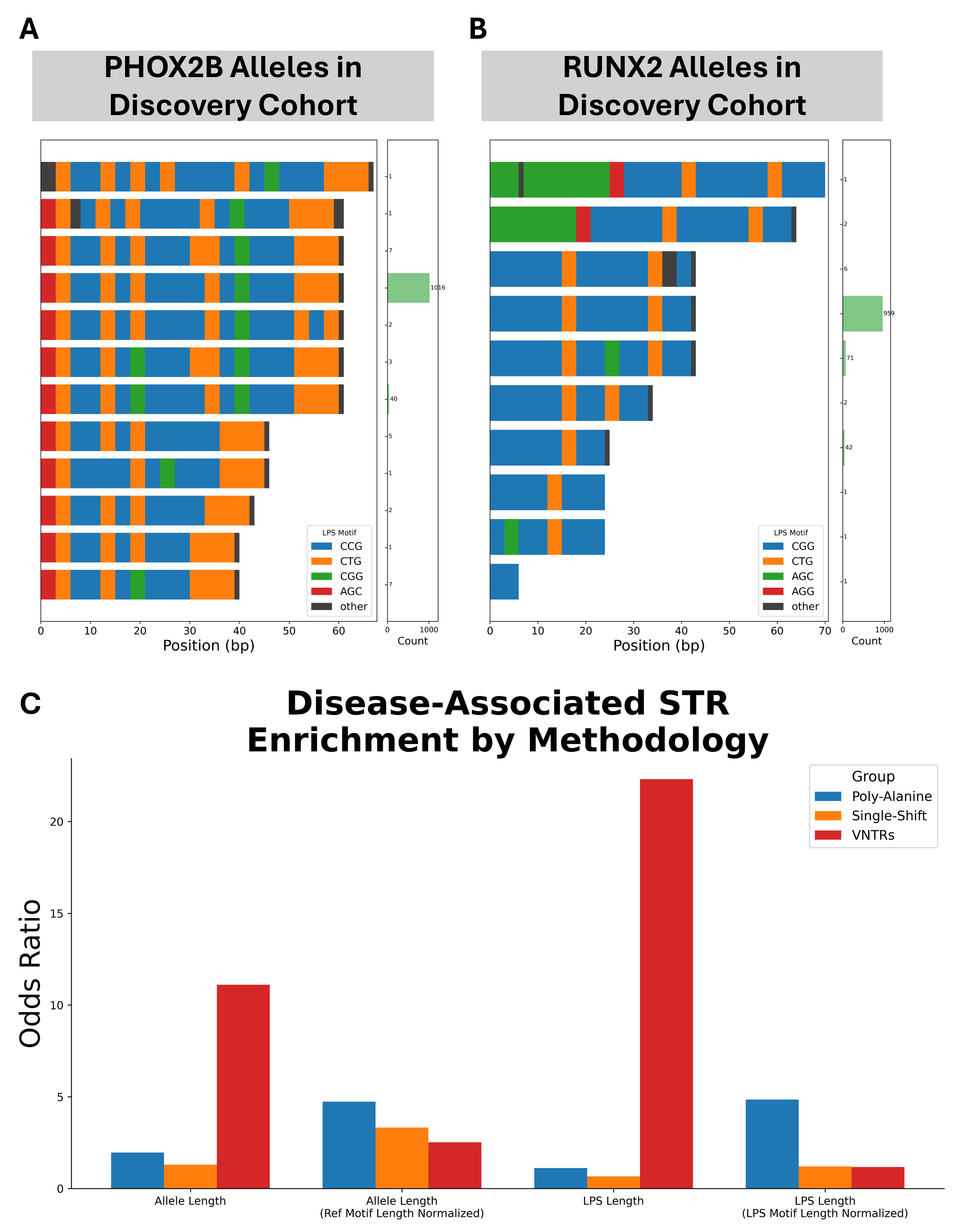

Supplementary Figure 7: Limitations of disease-associated TR enrichment with population variation

Waterfall plots of A) PHOX2B and B) RUNX2 poly-alanine disease-associated loci showcase how their GCN repeats pose a challenge for LPS-based analyses. C) Bar plot of the generalized odds ratio of the enrichment of disease-associated poly-alanine, single-shift, and VNTR loci among the most polymorphic loci in the catalog using overall allele length without normalization (first bar), allele length normalized by reference motif length (second bar), LPS length without normalization (PLV) (third bar), and LPS length normalized by LPS motif length (PLV Index) (fourth bar).

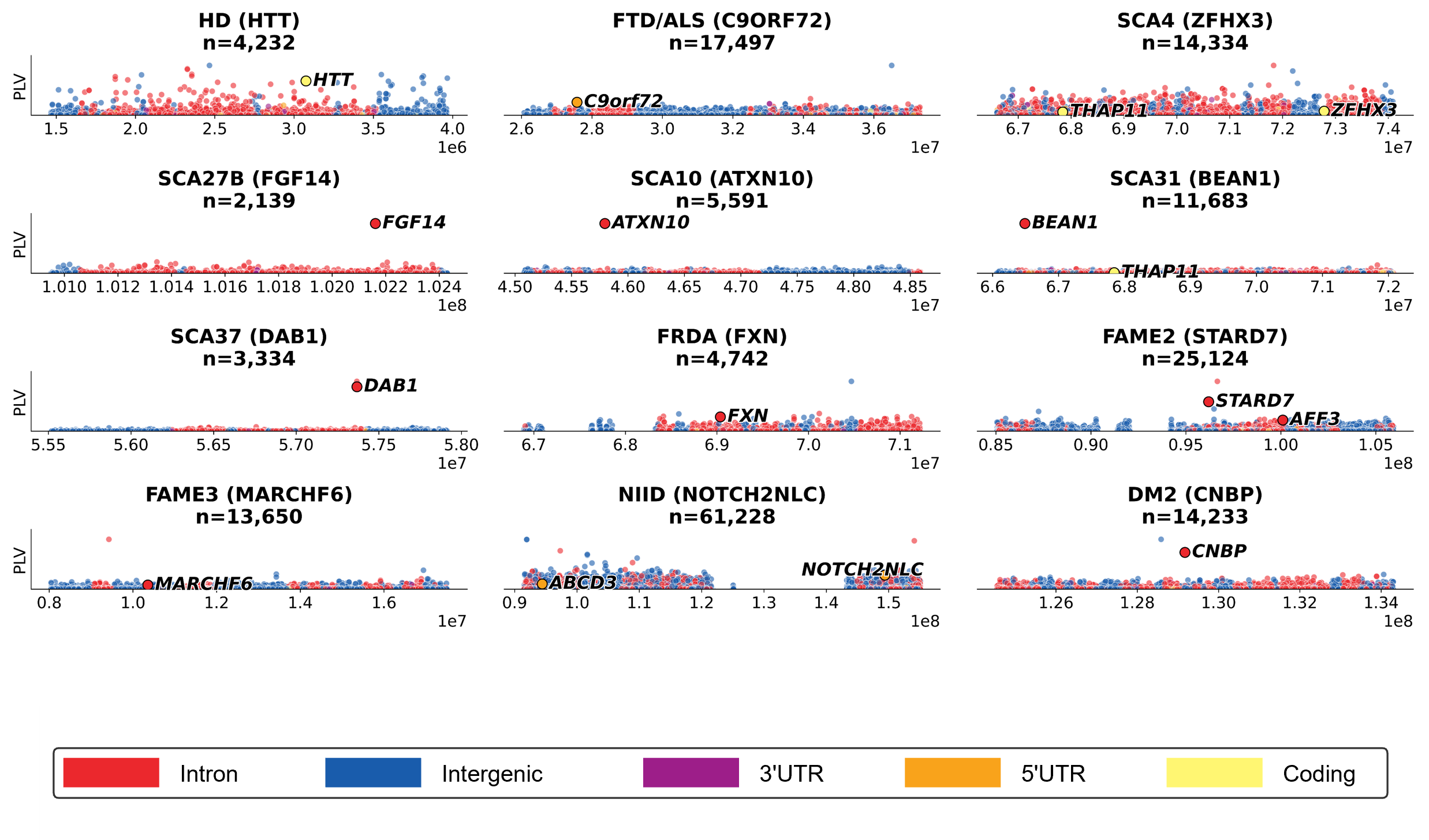

Supplementary Figure 8: Additional linkage regions

Series of Mahattan plots of STRs across 12 published linkage regions. In each panel, the x-axis is the genomic coordinates across the linkage region span and the y-axis is the LPS length standard deviation of each TR using its most common LPS motif. Known pathogenic TRs are specifically labelled in each plot.

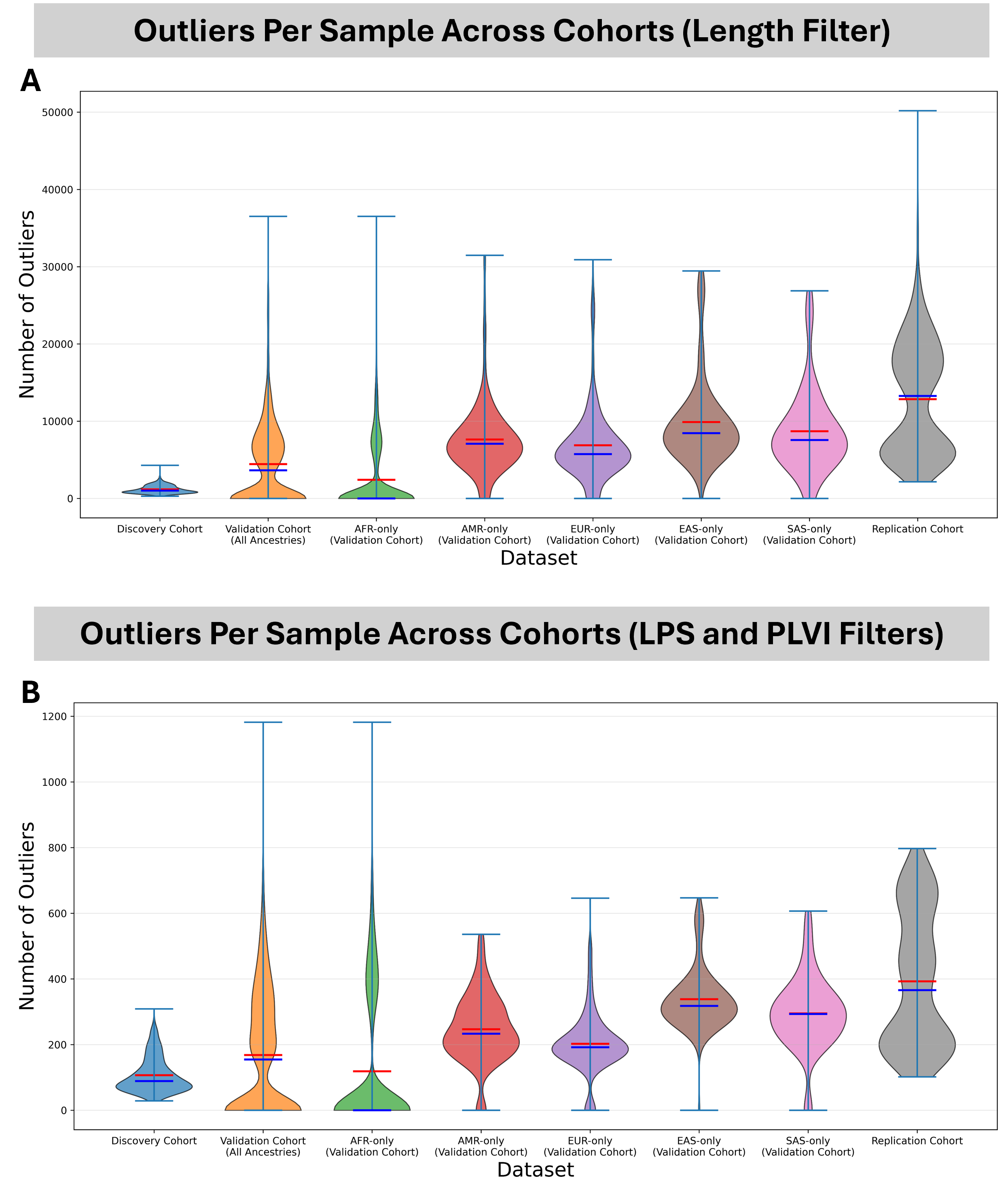

Supplementary Figure 9: Outliers per sample

Violin plots of the number of “outlier” STR loci observed per individual within each cohort or cohort set. In A) an “outlier” is defined as an STR whose allele length exceeds the 99.9^th^ percentile for that locus in the discovery cohort. In B) an “outlier” is defined as an STR above the 95^th^ percentile of LPS variability (normalized by LPS motif length) whose LPS length exceeds the 99.9^th^ percentile for that locus in the discovery cohort for that locus with the same LPS motif.

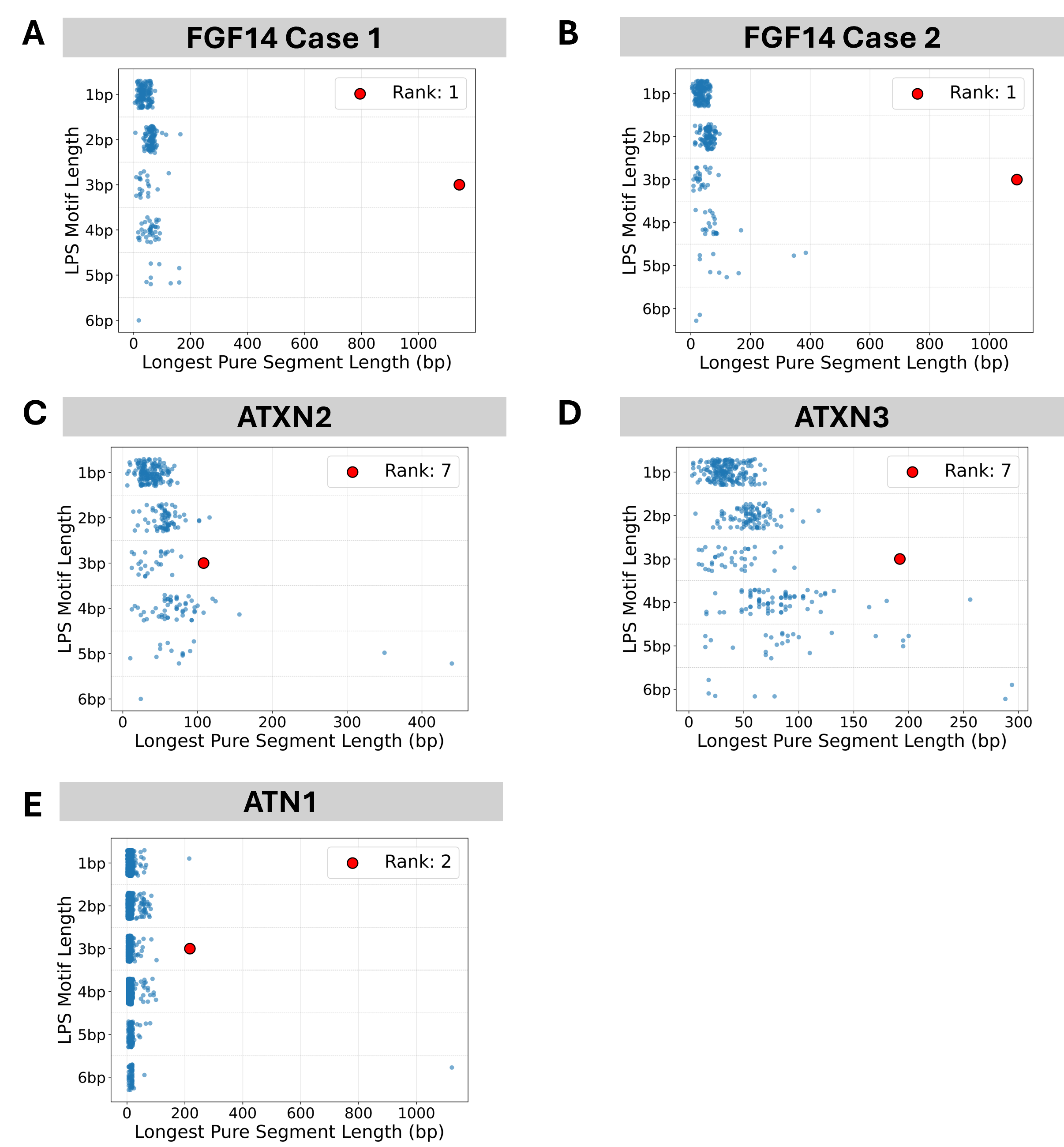

Supplementary Figure 10: Prioritization of pathogenic TR expansions in individuals

A-E) Swimlane plots showing the prioritization of pathogenic alleles in four individuals: A) FGF14 Case 1, B) FGF14 Case 2, C) ATXN2, D) ATXN3, and E) ATN1.

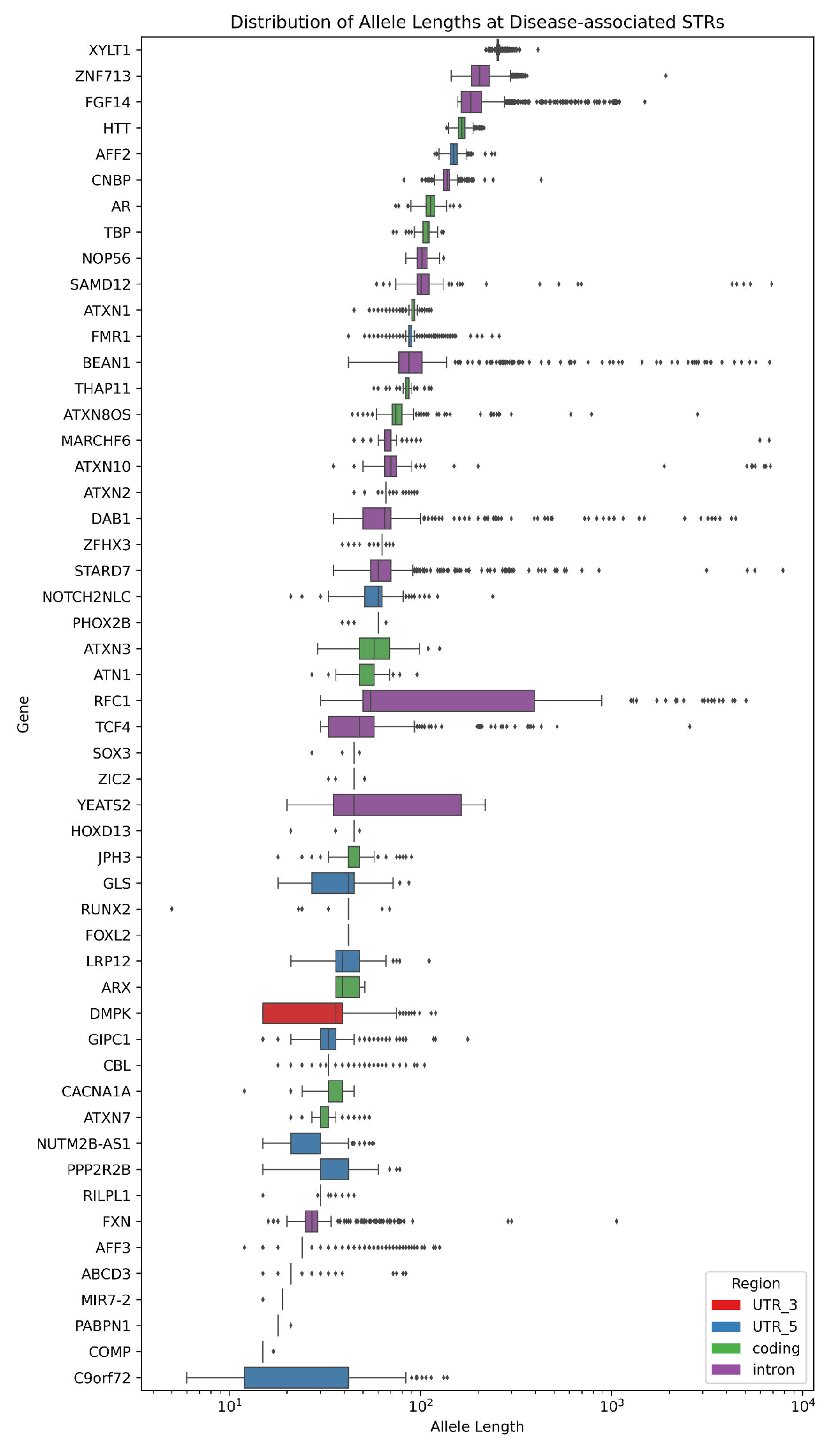

Supplementary Figure 11: Characterization of variation observed at disease-associated STRs

Boxplots of repeat length (base pairs) of disease-associated STR loci in the 1,086 haplotypes of the discovery cohort.

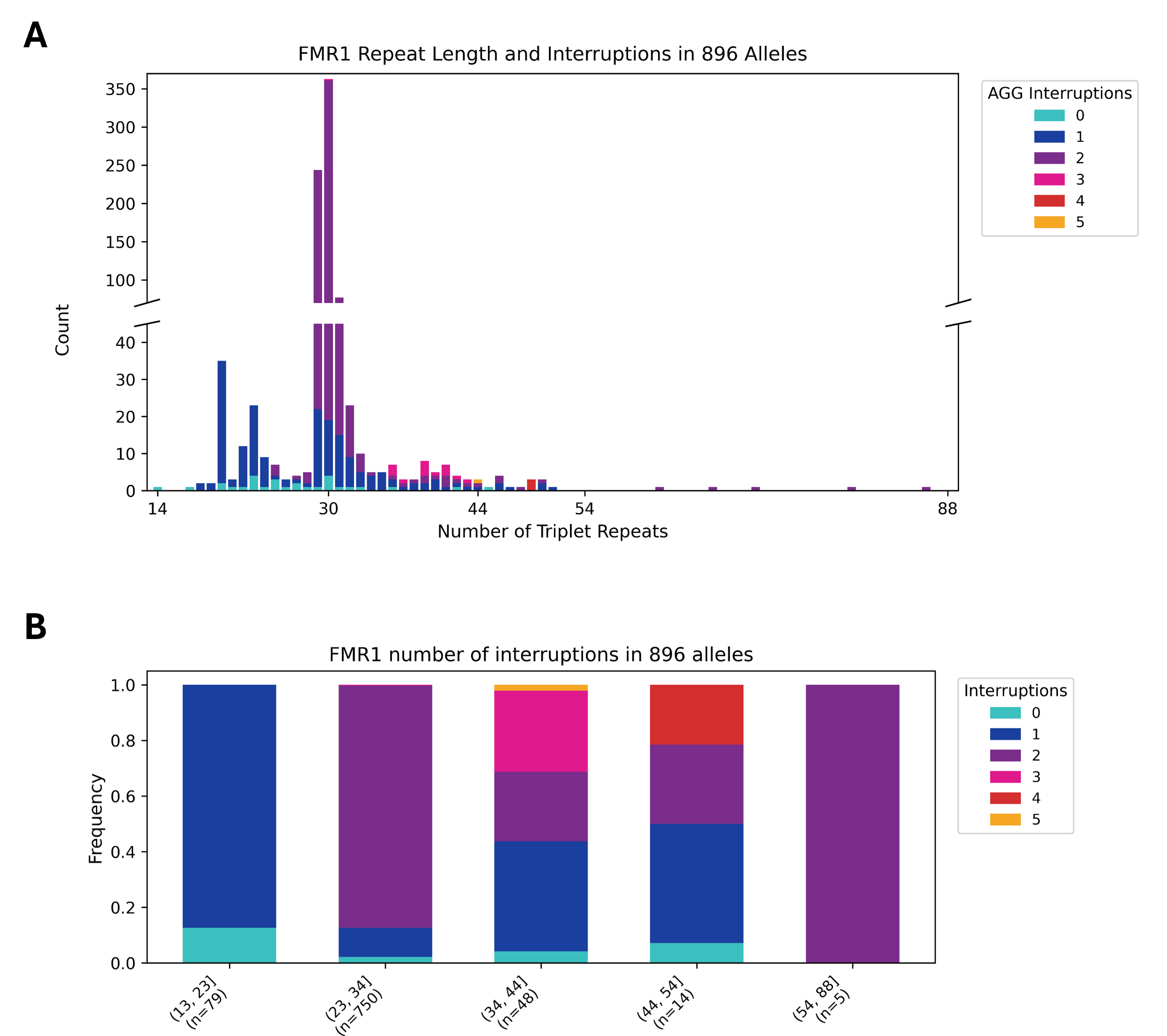

Supplementary Figure 12: Length variation and AGG interruptions in *FMR1*

A) Histogram depicting the frequency of repeat lengths and AGG interruptions in the *FMR1* repeat locus, n=896. B) Stacked bar plot showing the proportion of different amounts of AGG interruption in *FMR1* alleles for five different ranges of total repeat length.

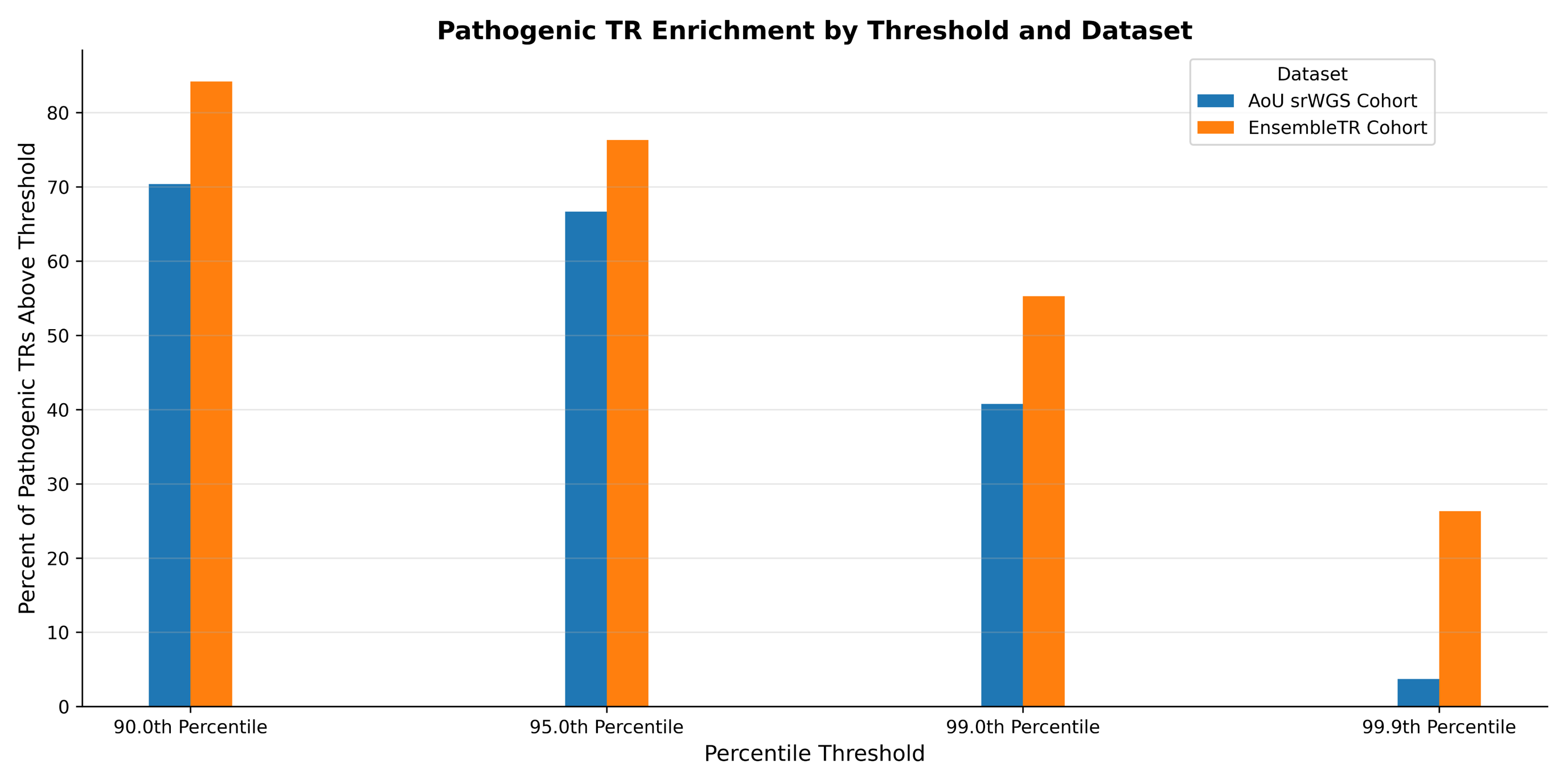

Supplementary Figure 13: enrichment of disease-associated STRs among most polymorphic loci in srWGS data

Grouped bar plot of the percentage of disease-associated STRs among the top N percentile groups across two srWGS cohorts. Each bar shows the percentage of disease-associated STR loci that ranked among the top N percentile in that dataset when using allele length variation normalized by reference motif length. The values of N used are the 90^th^, 95^th^, 99^th^, and 99.9^th^ percentiles. The number of disease-associated STR loci was 27 in the AoU srWGS cohort and 38 in the EnsembleTR cohort. In both cases, the loci used are a subset of those used in Figure 3 and Supplementary Figure 6.

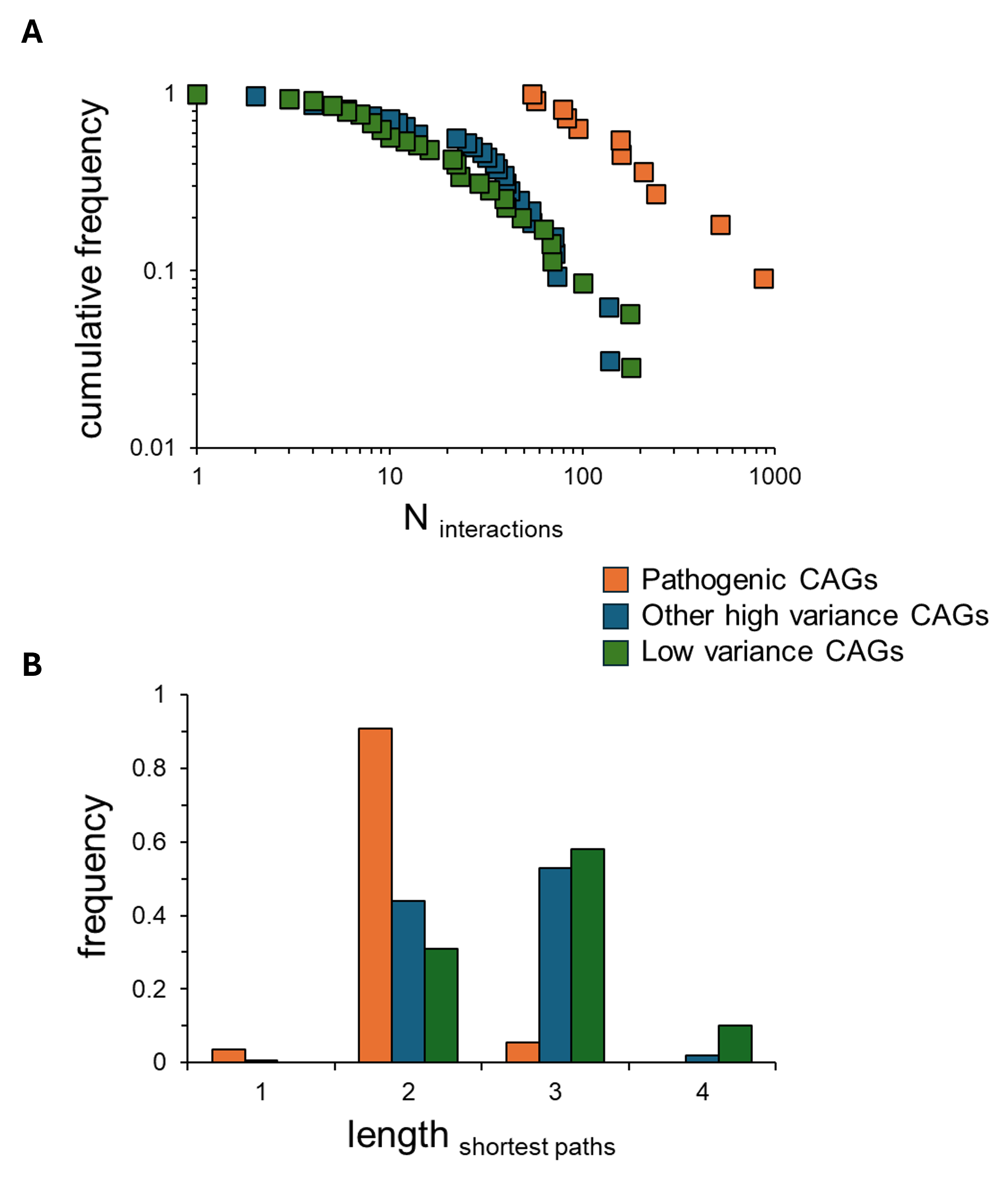

Supplementary Figure 14: Network analysis of genes with the most and least polymorphic CAG loci

A) Cumulative frequency plot of the number of protein-protein interactions for the genes containing either the known pathogenic coding CAG TRs (orange), the other high variance coding CAGs identified in Figure 2E (blue), or a set of low variance coding CAGs. B) Distribution of shortest path lengths between members of the gene sets in panel A across the protein-protein interaction network.

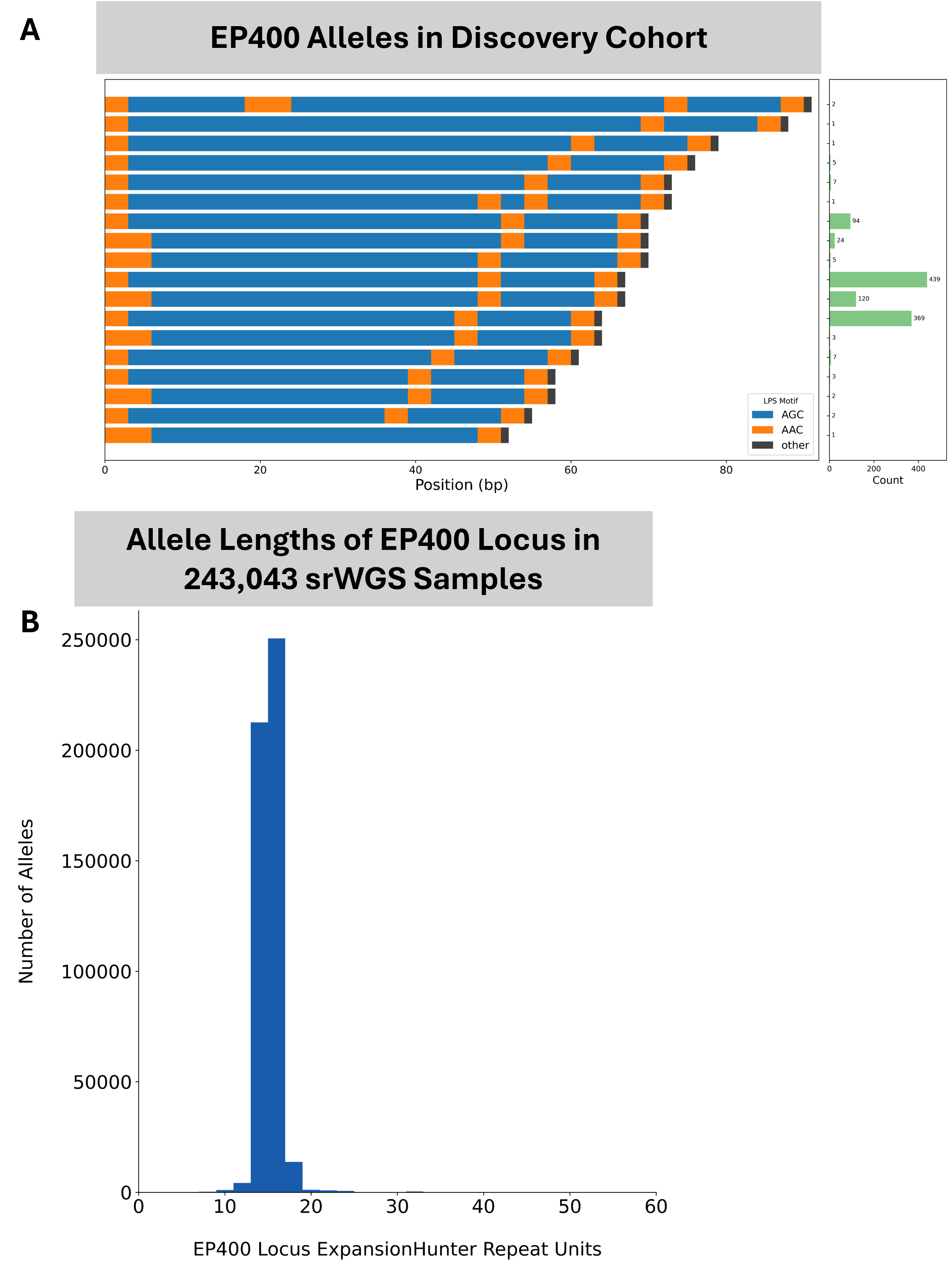

Supplementary Figure 15: *EP400* population statistics

A) Waterfall plot of unique repeat alleles for *EP400* in the discovery cohort. The green bars on the right side of each plot reflect the number of times each allele was observed in the cohort. B) allele length distribution in 243,043 srWGS general population samples.

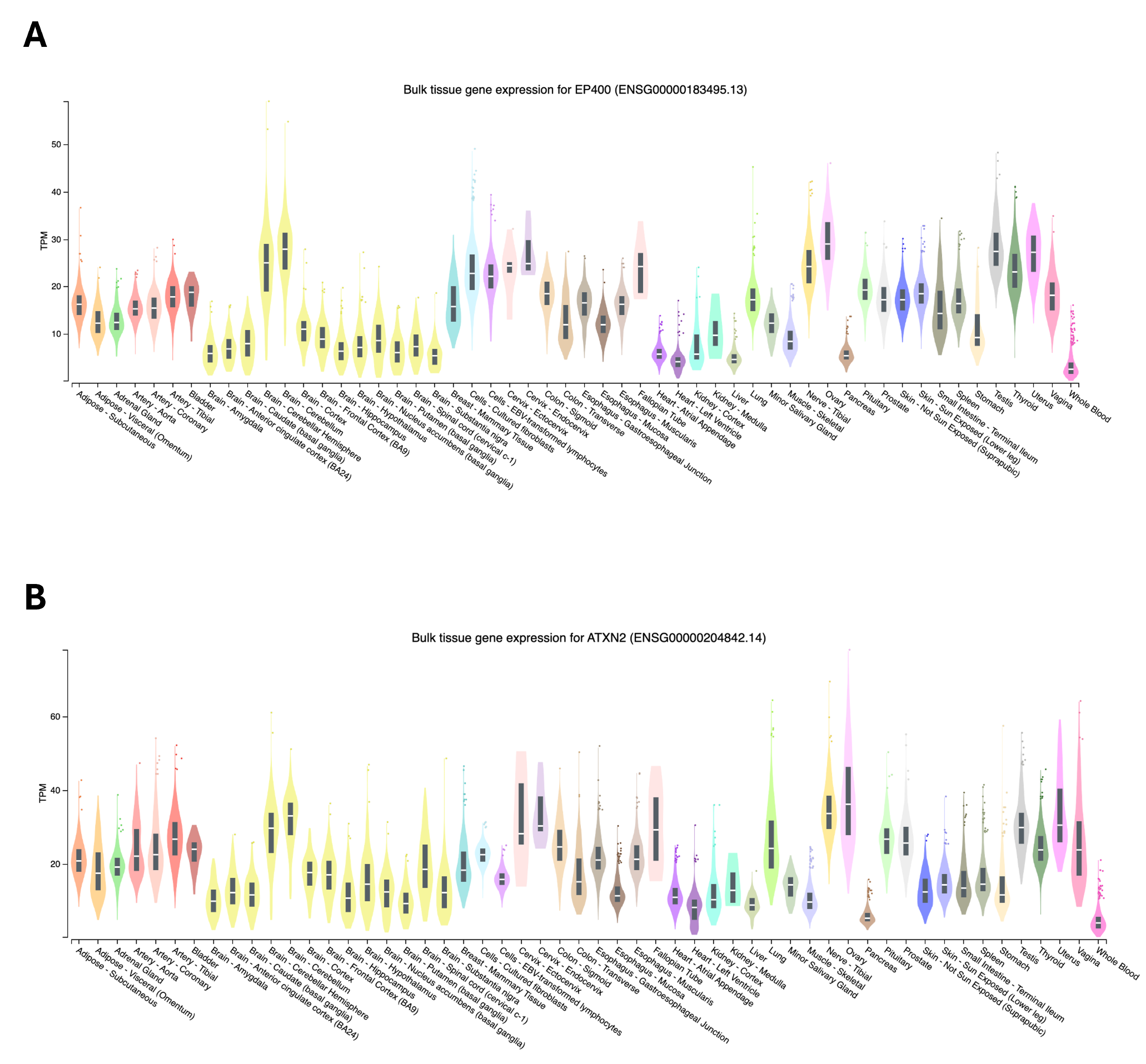

Supplementary Figure 16: Expression variation of *EP400* and *ATXN2*

A) Gene expression levels of *EP400* across many tissues as reported by GTEx. B) Gene expression levels of *ATXN2* across many tissues as reported by GTEx.

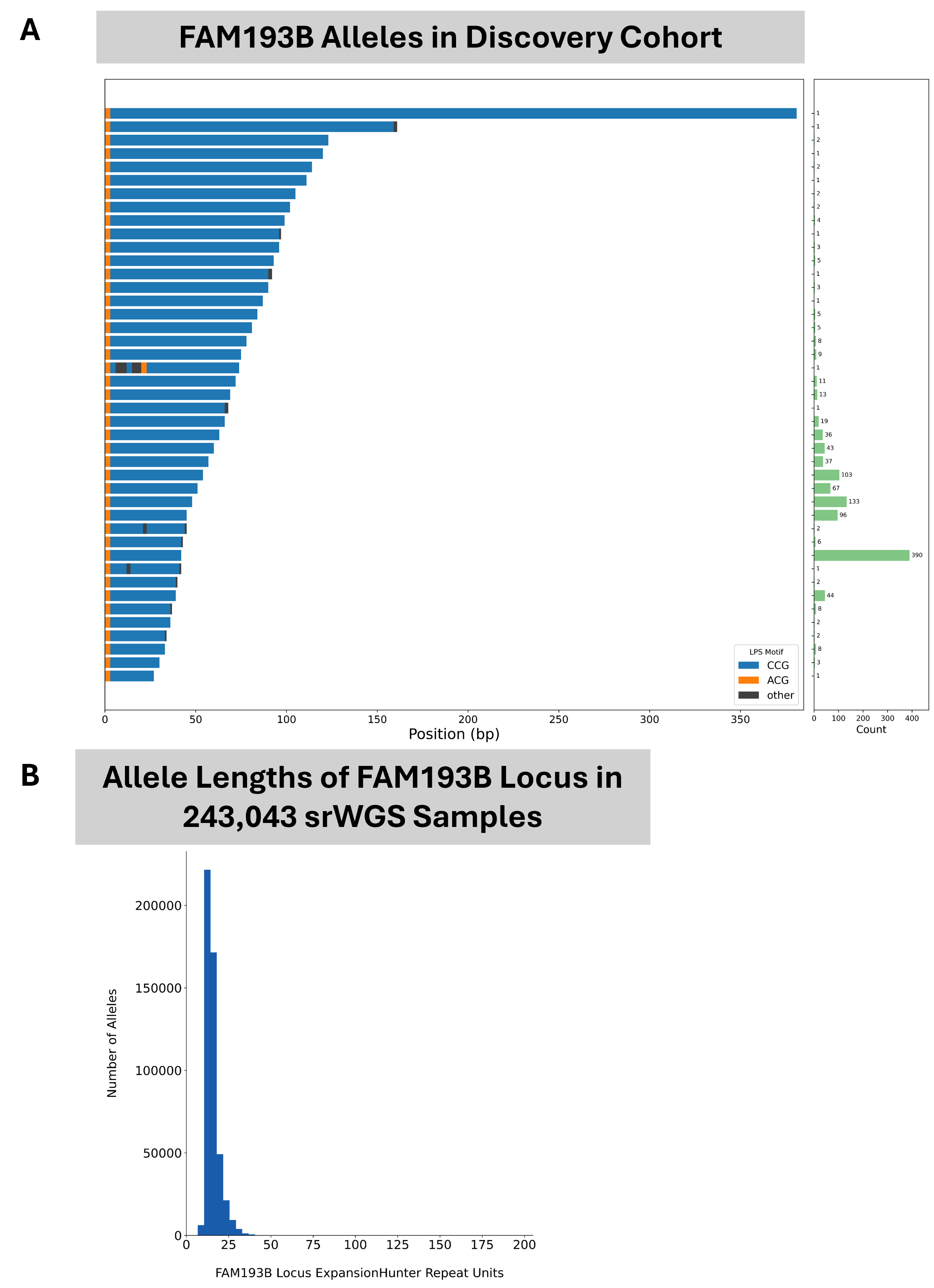

Supplementary Figure 17: *FAM193B* population statistics

A) Waterfall plot of unique repeat alleles for *FAM193B* in the discovery cohort. The green bars on the right side of each plot reflect the number of times each allele was observed in the cohort. B) allele length distribution in 243,043 srWGS general population samples.

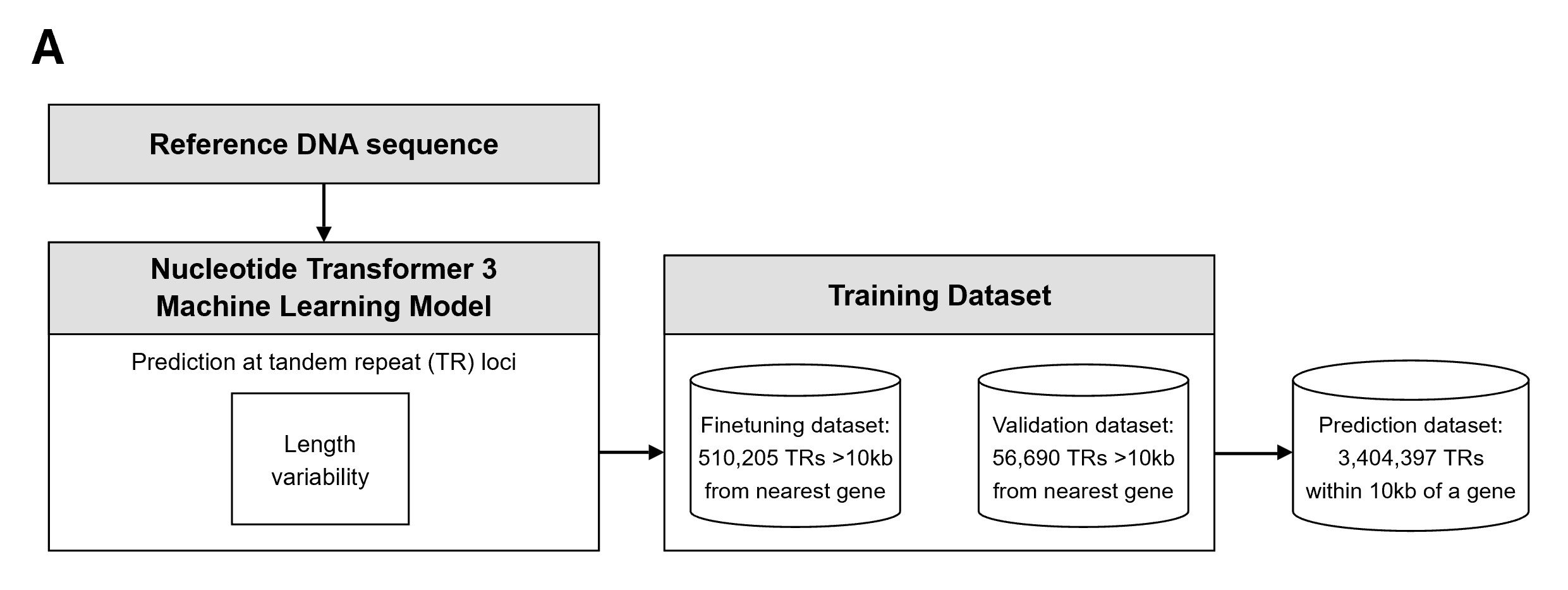

Supplementary Figure 18: TR Constraint training and model overview

Schematic of the deep learning system to predict the length variation of TRs based only on their reference DNA sequence by fine-tuning a 650 million parameter Nucleotide Transformer 3 model. The three datasets used to fine-tune, validate, and evaluate the constraint model are also described.

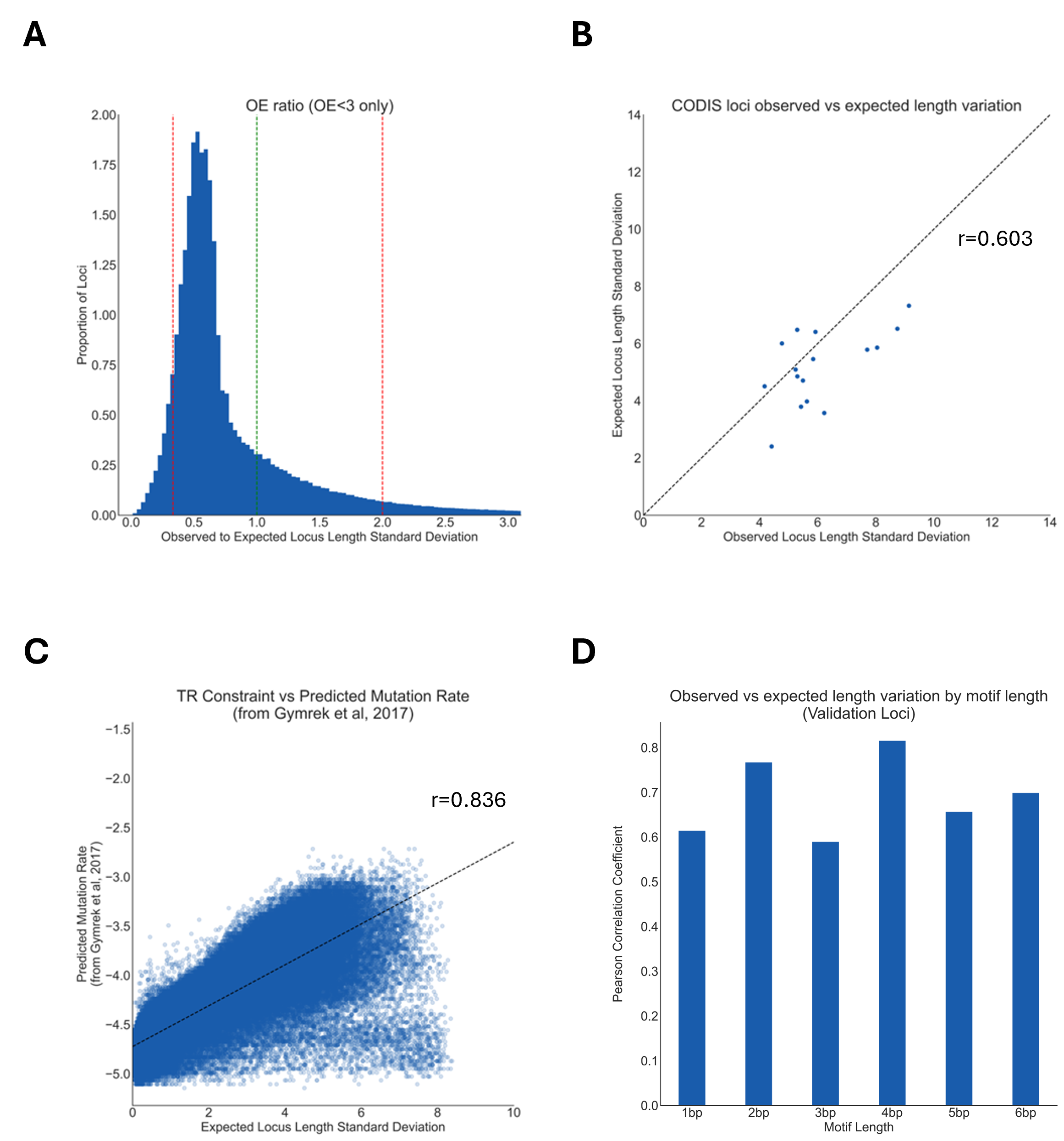
Supplementary Figure 19: Technical characteristics of TR length constraint

A) Histogram of observed-to-expected ratio for length variation for loci within 10kb of a gene. This plot only shows values below 3 for visual clarity. Red vertical lines mark values of 0.333 (representing the threshold of negative selection) and 2.0 (representing the threshold of positive selection). The green vertical line marks the value of 1.0. B) Scatterplot of the observed vs expected length variation for the 16 CODIS loci within 10kb of a gene. The Pearson correlation coefficient for this set of values is 0.603. C) Scatterplot of the predicted mutation rate from Gymrek, et al 2017 vs expected length variation from this model. The Pearson correlation coefficient for this set of values is 0.836. D) Bar plot of the Pearson correlation coefficient of the observed vs expected length variation for loci in the validation set, stratified by the motif length of the most common LPS motif.

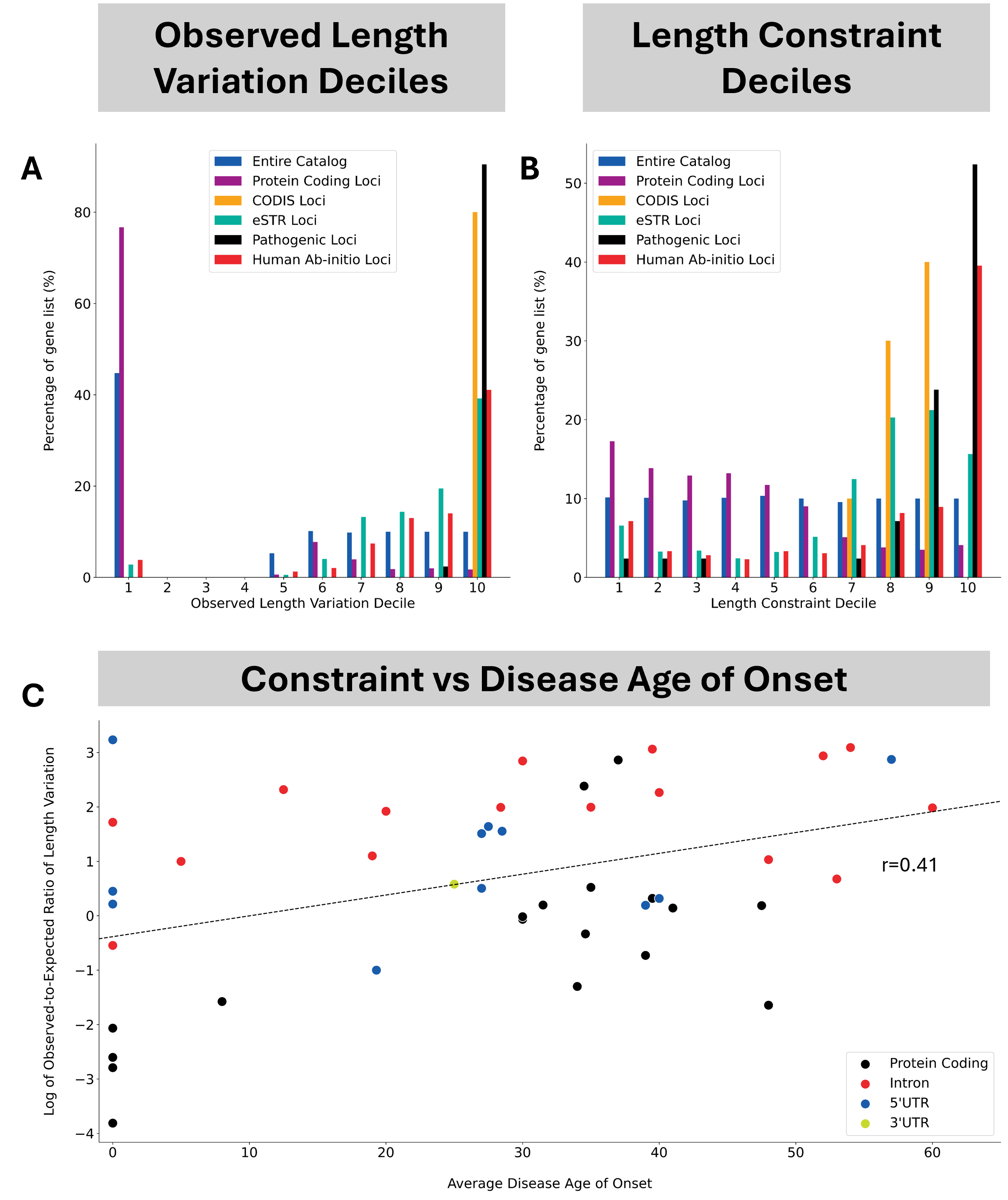

Supplementary Figure 20: Biological characteristics of TR length constraint

A) Grouped bar plot of the percentage of STRs belonging to each of the five specified sets which fall into each of the deciles of observed length variation. Observed length variation for each STR is the standard deviation of the length of the longest pure segment. B) Grouped bar plot of the percentage of STRs belonging to each of the five specified sets which fall into each of the deciles of length constraint. C) Scatterplot of the average age of onset of disease for each of 46 disease-associated STRs plotted against the logarithm of their observed-to-expected ratio of length variation (TR length constraint).

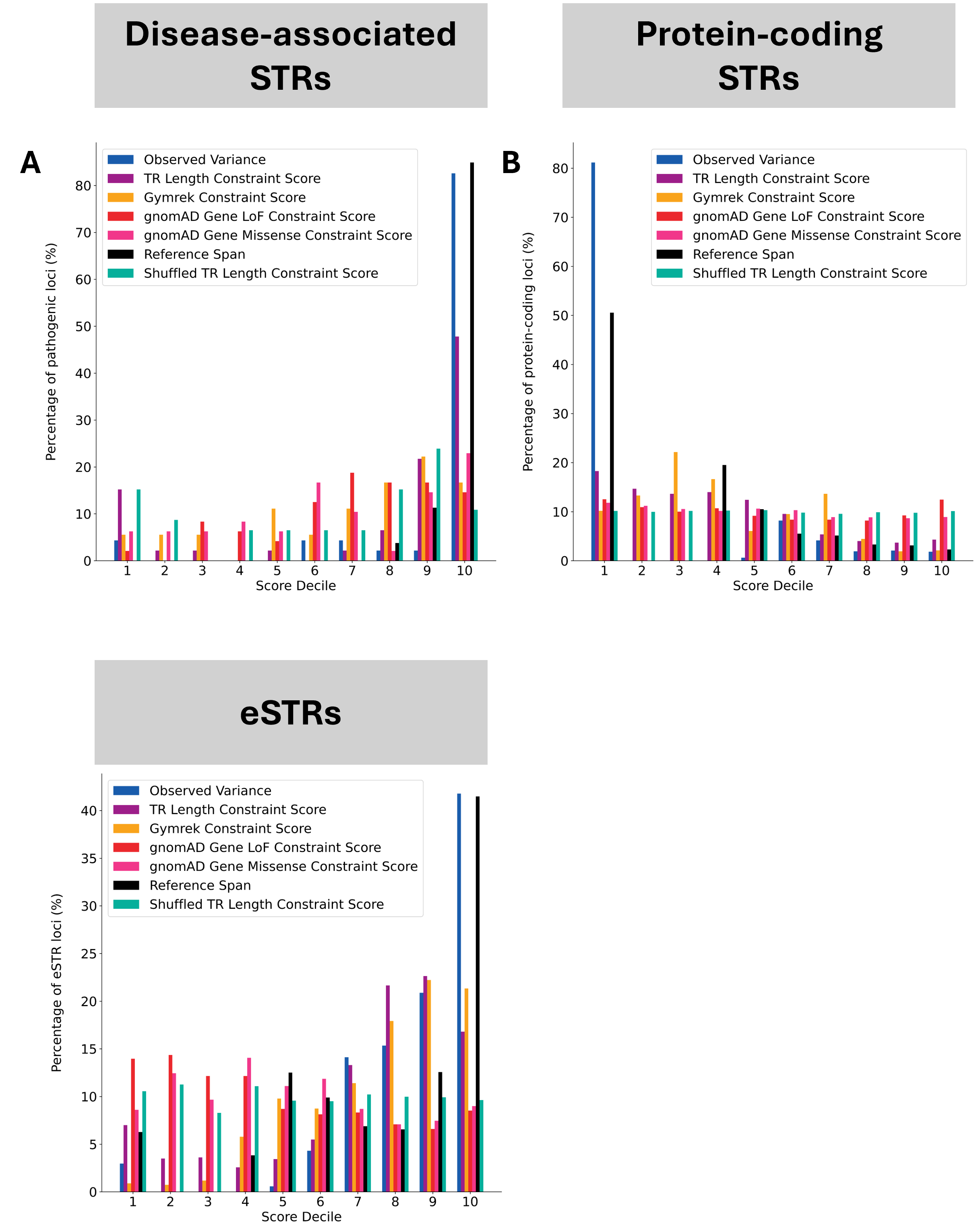

Supplementary Figure 21: Benchmark of TR Length Constraint against other approaches

A-C) Grouped bar plots of the given set of STRs belonging to each decile of values for each of the five listed methods. The sets used are as follows: A) disease-associated STRs, B) protein-coding STRs, and C) eSTRs.

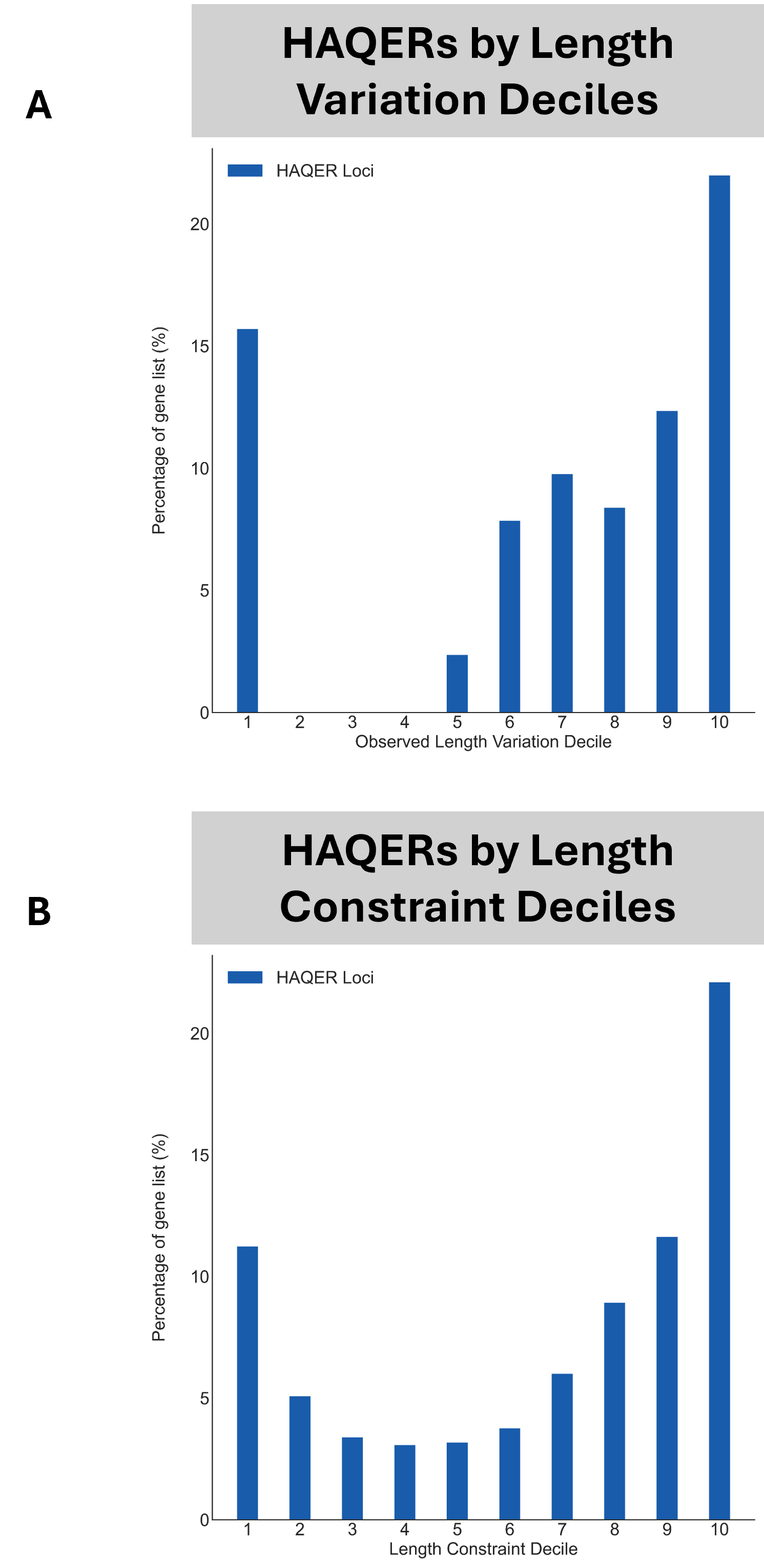
Supplementary Figure 22: LPS variation and TR Constraint scores enrich for Human Ancestor Quickly Evolved Regions

A) Barplot of deciles of LPS variation where each is measured for its overlap with the HAQER regions derived by Yoo, et al^43^. B) Barplot of deciles of TR Length Constraint where each is measured for its overlap with HAQER regions.

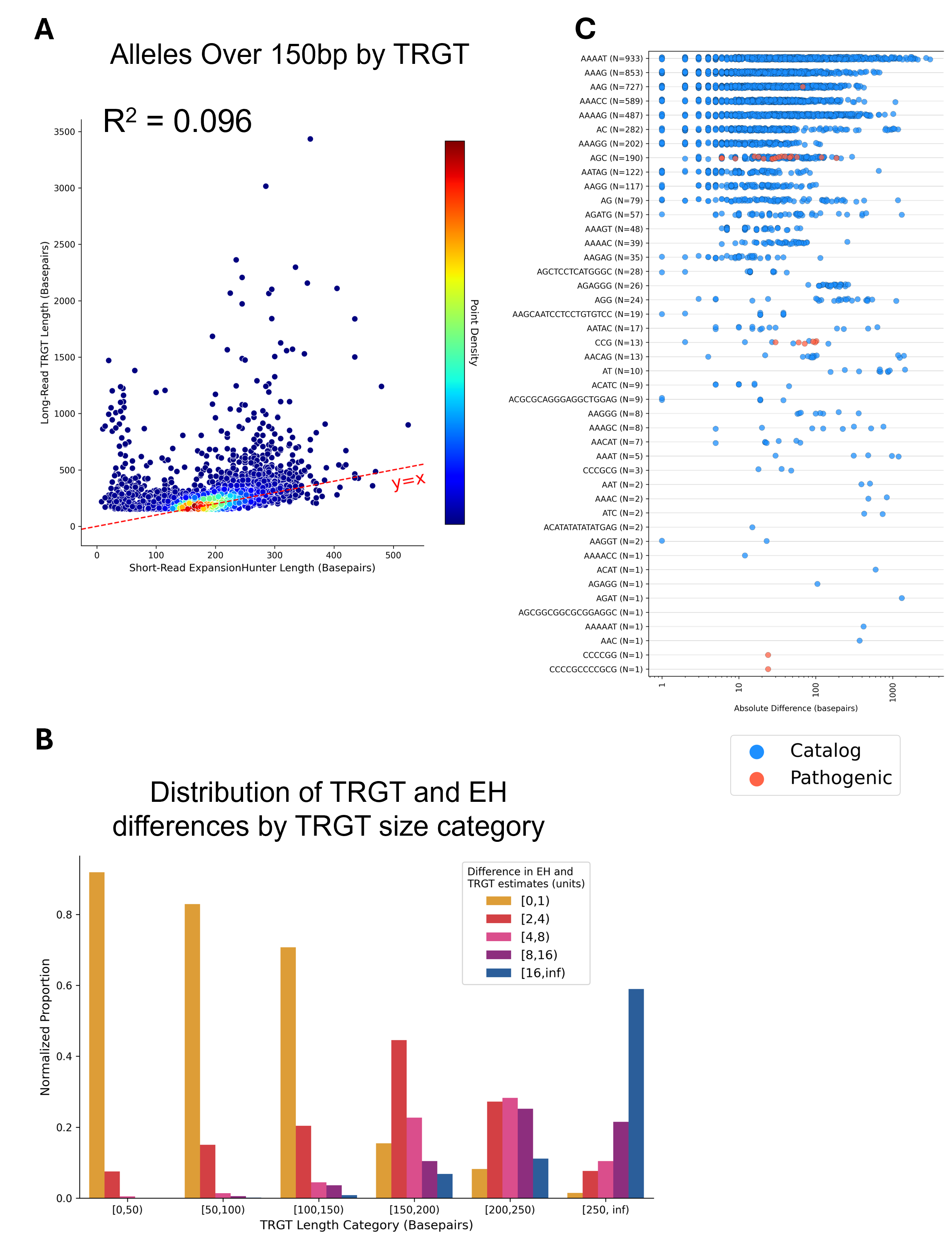

Supplementary Figure 23: Short-read vs long-read comparison using ExpansionHunter

A) Comparison of estimated length of tandem repeats from paired short-read and long-read datasets for alleles estimated by TRGT to be above 150 bp. Short-read data TR lengths estimated with ExpansionHunter (EH) and long-read data TR lengths estimated with TRGT. B) Grouped bar plot of the proportion of alleles in each category of difference in length between TRGT and EH estimates, stratified by TRGT estimate of allele length. C) Stripplot of difference in estimate of allele length by TRGT and EH for alleles estimated by TRGT to be above 150 bp, stratified by motif.

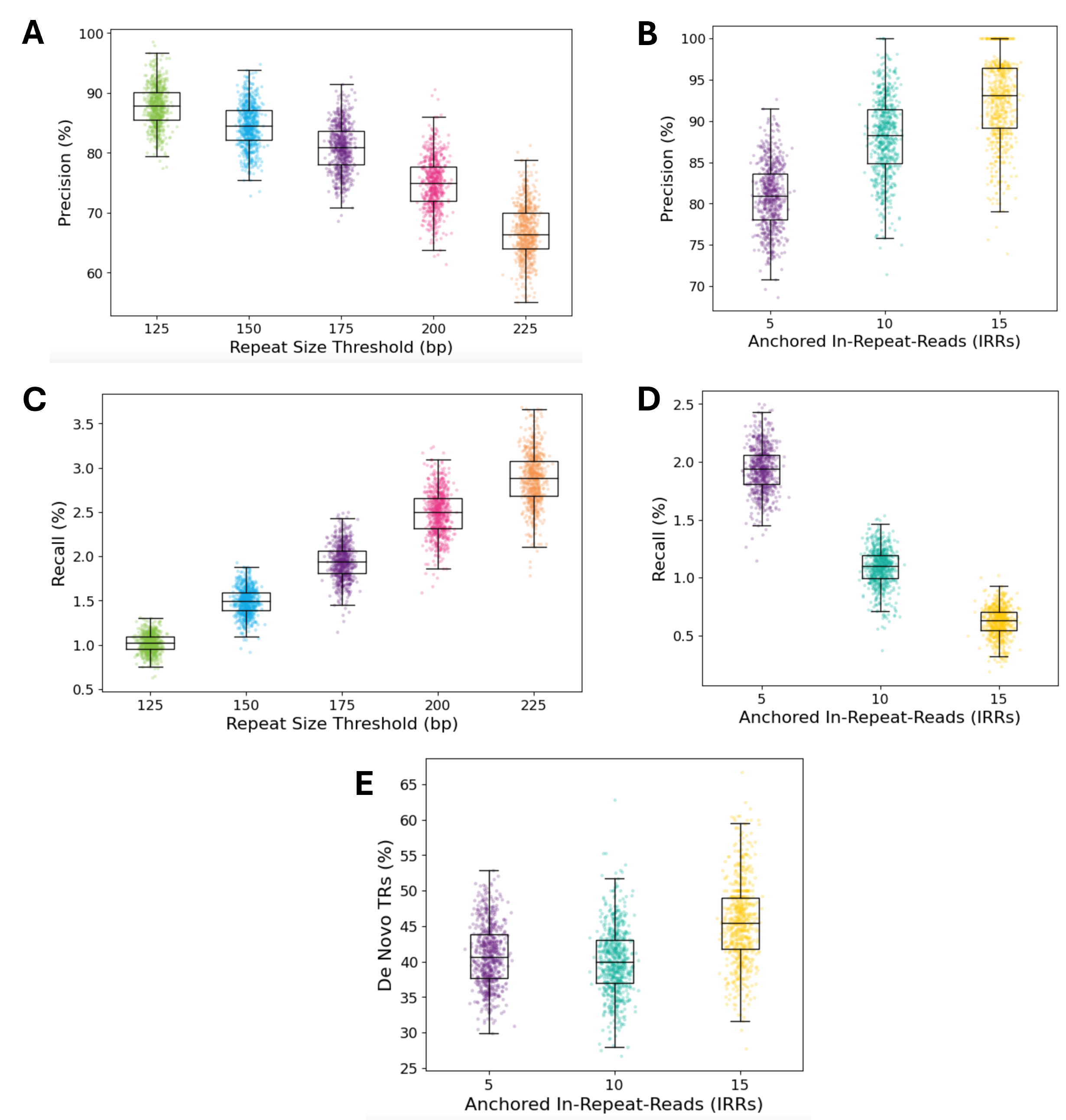

Supplementary Figure 24: Short-read approaches achieve high precision but low recall at identifying large TRs

A-B) Precision of ExpansionHunter Denovo (EHDn) in identifying large TRs. A) Precision of EHDn over a range of threshold TR sizes as determined by TRGT on the long-read data. B) Precision of EHDn at the 175 bp threshold size over a range of thresholds of anchored in-repeat-reads. C-D) Recall of EHDn in identifying large TRs. C) Recall of EHDn over a range of threshold TR sizes as determined by TRGT on the long-read data. D) Recall of EHDn at the 175 bp threshold size over a range of thresholds of anchored in-repeat-reads. E) Percentage of EHDn calls that did not correspond to any locus genotyped by TRGT (de-novo TRs) over a range of thresholds of anchored in-repeat-reads.

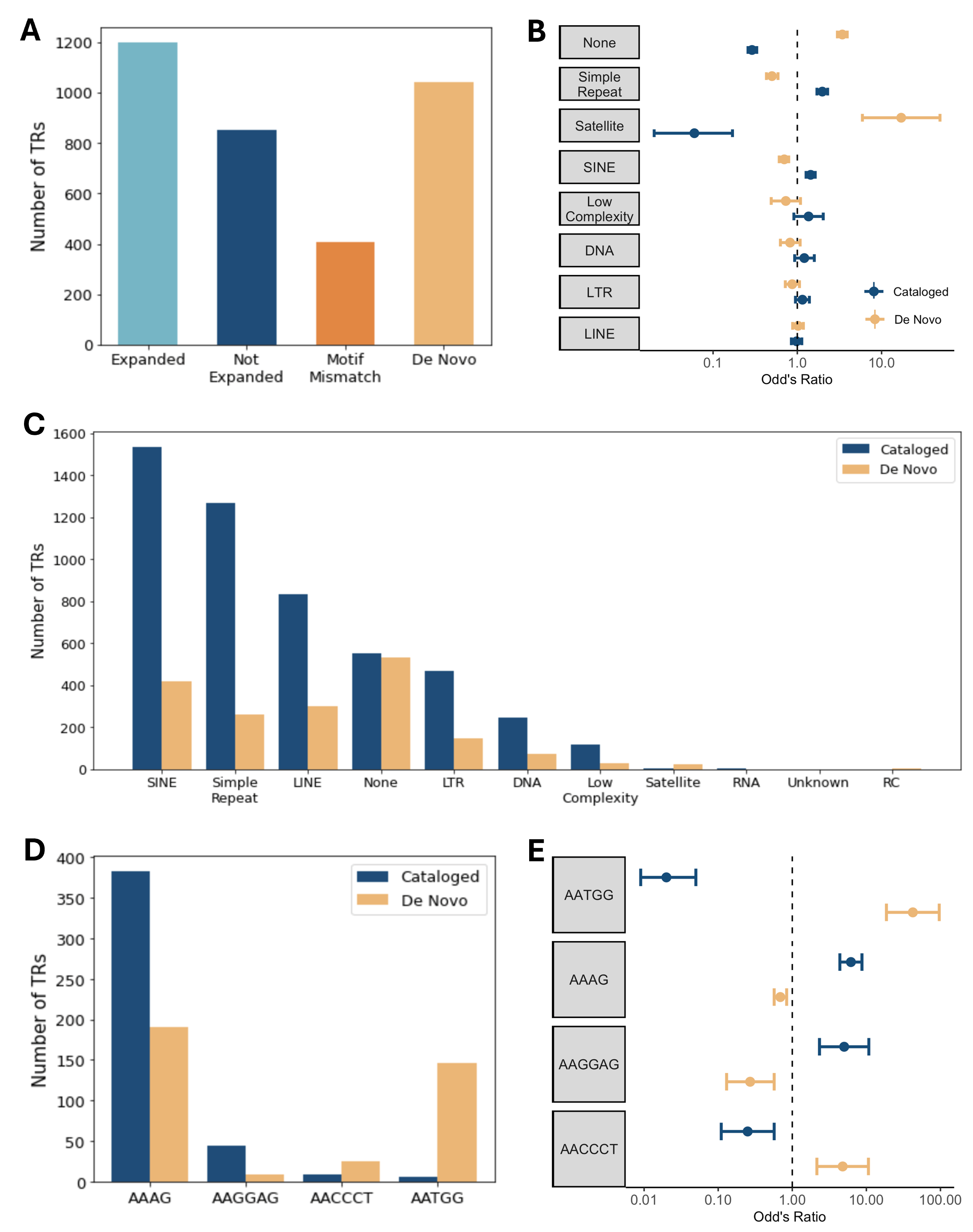

Supplementary Figure 25: Enrichment of repeat classes and repeat motifs in de novo TRs

A) Number of TRs called by EHDn categorized into groups: truly expanded as evidenced by the TRGT call (expanded), not expanded as evidenced by the TRGT call (not expanded), having an alternative motif than that called by TRGT (motif mismatch), or not genotyped by TRGT at all (de novo). B) Enrichment of different repeat classes in the cataloged and de novo TR groups. C) Frequencies of different repeat classes in the cataloged and de novo TR groups. D) Frequencies of significantly observed motifs between the cataloged and de novo TR groups. E) Odds ratios observed for the significantly different motifs between the cataloged and de novo TR groups.

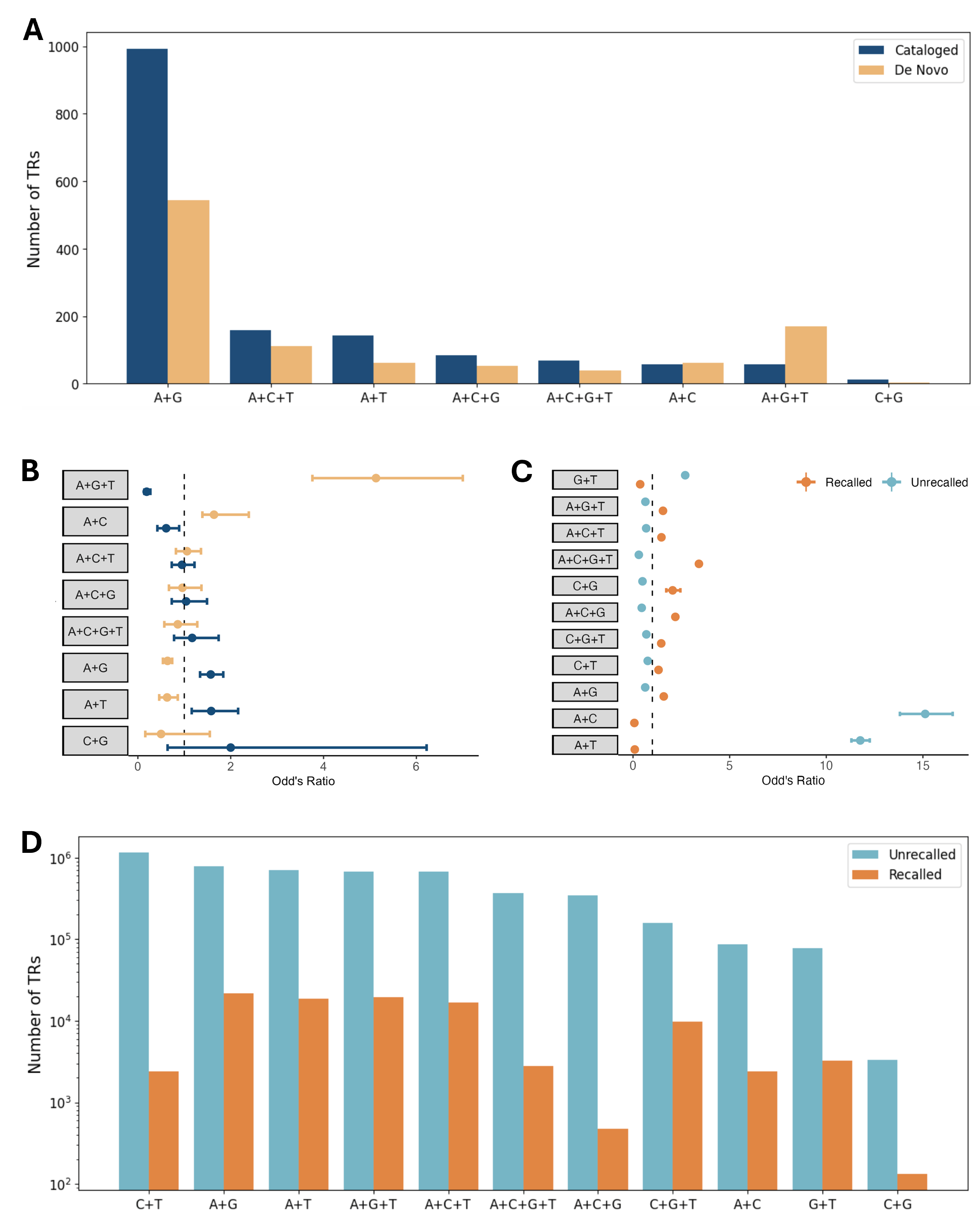

Supplementary Figure 26: Nucleotide compositions of the motifs called by EHDn

A) Frequencies of different nucleotide compositions of motifs in the cataloged and de novo TR groups. B) Odds ratios of the different nucleotide compositions of motifs in the cataloged and de novo TR groups. C) Odds ratios of the different nucleotide compositions of motifs in the recalled and unrecalled TR groups. D) Frequencies of the different nucleotide compositions of motifs in the recalled and unrecalled TR groups.

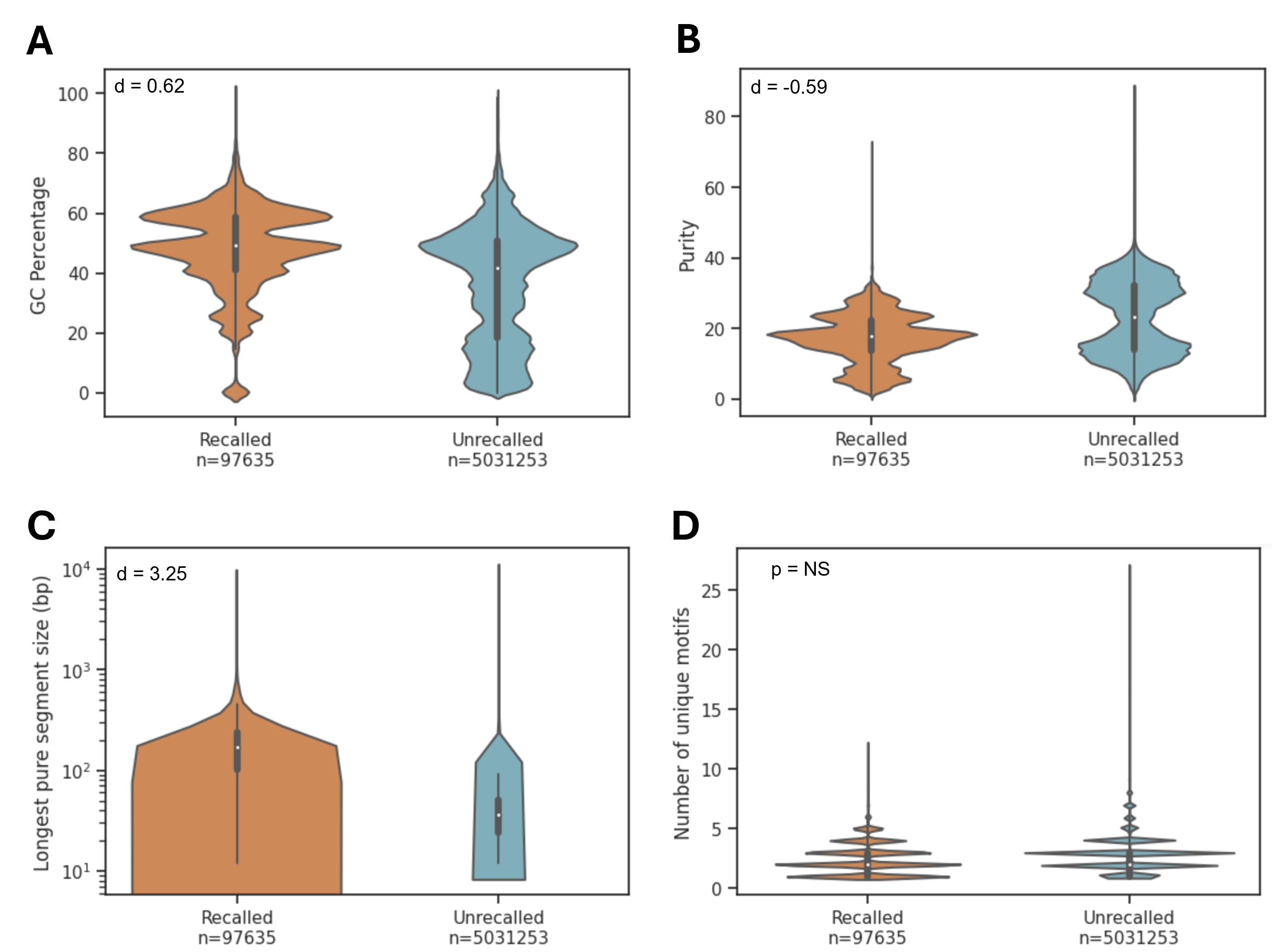

Supplementary Figure 27: Sequence characteristics of large TRs called and not called by EHDn

A-D) Violin plots of sequence characteristics of TRs determined by TRGT to be longer than 175 bp which were identified by (recalled) and not identified by (unrecalled) EHDn: A) GC percentage, B) TR purity, C) length of the longest pure segment, and D) number of unique motifs.

### **SUPPLEMENTARY TABLES**

#### **Supplementary Table 1: Linkage region ranking statistics**

| Region size (bp) | Phenotype | Gene | Motif | Rank | STRs in region | Percentile (%) |
| --- | --- | --- | --- | --- | --- | --- |
| 1,734,113 | CANVAS | RFC1 | AAAAG | 1 | 3,778 | 100 |
| 9,755,100 | DM2 | CNBP | AGGC | 1 | 14,233 | 100 |
| 2,500,000 | HD | HTT | AGC | 1 | 4,232 | 100 |
| 3,510,527 | SCA10 | ATXN10 | ATTCT | 1 | 5,591 | 100 |
| 1,481,291 | SCA27B | FGF14 | AAG | 1 | 2,139 | 100 |
| 5,392,165 | SCA3 | ATXN3 | CTG | 1 | 8,798 | 100 |
| 4,332,499 | FRDA | FXN | AAG | 2 | 4,742 | 99.98 |
| 6,009,239 | SCA31 | BEAN1 | AAAAT | 2 | 11,683 | 99.99 |
| 2,399,675 | SCA37 | DAB1 | AAAAT | 2 | 3,334 | 99.97 |
| 7,164,849 | FAME1 | SAMD12 | AAAAT | 3 | 11,130 | 99.98 |
| 63,599,999 | NIID | NOTCH2NLC | CCG | 15 | 61,228 | 99.98 |
| 20,870,206 | FAME2 | STARD7 | AAAAT | 19 | 25,124 | 99.93 |
| 11,251,529 | FTD/ALS | C9orf72 | CCCCGG | 23 | 17,497 | 99.87 |
| 7,499,999 | SCA4 | ZFHX3 | CCG | 186 | 14,334 | 98.71 |
| 9,507,126 | FAME3 | MARCHF6 | ATTTT | 296 | 13,650 | 97.84 |

For each linkage region presented in Figure 4a or Supplementary Figure 8, this table provides the exact size of the linkage region, the phenotype for which it was defined, the gene containing the causal STR, the motif of the causal STR, the rank of the causal STR within the linkage region (ranking by genome-wide PLVI), the number of STRs in the linkage region, and the percentile of the causal STR within the linkage region. Note that *CNBP* required specification of the pathogenic motif rather than simply using its most common LPS motif to achieve its rank.

### **SUPPLEMENTARY FILES**

#### **Supplementary File 1: Total allele lengths at disease-associated pathogenic loci in discovery cohort**

Lists the distribution of raw allele lengths (in bp) observed for 42 disease-associated loci across the cohort. For each locus, the ID of the TR is given, along with the name of the corresponding gene, then the allele length in the cohort for the following percentiles is given: 0, 1, 5, 10, 15, 20, 25, 30, 35, 40, 45, 50, 55, 60, 65, 70, 75, 80, 85, 90, 95, 99, 99.9, and 100. The lengths listed refer to the measurements of the genomic region specified by the TRID, which is often not identical to the typical coordinates used for these pathogenic loci. Some loci contain multiple repetitive regions (e.g., *FMR1* and *HTT*).

#### **Supplementary File 2: Longest pure segment lengths at disease-associated loci in discovery cohort**

Lists the distribution of longest pure segment lengths (in bp) observed for 42 disease-associated loci across the cohort. For each locus, the ID of the TR is given (TRID), along with the name of the corresponding gene (TRName), the motif being measured (longestPureSegmentMotif), and the number of alleles in which that motif accounted for the longest pure segment at this locus (N_motif). Each locus can appear numerous times with different motifs accounting for the longest pure segment and the sum of the values of ‘N_motif’ can reach as high as 1,086 (two alleles for each of 543 individuals), though there are missing genotype calls for many loci, so the values do not always sum to 1,086 exactly. For each row, the length of the longest pure segment (for the corresponding motif) in the cohort for the following percentiles is given: 0, 1, 5, 10, 15, 20, 25, 30, 35, 40, 45, 50, 55, 60, 65, 70, 75, 80, 85, 90, 95, 99, 99.9, and 100.

#### **Supplementary File 3: TR Constraint results for loci within 10kb of genes**

For each of the TR-Explorer catalog loci, this table lists the observed and expected (predicted) values for standard deviation of LPS length (combined across all LPS motifs). It then gives the computed observed-to-expected ratio. It also lists whether the locus was used as part of the training, validation, or testing of the model.
